# Gene flow in phylogenomics: Sequence capture resolves species limits and biogeography of Afromontane forest endemic frogs from the Cameroon Highlands

**DOI:** 10.1101/2020.10.09.332767

**Authors:** Matej Dolinay, Tadeáš Nečas, Breda M. Zimkus, Andreas Schmitz, Eric B. Fokam, Emily Moriarty Lemmon, Alan R. Lemmon, Václav Gvoždík

## Abstract

Puddle frogs of the *Phrynobatrachus steindachneri* species complex are a useful group for investigating speciation and phylogeography in Afromontane forests of the Cameroon Highlands (Cameroon Volcanic Line) in western Central Africa. The species complex is represented by six morphologically relatively cryptic mitochondrial DNA lineages, with only two of them distinguished at the species level – southern *P. jimzimkusi* and Lake Oku endemic *P. njiomock*, leaving the remaining four lineages with a pooled identification as ‘*P. steindachneri*’. In this study, the six mtDNA lineages are subjected to genomic sequence capture analyses to delimit species (together with morphology) and to study biogeography. Nuclear DNA data (387 loci; 571,936 aligned base pairs) distinguished all six mtDNA lineages, but the splitting pattern and depths of divergences supported only four main clades—besides *P. jimzimkusi* and *P. njiomock*, only two from the four ‘*P. steindachneri*’ mtDNA lineages. One is here described as a new species, *P. sp. nov*. Reticulate evolution (hybridization) was detected within the species complex with morphologically intermediate hybrid individuals placed between the parental species in phylogenomic analyses, forming a phylogenetic artefact – a ladder-like tree pattern. The presence of hybrids is undesirable in standard phylogenetic analyses, but is essential and beneficial in the network multispecies coalescent. This latter approach allowed us an insight into the reticulate evolutionary history of these endemic frogs. Introgressions likely occurred during the Middle and Late Pleistocene climatic oscillations, due to the cyclic connections (likely dominating during cold glacials) and separations (warm interglacials) of montane forests. The genomic phylogeographic pattern supports the earliest division between southern (Mt. Manengouba to Mt. Oku) and northern mountains at the onset of the Pleistocene. Further subdivisions occurred in the Early Pleistocene separating populations from the northernmost (Tchabal Mbabo, Gotel Mts.) and middle mountains (Mt. Mbam, Mt. Oku, Mambilla Plateau), as well as the microendemic lineage restricted to Lake Oku (Mt. Oku). Mount Oku harboring three species is of particular conservation importance. This unique model system is highly threatened as all the species within the complex have exhibited severe population declines in the past decade, placing them on the brink of extinction. We therefore urge for conservation actions in the Cameroon Highlands to preserve their diversity before it is too late.

## 1. Introduction

One of the most important evolutionary processes in the systematics of living organisms is the diversification of one population into more lineages, the process leading to a formation of new species. Phylogenomics (i.e. genome-wide genetic markers applied in phylogenetics) is now widely used as an important approach to reconstruct phylogenies. However, several evolutionary processes can cause discordant phylogenetic inference (Degnan, 2018). Gene flow between species or hybridization (interspecific introgression, reticulate evolution) is one of possible sources of gene-trees incongruences, gaining increased attention in the genomic era (Mason et al., 2019; Ebersbach et al., 2020). Hybridization has been proven to be quite common despite the fact that partial reproductive barriers have already been established (Good et al., 2008; Li et al., 2016; Dylite et al., 2019; Pyron et al., 2020). Gene flow occurring across species boundaries can have a significant impact on the phylogenetic inference (Leaché et al., 2014; Mallet et al., 2016), especially in taxa with rapid radiation history (Winker et al., 2018; Irisarri et al., 2018). Leaché et al. (2014) tested the influence of gene flow on speciestree estimations and found that gene flow led to overestimation of population sizes and underestimation of species divergence times. Another commonly used approach to construct species phylogenies is based on data concatenation. A supermatrix is created from multiple loci combined together and analyzed as if they represent only one locus (McVay and Carstens, 2013). The phylogenetic inference based on concatenation can also be difficult when gene flow is present due to a distortion of topology (Eckert and Carstens, 2008; Zarza et al., 2016). Therefore, careful sampling design and assignments of samples to “species” (for species-tree estimations) are necessary in genomic phylogeographic and species delimitation studies (Leaché et al., 2014). Further, both theoretical and empirical studies are needed to gain more information about the influence of gene flow on the phylogenomic inference. Here, we add an empirical sequence-capture phylogeographic study (i.e., with shallow divergences) to demonstrate that gene flow causes a ladder-like tree pattern in the concatenation and coalescent-based species-tree analyses.

We used phrynobatrachid frogs (Phrynobatrachidae) from Afromontane forests in western Central Africa as a model system. Montane forests represent so called ‘sky islands’ (Heald, 1951) - habitats temporarily connected and disconnected depending on topography of mountains and climatic oscillations. Sky islands are, similarly to oceanic islands, an ideal environment for studies of evolutionary processes (Brown, 1978). These isolated, high-elevation habitats are geographically distributed among different mountain ranges, and patterns of their distributions act like a trigger for diversification induced by ecological and microevolutionary factors (McCormack et al., 2009). Organisms living on sky islands have limited dispersal opportunities due to isolation by lowland habitats in areas separating different mountains. Populations separated by unfavorable environmental conditions are more prone to undergo speciation, if separations persist for longer periods. During climatic fluctuations, habitat connections can be re-established between and among these areas. This allows gene flow between populations that have accumulated genetic differences, although reproductive isolation remains incomplete. Typically, the Pleistocene climatic cycles modulated levels of gene flow and shaped genetic structure of organisms living on these sky islands (Carstens and Knowles, 2007; Leaché et al., 2013; Favé et al., 2015). However, most previous studies focused on organisms from sky islands in the temperate climatic zone, and relatively little is known about the speciation processes on tropical sky islands, especially in Africa.

The mainland highlands of the Cameroon Volcanic Line (CVL; Burke, 2001) are an example of Afrotropical sky islands, including the ecoregion known as the Cameroon Highlands forests (Burgess et al., 2004; Dinerstein et al., 2017) lying on the biogeographical border between Central (Cameroon) and West Africa (Nigeria). The Cameroon Highlands forests extend from Mount Manengouba in the southwest to the Gotel Mountains and Tchabal Mbabo in the northeast, and include forests on the Obudu and Mambilla Plateaus in Nigeria and some more northern and eastern peripheral mountains (Fig. 1). Mount Oku (3011 m a.s.l.), with its crater lake Lake Oku (at 2227 m a.s.l.), is the highest peak within the region, while the highest mountain of the CVL, Mount Cameroon, is considered a separate ecoregion (Burgess et al., 2004; Dinerstein et al., 2017). Geologically, the Cameroon Highlands were uplifted mostly during the Miocene, approximately 21–7 Mya, with the oldest being the Oku Massif, which originated 31–22 Mya, and the youngest Mt. Manengouba originating only 1 Mya (Marzoli et al., 2000). Bioclimatically, there is an increasing northeastern gradient of aridity at lower elevations with the southernmost Mt. Manengouba surrounded by lowland coastal rainforests while more northern mountains by the Guineo-Congolian forest-savanna mosaic ecoregions (Burgess et al., 2004; Dinerstein et al., 2017). Only limited phylogeographic research has been conducted about the montane forests of the Cameroon Highlands, both in terms of number of studies, or number of sampling sites and genetic markers (Smith et al., 2000; Kadu et al., 2011; Missoup et al., 2012, 2016; Zimkus and Gvoždík, 2013; Taylor et al., 2014). This study aims to broaden the knowledge of the phylogeography of the Cameroon Highlands forests.

**Fig. 1.**
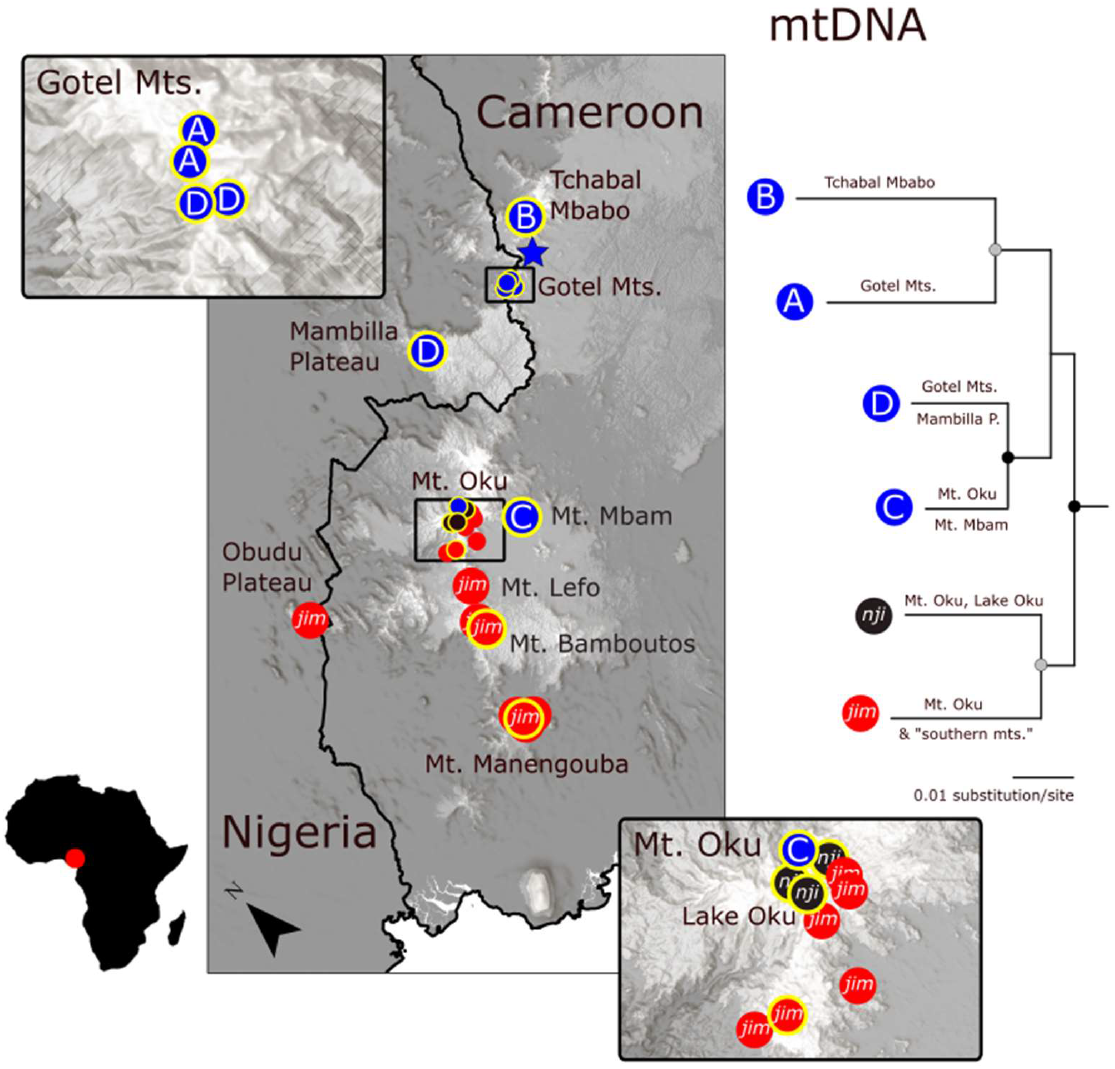
Puddle frogs of the *Phrynobatrachus steindachneri* complex as a model for a speciation study and the phylogeography of montane forests of the Cameroon Highlands (a part of the Cameroon Volcanic Line), western Central Africa. The species complex is represented by six morphologically relatively cryptic mitochondrial DNA lineages (Zimkus & Gvoždík, 2013), with two of them distinguished at species level – southern *P. jimzimkusi* (*jim* in red) and Lake Oku endemic *P. njiomock* (*nji* in black), and leaving the remaining four under the name ‘*P. steindachneri*’ (A, B, C, D in blue). Maps show sampling localities with insets detailing contact zones within the Oku Massif and Gotel Mountains. Yellow-bordered circles denote localities represented in the sequence-capture genomic analyses. Blue star marks type locality of *P. steindachneri*. The maximum likelihood mtDNA phylogenetic tree is adapted from Gvoždík et al. (2020) with black and gray dots meaning high and intermediate supports, respectively.

Puddle frogs (*Phrynobatrachus*) are mostly small to minute (body size typically < 40 mm), generally ground-dwelling anurans with aquatic larvae. They represent one of the most diverse sub-Saharan anuran lineages (Zimkus et al., 2010), with 95 species presently recognized (Frost, 2020). One clade is primarily distributed in and around the Cameroon Highlands (Zimkus et al., 2010), and thus is referred to as the Cameroon radiation clade (Gvoždík et al., 2020). Within this radiation exists a monophyletic species group (*P. steindachneri*) containing one submontane species *P. cricogaster* Perret, 1957 and a complex of closely related species strictly confined to the proximity of water bodies in montane forests, including small stream-side fringing gallery forests. Frogs within the *P. steindachneri* species complex are primarily ground dwelling, but also spend their time in and around water (typically in calm, pool-like passages of streams, where they lay eggs) and climb lower vegetation (up to 1 m high, usually for rest). However, little is known about their ecology, and the taxonomy of this group has been long debated (Amiet, 1971; Gartshore, 1986; Zimkus and Gvoždík, 2013; Amiet and Goutte, 2017). This species complex is known from the elevation range 1300 to 2800 m a.s.l. (Doherty-Bone and Gvoždík, 2017), and contains six morphologically relatively cryptic evolutionary lineages as based on the mitochondrial DNA (Zimkus and Gvoždík, 2013; Gvoždík et al., 2020). Two of the six lineages were distinguished at species level: *P. jimzimkusi* Zimkus, Gvoždík, Gonwouo, 2013 from the southern mountains (Mt. Manengouba to Mt. Oku, including Mt. Bamboutos, Mt. Lefo, and Obudu Plateau in Nigeria), and *P. njiomock* Zimkus, Gvoždík, 2013 endemic to Lake Oku and nearby habitats (Zimkus and Gvoždík, 2013). The remaining four lineages from northern mountains are currently identified as *P. steindachneri* Nieden, 1910 (Zimkus and Gvoždík, 2013), and to distinguish them they have been assigned capital letters as ‘*P. steindachneri*’: ‘A’ from the Gotel Mts., ‘B’ from Tchabal Mbabo, ‘C’ from Mt. Mbam and Mt. Oku, and ‘D’ from the Mambilla Plateau and Gotel Mts. (Gvoždík et al., 2020, supplemental information). Individuals having mitochondrial DNA (mtDNA) haplotypes of the presumed Lake Oku endemic, *P. njiomock*, were found as far as 8 km from Lake Oku in a forest on the north-eastern slope of Mt. Oku (Zimkus and Gvoždík, 2013). This opens a question whether such individuals are ‘true’ *P. njiomock* or rather hybrids. And as Mt. Oku and Gotel Mts. harbor three and two lineages, respectively—though none were found syntopically—these regions represent potential areas for inter-lineage gene flow, i.e. introgression/hybridization. For the present knowledge on the phylogenetic relationships (mtDNA-based) and distributions of all the six lineages see Fig. 1.

In this study, using a combination of mtDNA, genomic data acquired by the sequencecapture method anchored hybrid enrichment (Lemmon et al., 2012), and morphology, we aim to (1) evaluate if gene flow/hybridization is present and its effect on phylogenetic reconstruction; (2) delimit species within the six mtDNA lineages, accompanied by morphological investigation; (3) interpret phylogeographic patterns in the context of historical biogeography of montane forests of the Cameroon Highlands.

## 2. Material and methods

### 2.1. Genetics

#### 2.1.1. Sampling

Altogether, we compiled molecular data from 115 individuals of the *P. steindachneri* species complex from the whole known range and three outgroups (Fig. 1, Table S1). Submontane *P. cricogaster*, the sister lineage to the *P. steindachneri* complex, was included as the nearest outgroup, while *P. mbabo* and *P. batesii* were representative of two more distant outgroups from within the Cameroon radiation (Gvoždík et al., 2020). Mitochondrial DNA data were available for all these samples, while nuclear DNA (nDNA) sequence capture genomic data were generated for a subset of 20 samples plus two outgroups (*P. cricogaster, P. batesii*). According to mtDNA, the following sample numbers were analyzed within the ingroup (lineage abbreviation/mtDNA/nDNA): *P. jimzimkusi* (*jim*/62/3), *P. njiomock* (*nji*/23/3), ‘*P. steindachneri*’ A (A/5/3), ‘*P. steindachneri*’ B (B/3/3), ‘*P. steindachneri*’ C (C/14/4), and ‘*P. steindachneri*’ D (D/8/4). The sampling represented in the genomic approach comprised *P. jimzimkusi* from three separate mountain massifs, and *P. njiomock* from the core distribution area around Lake Oku and a site distant from the lake. Furthermore, specimens from multiple sympatric lineages from Mt. Oku (*jim*: 1, *nji*: 3, C: 2) and Gotel Mts. (A: 3, D: 3) were included to identify if some might represent hybrids. Throughout this study, we ordered ingroup lineages from the north to the south.

#### 2.1.2. mtDNA – laboratory and alignment

Within the ingroup, 64 mtDNA sequences were included from GenBank [originating primarily from Zimkus (2009), and Zimkus and Gvoždík (2013); Table S1], and 51 samples were newly sequenced (deposited in GenBank xxxxxxxx–xxxxxxxx). We amplified a portion of the 16S rRNA gene *(16S*) using the primers 16SL1 and 16SH1 adapted from Palumbi et al. (1991) by a previously described PCR protocol (Gvoždík et al., 2020). A shorter fragment was generated from sequencing in a single new sample. After low-quality beginnings of sequences were trimmed and the sequences were aligned using MAFFT v7 (Katoh and Standley, 2013) with default setting as incorporated in Geneious R8.1 (Biomatters Ltd.; Kearse et al., 2012), the resulting alignment had 512 base pairs (bp) for the ingroup (and also with the nearest outgroup, *P. cricogaster*), and 519 bp with all three outgroups.

#### 2.1.3. Genomic sequence capture procedure

For 22 samples, including two outgroups, genomic sequence capture data were generated using the anchored hybrid enrichment (AHE) methodology described by Lemmon et al. (2012). At the Florida State University’s Center for Anchored Phylogenomics (www.anchoredphylogeny.com), Illumina libraries were prepared from DNA extracts following the protocol of Meyer and Kircher (2010), with modifications outlined in Prum et al. (2015). After adding 8bp adapter indexes, libraries were pooled in equal concentrations, then enriched using the Amphibian AHE probe design described in Hime et al. (2020). Enriched libraries were sequenced on one Illumina HiSeq2500 lane with a PE150bp protocol in the Translational Lab at the FSU College of Medicine. The total sequence data collected was 19.5 Gb.

After demultiplexing the Illumina reads, adapters were removed, and sequencing errors were corrected while overlapping read pairs were merged as outlined in Rokyta et al. (2012). Reads were assembled into clusters using a quasi-denovo assembly approach described by Hamilton et al. (2016), using the *Pseudacris nigrita* (Hylidae) and *Gastrophryne carolinensis* (Microhylidae) sequences from the Amphibian AHE probe design as divergent references. Consensus sequences were constructed from assembly clusters containing at least 117 reads, a threshold which allowed the removal of low-copy paralogs and any potential low-level contamination, while still retaining the large majority of target loci. After homologous consensus sequences were gathered across the 22 individuals, orthology was assessed using a neighbor-joining approach that utilized an alignment-free pairwise distance matrix (methodological details provided in Hamilton et al., 2016). Orthologous sets of sequences were then aligned using MAFFT (v7.023b, Katoh and Standley, 2013). Raw alignments were subjected to an automated trimmer/masker that identifies misaligned regions, as described in Hamilton et al. (2016; but with parameters: MINGOODSITES=16, MINPROPSAME=0.3). In order to mitigate the possible effects of missing data (Lemmon et al., 2009), we removed sites represented by fewer than 9 of the 22 taxa. The alignments were inspected by eye in Geneious R9 (Biomatters Ltd.; Kearse et al., 2012) to ensure the automated trimmer/masker settings were suitable.

The probes used for the hybrid enrichment successfully captured 387 loci. Altogether, 344 of the alignments contained all 22 samples, while 43 alignments had one or more samples missing. The full concatenated dataset comprised 571,936 aligned bp.

#### 2.1.4. Maximum-likelihood trees & genetic distances: mtDNA and concatenated genomic data

Maximum-likelihood (ML) trees based on the mtDNA (*16S*) and concatenated genomic datasets were generated by RAxML v8 (Stamatakis, 2014), using the GTR model with gamma-distributed rate heterogeneity [without a combination with the proportion of invariable sites following the author’s recommendation (Stamatakis, 2016)]. The topology supports were assessed by 100 rapid bootstrap inferences (Stamatakis et al., 2008). The genomic data were analyzed in two ways, with and without detected hybrid individuals (see below).

Genetic distances among and within the six main lineages as defined by mtDNA (Zimkus and Gvoždík, 2013; Gvoždík et al., 2020) were calculated as means using uncorrected p-distances for *16S* and concatenated genomic data after exclusion of hybrids in MEGA X (Kumar et al., 2018). All positions containing gaps and missing data were eliminated by pairwise and complete deletions, the latter producing 508 bp (16S) and 424,500 bp (genomic dataset), respectively.

#### 2.1.5. Detection of hybridization

Based on theoretical assumptions (sympatry of multiple lineages on Mt. Oku and in the Gotel Mts.) and phylogenetic positions of some individuals (genomic ML tree with all samples; Fig. 2), we identified whether hybrid specimens were present in our model system. We began by calling single nucleotide polymorphisms (SNPs) in the concatenated genomic alignment. SNP sites were identified as those with at least 5% minor allele frequency flanked by at least five sites on each side with less than a 5% minor allele frequency. SNPs were not retained unless at least 80% of the individuals were represented at the SNP and flanking sites. The script used to extract the SNPs is available as a supplemental file. We analyzed the SNPs in GenoDive v2.0b27 (Meirmans and Van Tienderen, 2004; Meirmans, 2013) to calculate maximumlikelihood estimations of hybrid indices (Buerkle, 2005). We tested putative hybrid individuals against a set of combinations of putative parental populations/species (i.e. reference and alternative population) based on phylogenetic and biogeographic rationales (Table 1). In addition, phasing to gametic haplotypes was performed in an experimental subset of 20 loci using the coalescent-based Bayesian algorithm of Phase v2.1 (Stephens et al., 2001; Stephens and Scheet, 2005) to check if gametic haplotypes of putative hybrid individuals segregate in different parental lineages. Input/output files were processed in SeqPHASE (Flot, 2010). Phasing was conducted under the parent-independent mutation model with a burn-in-period of 100 iterations, which was followed by 100 iterations. Results with a probability >0.70 were accepted, and RAxML trees were computed for each phased locus with the same settings as described above.

**Table 1.** Testing putative hybrids against potential parental populations by calculations of maximum-likelihood estimations of hybrid indices (HI). HI values closer to 1 indicate affinity to the reference population, values closer to 0 indicate affinity to the alternative population. The dominant parental population in bold. HI lower/upper stand for the lower and upper limits of the 95% confidence interval.

| Massif / Putative hybrid | Catalogue No. | Rationale | Reference population | Alternative population | SNPs | Hybrid index (HI) | HI lower | HI upper | Ln(likelihood) |
| --- | --- | --- | --- | --- | --- | --- | --- | --- | --- |
| Gotel Mts. / D<sub>Gotel</sub> | NMP-P6V 74521/1 | phylogenetic | A+B | **C+D<sub>Mambilla</sub>** | 3480 | 0.37 | 0.35 | 0.40 | -2456.0 |
| Gotel Mts. / D<sub>Gotel</sub> | NMP-P6V 74519/5 |  | **A+B** | C+D<sub>Mambilla</sub> |  | 0.59 | 0.57 | 0.62 | -2503.8 |
| Gotel Mts. / D<sub>Gotel</sub> | NMP-P6V 74521/2 |  | **A+B** | C+D<sub>Mambilla</sub> |  | 0.63 | 0.60 | 0.65 | -2354.9 |
| Gotel Mts. / D<sub>Gotel</sub> | NMP-P6V 74521/1 | phylogenetic–<br>biogeographic | **A** | B | 2724 | 0.78 | 0.76 | 0.81 | -2142.9 |
| Gotel Mts. / D<sub>Gotel</sub> | NMP-P6V 74519/5 |  | **A** | B |  | 0.77 | 0.74 | 0.80 | -1789.3 |
| Gotel Mts. / D<sub>Gotel</sub> | NMP-P6V 74521/2 |  | **A** | B |  | 0.77 | 0.74 | 0.80 | -1663.2 |
| Gotel Mts. / D<sub>Gotel</sub> | NMP-P6V 74521/1 | biogeographic | A | **D<sub>Mambilla</sub>** | 4714 | 0.46 | 0.44 | 0.48 | -2735.7 |
| Gotel Mts. / D<sub>Gotel</sub> | NMP-P6V 74519/5 |  | **A** | D<sub>Mambilla</sub> |  | 0.61 | 0.59 | 0.62 | -2852.3 |
| Gotel Mts. / D<sub>Gotel</sub> | NMP-P6V 74521/2 |  | **A** | D<sub>Mambilla</sub> |  | 0.64 | 0.63 | 0.66 | -2686.2 |
| Gotel Mts. / D<sub>Gotel</sub> | NMP-P6V 74521/1 | comparison –<br>testing D<sub>Gotel</sub><br>together with<br>D<sub>Mambilla</sub> | A+B | **C** | 3480 | 0.46 | 0.43 | 0.48 | -2388.1 |
| Gotel Mts. / D<sub>Gotel</sub> | NMP-P6V 74519/5 |  | **A+B** | C |  | 0.65 | 0.63 | 0.67 | -2337.9 |
| Gotel Mts. / D<sub>Gotel</sub> | NMP-P6V 74521/2 |  | **A+B** | C |  | 0.68 | 0.65 | 0.70 | -2228.9 |
| Mambilla P. / D<sub>Mambilla</sub> | MCZ A-139608 |  | A+B | **C** |  | 0.23 | 0.21 | 0.25 | -2141.4 |
| Mt. Oku / *nji*<sub>Forest</sub> | MCZ A-138119 | phylogenetic | *nji*<sub>Lake</sub> | **jim** | 2741 | 0.41 | 0.39 | 0.44 | -2655.2 |
| Mt. Oku / *nji*<sub>Forest</sub> | MCZ A-138119 | phylo–biogeo | *nji*<sub>Lake</sub> | C+D<sub>Mambilla</sub> | 3891 | 0.64 | 0.63 | 0.66 | -3445.8 |
| Mt. Oku / *nji*<sub>Forest</sub> | MCZ A-138119 | biogeographic | *nji*<sub>Lake</sub> | C<sub>Oku</sub> | 3272 | 0.64 | 0.63 | 0.66 | -3305.2 |

**Fig. 2.**
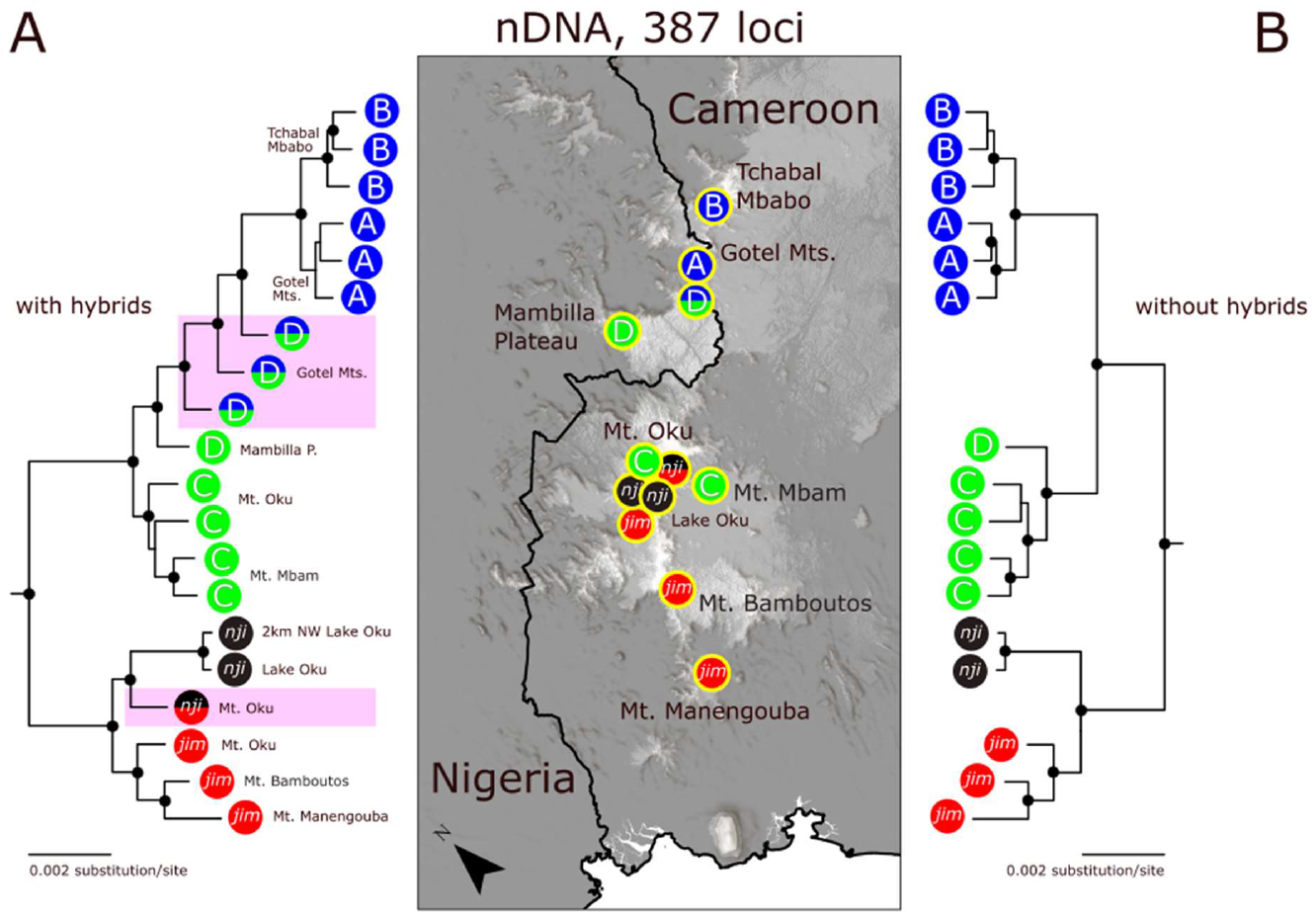
Maximum likelihood trees (RAxML) based on concatenated data from 387 sequence-capture loci (571,936 aligned bp), (A) including all specimens on left, and (B) without detected hybrids on right. Hybrids are highlighted by purple rectangles and double-colored circles. Note the ladder-like tree pattern when hybrids are included. Most of the nodes received high supports (black dots). For other explanations see Fig. 1. Outgroups (*P. cricogaster, P. batesii* not shown).

#### 2.1.6. Coalescent-based species tree and species network

We applied three coalescent-based approaches to infer phylogenomic relationships among the main evolutionary lineages, and among populations from different mountain massifs (see 2.1.7. *Phylogeography* for the latter). First, a species-tree inference based on gene trees was applied using ASTRAL-III (Zhang et al., 2018b). Unrooted gene trees were estimated from 387 loci by RAxML v8 (Stamatakis, 2014) as specified above. Two datasets were processed – without and with hybrid individuals to explore how the ASTRAL-III species tree changes when hybrids are included in a dataset. The inferred ML trees were used as input files for the ASTRAL-III analyses. The “species” were set according to mtDNA identities, comprising six tree terminals (plus two outgroups) that corresponded to the main mitochondrial lineages, and including the hybrid individuals in the analysis with hybrids. In addition, ASTRAL-III analyses were run with each individual treated as a “species”, again without and with hybrids. Branch lengths in coalescent units and node supports from gene tree quartet frequencies (i.e. local posterior probabilities) were computed as described by Sayyari and Mirarab (2016).

Further, we applied two approaches within the Bayesian framework using the BEAST 2 platform (Bouckaert et al., 2019). Due to the computational intensity of the coalescent-based approaches using sequence data, the naive binning method (Bayzid and Warnow, 2013) was applied to create 20 bins, i.e. “supergenes” from the all 387 loci with the average length of 28,597 aligned bp (23,384–31,219 bp). We followed the results of Streicher et al. (2018), who received better results with the naive binning than statistical binning (Mirarab et al., 2014), and when a limited number of naive bins was used (up to 40 in their case). We used only the nearest outgroup, *P. cricogaster*, in the Bayesian coalescent-based inferences. One applied Bayesian approach is described hereafter, and the other in the section 2.1.7.

The second approach was to estimate the reticulate phylogeny. We employed a Bayesian analysis of species networks using the birth-hybridization process prior as implemented in SpeciesNetwork (Zhang et al., 2018a). The multispecies network coalescent model allows the inclusion of reticulation branches apart from the bifurcating tree that can be interpreted as hybridization events (Degnan, 2018). Similar to the ASTRAL-III analysis, the ingroup “species” were set to the six groups corresponding to the main mitochondrial lineages with the hybrid individuals included. Following the settings of a Bayesian species-tree analysis running in parallel (StarBEAST2; see below), the species-network analysis was performed with the HKY site model, strict clock, and clock rate of 0.001375 as inferred in the Bayesian species-tree analysis (see below). The network analysis was run for 500 million generations, saving each 5000th generation, and discarded the first 10% as burn-in after an inspection in Tracer v1.7 (Rambaut et al., 2018), where it was checked that the effective sample size (ESS) was satisfactorily high for all parameters. To check for stability of parameter estimations as a sign of convergence, we ran the analysis twice using different starting seeds. Resulting species networks were visualized and inspected in IcyTree (Vaughan, 2017).

#### 2.1.7. Phylo geo graphy

The third coalescent-based approach allowed the inference of phylogenetic relationships among populations from different mountain massifs. A Bayesian species-tree approach using the multispecies coalescent model as implemented in StarBEAST2 (Ogilvie et al., 2017) was applied on the binned genomic dataset without hybrid individuals. Populations of the montane *P. steindachneri* species complex from different mountain massifs (per species) were treated as “species” producing nine tree terminals, with the addition of the sister species of the complex (submontane *P. cricogaster*). Several preliminary inspection analyses with different settings were run as recommended by Drummond and Bouckaert (2015), and these showed problems reaching convergence and sufficient ESS values when a relaxed clock (uncorrelated log-normal) and too complex site models were applied on this large genomic dataset. Therefore, to avoid overparametrization (Barido-Sottani et al., 2018) a strict clock and the HKY site model unlinked among the 20 “supergenes” (bins) were applied, together with the birth-death model as the tree prior unlinked among the 20 bins and analytical integration of population sizes. Time calibration priors were set for the splits between the *P. steindachneri* complex and *P. cricogaster*, and *P. jimzimkusi* and *P. njiomock*, respectively, following Gvoždík et al. (2020). Normal distributions were used with the means and standard deviations of 3.78 ± 0.7 Mya and 1.43 ± 0.4, which gave central 95% ranges about 2.4–5.2 Mya and 0.6–2.2 Mya, respectively. The analysis was run in duplicates with different starting seeds, each for 100 million generations and saving each 50,000th generation.

To generate comparative information from an independent molecular marker, which can be advantageous for identifying incomplete lineage sorting and making estimates of topological uncertainty and divergence times (Leaché et al., 2019), a species tree with StarBEAST2 was also inferred for the mitochondrial *16S* gene. Taxon sets (“species”) corresponding to the main haplogroups (as inferred by the ML tree; Fig. S2, Table S1) with pronounced geographical confines to mountain massifs resulted in 14 terminals plus the nearest outgroup, *P. cricogaster*. The site model HKY with proportion of invariant sites (0.789) was selected as the best-fit model using the Bayesian information criterion in jModelTest v2.1 (Darriba et al., 2012). An uncorrelated log-normal relaxed clock (ucln) was applied as preliminary analyses inferred the parameter of the standard deviation of ucln not close to zero (Drummond and Bouckaert, 2015). Gene ploidy was set to haploid (1.0). The time calibrations, tree prior, and other settings were the same as in the genomic analysis.

In both analyses, the first 10% of samples were discarded as burn-in after an inspection in Tracer v1.7 (Rambaut et al., 2018), which confirmed satisfactorily high ESS values in all parameters. The posterior probability distributions of species trees were visualized in DensiTree v2.2 (Bouckaert and Heled, 2014). The post burn-in samples were combined in the BEAST module LogCombiner, and another BEAST module, TreeAnnotator, was used to infer the final species tree as a maximum clade credibility tree (MCCT) with node ages calculated as common-ancestor heights and confidence intervals with 95% highest posterior densities (HPD).

### 2.2. Morphology

#### 2.2.1. Morphometrics

Body size (snout-vent length, SVL; snout-urostyle length, SUL) and 27 morphometric variables (including nine lengths of digits) were measured in the type series of the new species (as defined below) using a digital caliper to the nearest 0.1 mm. Eighteen taxonomically potentially important variables (Zimkus and Gvoždík, 2013; Watters et al., 2016) were measured in all available adult individuals (52 males/63 females; males have small spines on the plantar and ventral tarsal surface, Blackburn, 2010) representing all three taxa and six mitochondrial lineages of the *P. steindachneri* species complex. Specimens are stored in collections of the National Museum in Prague (NMP), Harvard University’s Museum of Comparative Zoology (MCZ), and Zoological Research Museum Alexander Koenig in Bonn (ZFMK). For a specimen list, and a list and definitions of the morphometric variables and their graphical illustrations see Supplementary material (Table S1, Fig. S1, Appendix S1).

#### 2.2.2. Statistical analyses

Basic descriptive statistics were performed for the measured variables. Body size variation was visualized using boxplots, and the one-way analysis of variance (ANOVA) was used to compare selected groups. For multivariate morphometric analyses of body shape variation, the variables were corrected for body size using the geometric means method of Mosimann (1970). A size index of each individual was calculated as the arithmetic mean of natural logarithms of all 18 variables (equivalent to the geometric mean of original variables). The individual’s size index was then subtracted from the value of each logarithmized variable. Such data transformation and size-standardization remove the effect of body size, and therefore eliminates possible effects of different age and sexual size dimorphism. One size-corrected variable (size-adjusted SUL) was excluded from the following multivariate morphometric analyses to obtain linearly independent shape variables. The principal component analyses (PCAs) were applied to assess variation within the size-corrected variables using the R package *vegan* v2.5-6 (Oksanen et al., 2019). To test sexual dimorphism in body shape, we compared males and females by two-way t-tests on the first two principal components generated from a dataset comprising all adult individuals. Five consecutive independent PCAs were performed to explore the body shape variation in the following hierarchical design: (PCA 1) present taxonomic identifications corresponding to the three taxa, *P. jimzimkusi, P. njiomock*, and ‘*P. steindachneri*’; (PCA 2) four mtDNA lineages of ‘*P. steindachneri*’ A-D, including genetically untested individuals from the Gotel Mts. (ZFMK) assignable to either the lineage A or D (abbreviated AD); (PCA 3) four species as newly defined in this study, without hybrid and unassignable individuals; (PCA 4) *P. steindachneri* sensu stricto (s.s.), *Phrynobatrachus* sp. nov. (as defined below; abbreviated ami), and their hybrids; (PCA 5) individuals from Mt. Oku (including surrounding areas, i.e. Oku Massif) representing four categories, *P. jimzimkusi, P. njiomock*, hybrids of *P. njiomock* and *P. jimzimkusi* (all specimens from the site where a hybrid individual was detected), and *Phrynobatrachus* sp. nov. Multivariate analyses of variance (MANOVAs) were performed on the first three principal components of each PCA analysis to test body shape differentiations among the groups as shown graphically (Figs. 6, S5). The linear discriminant analyses (LDAs) performed in the R package *MASS* (Venables and Ripley, 2002) were applied (LDA 1) to inspect the level of differentiation in body shape of the four newly defined species, and (LDA 2) to distinguish specimens from within the Gotel Mts., from where genetically untested individuals of potentially two lineages/species were available (ZFMK). All statistical analyses were carried out in the program language and environment R v3.6.3 (R Core Team, 2020).

## 3. Results

### 3.1. Genetics

#### 3.1.1. mtDNA

The ML analysis of *16S* resulted in a tree that corresponded well to the previously published results with respect to the six main lineages within the *P. steindachneri* complex (Zimkus and Gvoždík, 2013; Gvoždík et al., 2020), i.e. *P. jimzimkusi*, *P. njiomock*, and ‘*P. steindachneri*’ A–D (Fig. S2). The main lineages were highly supported (bootstrap ? 83), except *P. njiomock* with a bootstrap value of 54. The tree was divided into two main clades, *P. jimzimkusi* + *P. njiomock* vs. ‘*P. steindachneri*’ A–D, with the latter being divided into A+B and C+D, although the topology received only intermediate to low support (bootstrap 40–69). According to the phylogenetic relationships of individual haplotypes, 14 haplogroups (clusters of closely related haplotypes) were detected: *P. jimzimkusi* (5), *P. njiomock* (2), ‘*P. steindachneri*’ A (1), B (1), C (2), and D (3), see Table S1. Most of the haplogroups had restricted geographic distributions. Interestingly, the microendemic *P. njiomock* was found to be comprised of two haplogroups, with one currently only known from a forest distant from the core distribution range around Lake Oku. Genetic distances among the main lineages are shown in Table 2.

**Table 2.** Genetic uncorrected *p*-distances (in percentages) averaged among and within (on diagonals) species/lineages of *Phrynobatrachus* based on the fragment of the mitochondrial 16S rRNA gene (below diagonals), and concatenated genomic anchored hybrid enrichment data without hybrids (above diagonals). Intraspecific distances are in bold. The upper part is based on the complete deletion (16S: 508 bp; genomic data: 424,500 bp) and the lower part on the pairwise deletion (16S: 510 bp; genomic data: 571,936 bp) approaches in treating gaps and missing data.

| **Complete deletion** | *B steindachneri* | *A steindachneri* | *D sp. nov.* | *C sp. nov.* | *njiomock* | *jimzimkusi* | *cricogaster* | *mbabo* | *batesii* |
| --- | --- | --- | --- | --- | --- | --- | --- | --- | --- |
| *B steindachneri* | **0.1/0.02** | **0.03** | 0.12 | 0.11 | 0.26 | 0.25 | 0.71 | n.a. | 0.90 |
| *A steindachneri* | **3.5** | **0.0/0.01** | 0.11 | 0.10 | 0.25 | 0.24 | 0.71 | n.a. | 0.89 |
| *D sp. nov.* | 3.1 | 2.6 | **0.3/n.a.** | **0.06** | 0.26 | 0.25 | 0.73 | n.a. | 0.91 |
| *C sp. nov.* | 2.8 | 2.5 | **1.3** | **0.1/0.03** | 0.24 | 0.23 | 0.72 | n.a. | 0.90 |
| *njiomock* | 3.0 | 3.4 | 2.5 | 2.2 | **0.1/0.01** | 0.12 | 0.74 | n.a. | 0.95 |
| *jimzimkusi* | 4.1 | 3.6 | 2.9 | 2.6 | 2.0 | **0.2/0.06** | 0.72 | n.a. | 0.93 |
| *cricogaster* | 6.2 | 7.1 | 6.6 | 6.3 | 5.5 | 5.8 | **n.a.** | n.a. | 0.97 |
| *mbabo* | 4.8 | 3.9 | 3.3 | 3.1 | 3.4 | 3.4 | 5.1 | **n.a.** | n.a. |
| *batesii* | 6.8 | 7.1 | 7.0 | 6.5 | 5.5 | 6.3 | 6.1 | 4.7 | **n.a.** |
| **Pairwise deletion** |  |  |  |  |  |  |  |  |  |
| *B steindachneri* | **0.1/0.07** | **0.13** | 0.36 | 0.40 | 0.69 | 0.63 | 1.06 | n.a. | 1.27 |
| *A steindachneri* | **3.7** | **0.0/0.07** | 0.32 | 0.36 | 0.67 | 0.61 | 1.05 | n.a. | 1.26 |
| *D sp. nov.* | 3.1 | 2.8 | **0.3/n.a.** | **0.17** | 0.64 | 0.58 | 1.07 | n.a. | 1.28 |
| *C sp. nov.* | 2.8 | 2.5 | **1.3** | **0.1/0.11** | 0.57 | 0.53 | 1.05 | n.a. | 1.25 |
| *njiomock* | 3.0 | 3.5 | 2.5 | 2.2 | **0.1/0.03** | 0.29 | 1.09 | n.a. | 1.33 |
| *jimzimkusi* | 4.0 | 3.8 | 2.9 | 2.6 | 2.0 | **0.2/0.16** | 1.01 | n.a. | 1.26 |
| *cricogaster* | 6.2 | 7.3 | 6.6 | 6.3 | 5.5 | 5.8 | **n.a.** | n.a. | 1.15 |
| *mbabo* | 4.8 | 4.1 | 3.3 | 3.1 | 3.3 | 3.4 | 5.1 | **n.a.** | n.a. |
| *batesii* | 6.8 | 7.3 | 7.0 | 6.5 | 5.5 | 6.3 | 6.1 | 4.7 | **n.a.** |

#### 3.1.2. Genomic concatenated trees

The ML analysis of the genomic data including all individuals resulted in a tree with all the lineages monophyletic, except lineage D, which formed a ladder-like cluster in between C and A+B (Fig. 2A). The general topology of the main lineages corresponded to the mtDNA topology, except C vs. D, and received the full bootstrap support (100) in all nodes. Beside D, all main lineages were highly supported (bootstrap ≥ 82) except for the lineage A, which received only a weaker support (55). Interestingly, the ladder-like topology of individuals from the lineage D received high support values (82–92). Within D, geographically, the single individual from the Mambilla Plateau was in the basal position in relation to the remaining three individuals from the Gotel Mts. One individual of *P. njiomock* originated from the forest on the northeastern slope of Mt. Oku (a site 8 km distant from the core distribution spot around Lake Oku) was in the highly-supported (bootstrap 82) sister relationship to the remaining two individuals (one from Lake Oku, one from 2 km afar from the Lake), though the divergence was relatively deep. Geographically, four of the five individuals originated from zones of sympatry of several lineages (Gotel Mts. and Mt. Oku), only the single D individual from the Mambilla Plateau did not originate from a known zone of sympatry. Therefore, these individuals with the main focus on the four individuals from the zones of sympatry were tested to identify if they might represent hybrids (see below).

When these four individuals were removed, the resulting tree had a ‘usual’ dichotomic topology that did not exhibit the ladder-like topology (Fig. 2B). Again, all the main lineages were found in the same topology as the mtDNA phylogeny, and now received full support (bootstrap 100). The sole D individual (from the Mambilla Plateau) was in the sister relationship to the four C individuals. In comparison to the mtDNA divergences, the divergences in the genomic data had a different pattern, with the relative depth of the split between A and B substantially smaller (approximately 6x). Genetic distances are shown after the pairwise and complete deletions of gaps and missing data in Table 2.

#### 3.1.3. Detection of hybridization

The four putative hybrids from the two zones of sympatry were embedded in the genomic concatenated ML tree (Fig. 2A) in a ladder-like cluster (three specimens from Gotel Mts., D_Gotel_) or an unusually divergent position (one individual from Mt. Oku, *nji*_Forest_). Their genomic compositions were inferred as hybrid indices between a reference and alternative population (Table 1). This was executed in several combinations based on phylogenetic and biogeographic rationales. Hybrid index was also inferred for the single individual from the Mambilla Plateau (D_Mambilla_) to examine its genomic composition in respect to its phylogenetic and biogeographic proximities to the putative hybrids from the Gotel Mts. The three individuals from the Gotel Mts. (D_Gotel_) clearly showed a mixed genetic background between the clades containing the lineages A+B and C+D_Mambilla_ (HI, hybrid index 0.37-0.68), with two individuals having the genomic profile more similar to A+B and one individual to C+D_Mambilla_. When the three D_Gotel_ individuals were tested against A and B (same phylogenetic distance and geographic proximity), they showed closer similarity to the sympatric lineage A (HI 0.77-0.78). The single individual from the Mambilla Plateau (D_Mambilla_) when tested between A+B and C showed only a small affinity to the first, being obviously closer in its genomic profile to the lineage C (HI 0.77). The phylogenomically divergent individual from the forest on the north-eastern slope of Mt. Oku having mtDNA of *P. njiomock* (*nji*_Forest_) also showed a mixed genomic profile. It was tested between *P. njiomock* from around Lake Oku (*nji*_Lake_) and i) *P. jimzimkusi*, ii) ‘*P. steindachneri*’ C+D_Mambilla_, and iii) ‘*P. steindachneri*’ C_Oku_. The inferred hybrid indices (0.41, 0.64, and 0.64) showed this individual having more genomic information from *P. jimzimkusi* when compared to *P. njiomock* (0.41), and more genomic information from *P. njiomock* when compared to ‘*P. steindachneri*’ C+D_Mambilla_ (0.64), and C_Oku_ (0.64).

The subset of 20 loci, which were phased to gametic haplotypes also supported the hypothesis of the four individuals from the two zones of sympatry representing hybrids. Twenty ML trees showed a tendency of the two gametic haplotypes to segregate in different parental lineages, specifically the D_Gotel_ individuals in C or D_Mambilla_ and A or B, and the hybrid from Mt. Oku mainly in *P. njiomock* and *P. jimzimkusi*, but in a lesser extent also in ‘*P. steindachneri*’ C (especially haplotypes from Mt. Oku in the latter two).

#### 3.1.4. Genomic species tree and species network

To empirically explore what happens to a species-tree analysis when hybrids are included, we applied two approaches - ASTRAL-III analyses without and with hybrids (Fig. 3). Interestingly, both resulting species trees received full support in all nodes (quartet support = 1.0) but differed in both topology and branch lengths. The hybrid individuals encoded by their mtDNA identities caused shortening of the divergence depth between *P. jimzimkusi* and *P. njiomock* (hybrid from Mt. Oku, *nji*_Forest_), and a paraphyly of ‘*P. steindachneri*’ C and D (hybrids from Gotel Mts., D_Gotel_), with the latter sister to a clade containing ‘*P. steindachneri*’ A and B. The general topologies of both species trees are congruent with the ML topologies of the main lineages (Fig. 2). The species tree without hybrids (Fig. 3B) is also congruent in its topology with the topology of mtDNA tree (Figs. 1, S2), but relative branch lengths differ. The branch leading to *P. njiomock* is remarkably long (tree is in coalescent units). The supplemental ASTRAL-III analyses performed with each individual treated as a “species” brought virtually the same and highly supported results, also congruent with the phylogenetic patterns received from the ML analyses, excluding hybrids (Fig. S3) and including them (not shown). The hybrid *nji* Forest was again embedded between *P. njiomock* (to which it was in the highly supported sister position) and *P. jimzimkusi*. The hybrids D_Gotel_ formed—similarly to the ML tree (Fig. 2A)—the ladder-like cluster between C+D_Mambilla_ and A+B. The only difference in the comparison to the ML tree was that the D_Mambilla_ individual was sister to lineage C (with the full support 1.0).

**Fig. 3.**
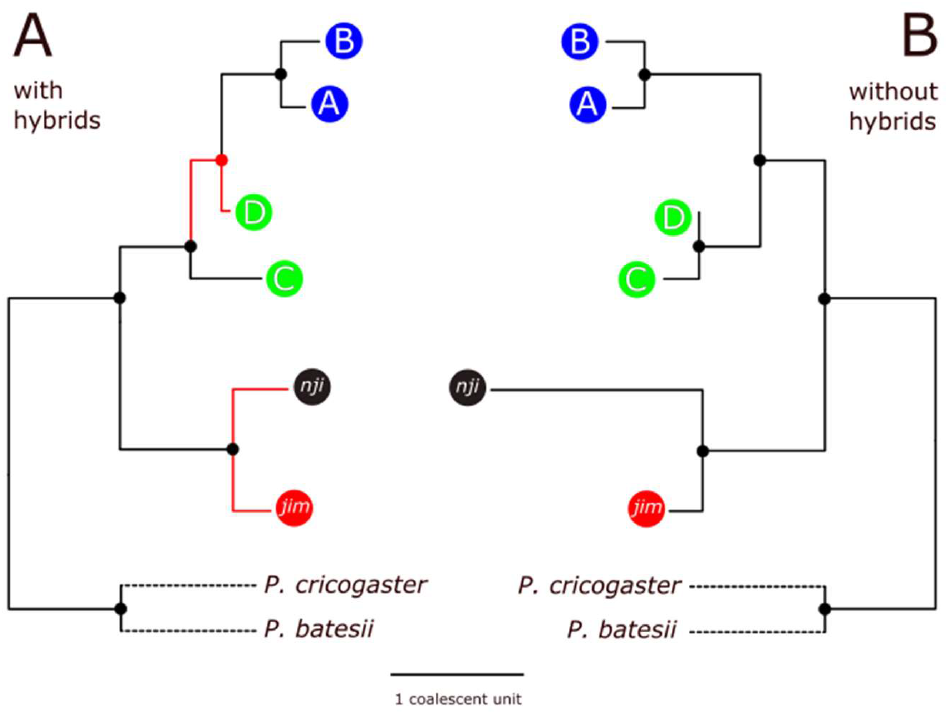
Coalescent-based species trees (ASTRAL-III) based on sequence-capture genomic data of the six terminals corresponding to the six mtDNA lineages. Individuals were encoded by mtDNA-lineage identity, (A) with hybrids included to demonstrate how they affect the species tree, and (B) without hybrids. All nodes received high supports (full dots). Red branches and dot highlight the differences in the topology and branch lengths.

The Bayesian species-network analysis (SpeciesNetwork) resulted in 13,304 network topologies. The topologies with highest posterior probabilities are consistent in reticulation patterns, which were inferred most frequently between *P. jimzimkusi* and *P. njiomock*, and ‘*P. steinachneri*’ (C and/or D) and (A and/or B). A network with the highest posterior probability is shown in Fig. 4. The species-network reconstruction is in general congruence with the other phylogenomic inferences. The deepest split was inferred at 2.5 Mya, separating a clade containing *P. jimzimkusi* and *P. njiomock* (split at 0.9 Mya) and a clade containing ‘*P. steindachneri*’. The latter split at 1.4 Mya into clades containing C+D (split at 0.6 Mya) and A+B (split at 0.2 Mya). Two ghost lineages were inferred in between C+D and A+B at about 0.8 and 0.3 Mya, with estimated reticulation events connecting A, D, and the ghost lineages. A reticulation event was also inferred between *P. jimzimkusi* and *P. njiomock* at 0.6–0.4 Mya. This analysis further supports the existence of gene flow within the *P. steindachneri* species complex, and specifies the most likely connections.

**Fig. 4.**
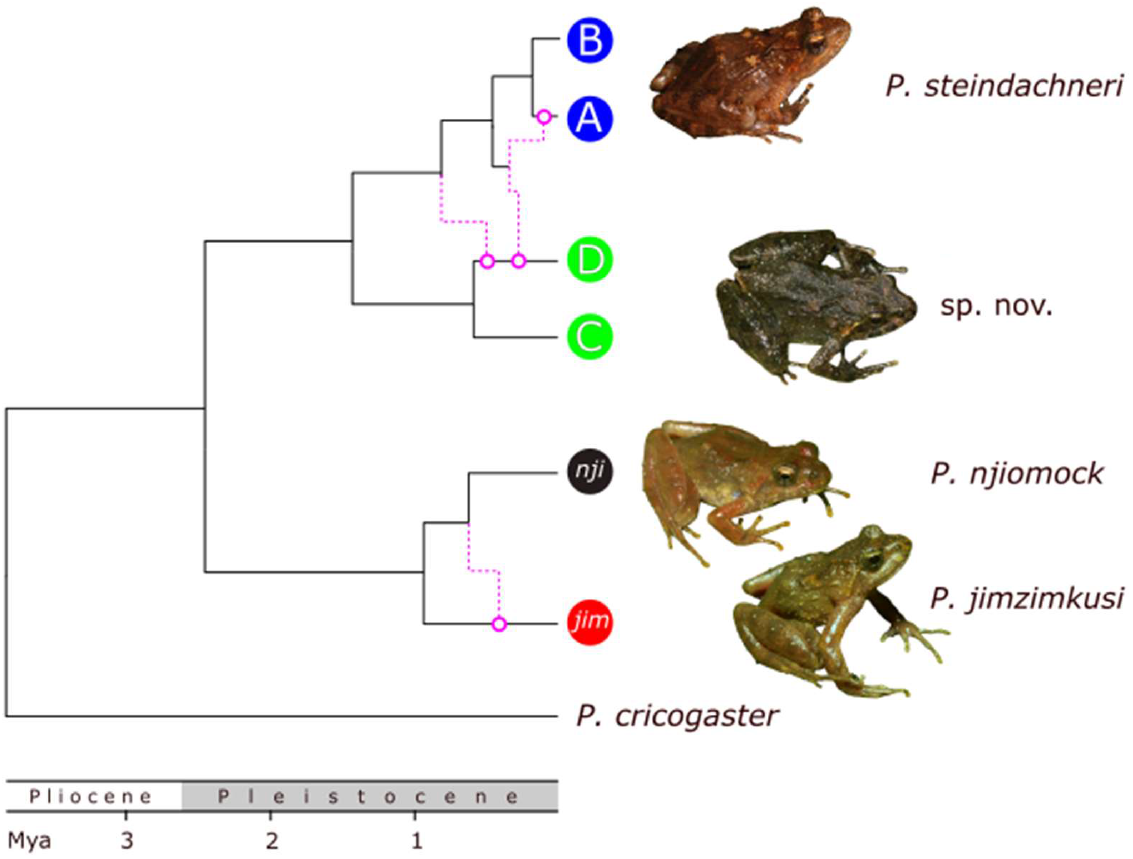
Bayesian coalescent-based species network (SpeciesNetwork). Hybridization events are highlighted by purple and correspond temporally to the Middle and Late Pleistocene climatic oscillations. The splitting pattern and depths of divergences are in a support of four species: *P. jimzimkusi (jim), P. njiomock (nji), P. steindachneri* (A+B), and *P*. sp. nov. (C+D). Photos by Václav Gvoždík.

#### 3.1.5. Phylo geo graphy

Phylogenetic relationships among populations from different mountain massifs of the Cameroon Highlands (Fig. 5) were inferred from both the nuclear genomic data (Fig. 5E) and for comparison, from mtDNA (Fig. 5F), which allowed a broader geographic and specimen sampling. Both trees were reconstructed with the same main topology, the genomic dataset receiving full support (posterior probability = 1.0) for all nodes except the intraspecific relationships within *P. jimzimkusi* (among Mt. Oku, Mt. Bamboutos, and Mt. Manengouba). In mtDNA, the node support values were lower, substantially high only in the lineages of *P. jimzimkusi*, *P. njiomock*, their common clade, and the lineages ‘*P. steindachneri*’ C and D. This is not surprising, when we consider the short length of the studied fragment (512 bp). Within the montane-forest endemic *P. steindachneri* species complex, the deepest split was found separating the southern and northern mountains with the division border on Mt. Oku (separating *P. jimzimkusi* + *P. njiomock* and ‘*P. steindachneri’)*. The split is dated to 2.7 Mya (95% HPD: 3.5–1.8 Mya) in nDNA [2.3 (3.0–0.8) Mya in mtDNA; splits in mtDNA are further given in brackets]. The next split dated to 1.5 Mya (2.0–1.0) [1.7 (2.4–0.4)] separated the northernmost mountains (Tchabal Mbabo, Gotel Mts.; ‘*P. steindachneri*’ A + B) and the middle mountains (Mt. Mbam, Mt. Oku, Mambilla Plateau; ‘*P. steindachneri*’ C + D) with the phylogeographic break in the Gotel Mts. The microendemic lineage restricted to Lake Oku on Mt. Oku (*P. njiomock*) split from the common ancestor with *P. jimzimkusi* at 0.9 Mya (1.2–0.5) [1.2 (1.9–0.5)]. A substantial differentiation between nDNA and mtDNA trees is in the depths of intraspecific diversifications. Origins of all intraspecific diversifications (when A+B and C+D are considered as two species, see below) were inferred relatively consistently in genomic data at around 0.4 Mya (0.8–0.1). In mtDNA, the split between ‘*P. steindachneri*’ C and D is dated to 0.9 Mya (1.9–0.2), and between ‘*P. steindachneri*’ A and B even to 1.5 Mya (2.0–0.2). The wider sampling of *P. jimzimkusi* for the mtDNA dataset allowed us to gain additional information regarding the phylogeography of the southern mountains. Some haplotypes or haplogroups were found to be present on more mountains, but there is a clear geographic signal in the phylogenetic relationships of *P. jimzimkusi* populations (Figs. 5F, S2). The basal split separates populations from Mt. Manengouba + Obudu Plateau and Mt. Bamboutos + Mt. Lefo + Mt. Oku and is dated to 0.5 Mya (1.0–0.1), and subsequent diversifications date to ca. 0.3–0.1 Mya.

**Fig. 5.**
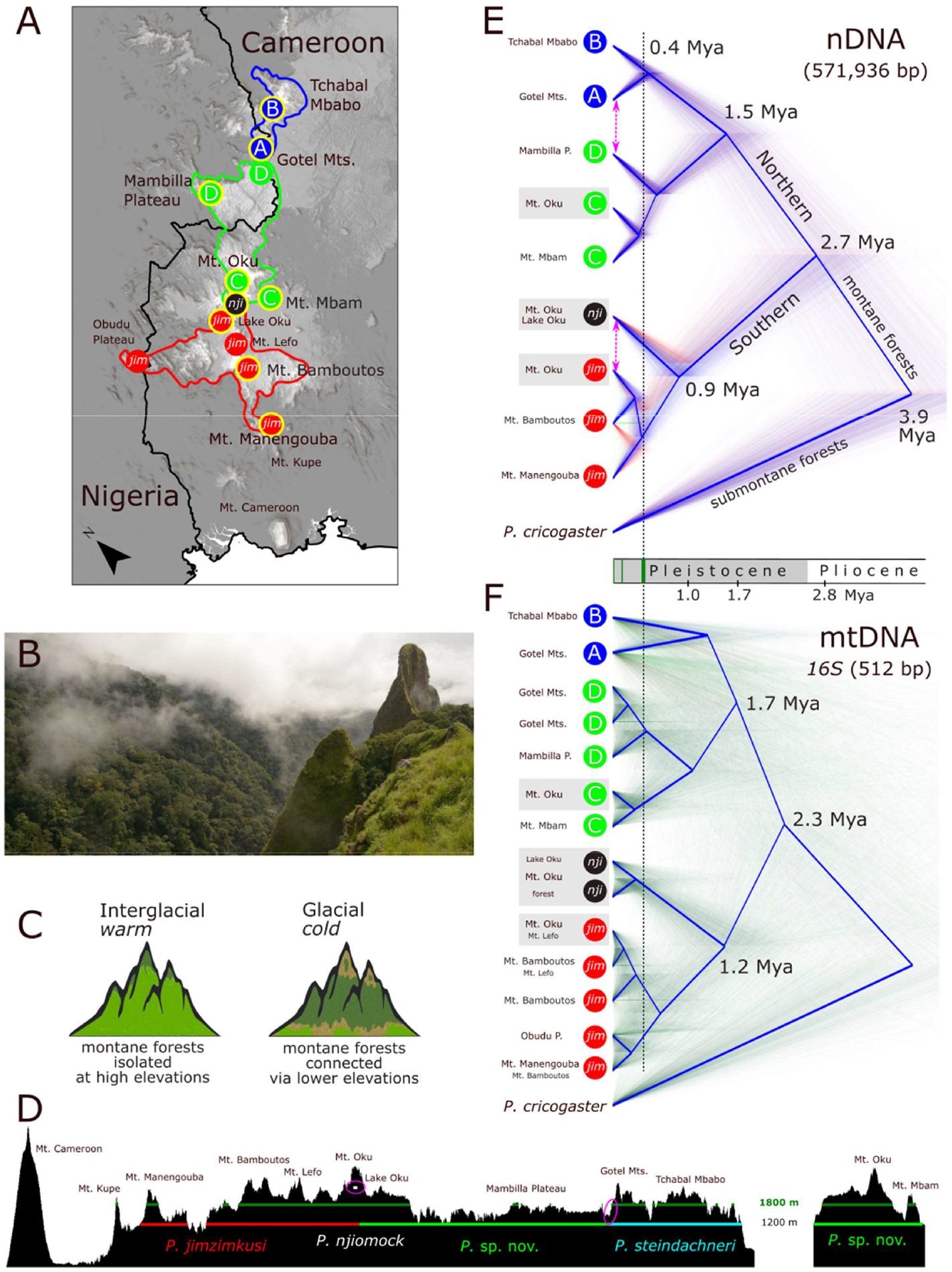
Phylogeography of the Cameroon Highlands forests as inferred from the model, puddle frogs of the *Phrynobatrachus steindachneri* complex. (A) Map showing three main groups of mountain blocks corresponding to the distribution of three species: *P. steindachneri* (blue), *P*. sp. nov. (green), and *P. jimzimkusi* (red); and highlighting Mt. Oku, where three species are in contact, including the endemic *P. njiomock* (black). Samples from sites encircled by yellow were included in the genomic species-tree analysis. (B) Montane forests in the Gotel Mts. (C) Pictograms showing scheme of connections/disconnections of montane forests. We hypothesize that montane forests were probably more connected during cold glacial periods, when lowland rainforests (light green) were retracted and cold-adapted montane forests (dark green) could spread to lower elevations, particularly to wet valleys. (D) Elevational profile of the Cameroon Volcanic Line showing main mountains (sampling sites), distributions of individual species (present-day lower elevational limit of the *P. steindachneri* complex is approx. 1200–1300 m a.s.l.), lower limit of montane forests at 1800 m a.s.l. (White, 1983; dark green line; nonetheless, montane forests descend as stream-side fringing gallery forests to lower elevations), and regions of hybrid zones on Mt. Oku and in the Gotel Mts. (purple ellipsoids). Mt. Mbam is shown separately as lying out of the main axis. (E–F) Dated species trees (StarBEAST2) based on the (E) genomic data and (F) mtDNA shown as “cloudograms” of post-burnin trees with root canal. Tree terminals are based on mountain blocks while respecting phylogenetic relationships (e.g., Mt. Oku is shown per each species; grey rectangles) and mtDNA haplogroups. The last major crown-group splits correspond approximately to the 50,000 years-long interglacial period about 400 kya. The Middle and Late Pleistocene climatic oscillations facilitated gene flows between divergent lineages/species (purple arrows). Time scale shows the geological epochs, periods of increased aridity in Africa (DeMenocal, 1995, 2004), and the last major interglacials (green bars).

### 3.2. Morphology

Body size of the four species as newly defined in this study (see Discussion) as average SVL (mm) in adult males/females are as follows: *P. jimzimkusi* (30.6/31.4), *P. njiomock* (26.5/27.5), *P. steindachneri* s.s. (29.3/29.1), and *Phrynobatrachus* sp. nov. (29.5/28.5), Fig. S4. The largest *P. jimzimkusi* differs significantly from the new species in SVL of females (ANOVA: F = 8.582, p = 0.006). Sexual dimorphism in body shape was not found significant (PC1: *t* = 1.93, *p* = 0.056; PC2: *t* = 0.64, *p* = 0.521), and therefore males and females were analyzed together in the subsequent multivariate analyses. Factor loadings of the first three principal components (broken-stick model identified mainly the first two PCs as the most important; Jackson, 1993) and results of MANOVAs (highly significant in all cases) are given in Table S2, and coefficients of the linear discriminants in Table S3. In general, head (HW, ED, EPD) and limb variables (RL, FT, TL) were the most contributing to the overall variation. PCA 1: PCA of all individuals (Fig. S5A) showed a substantial overlap of *P. jimzimkusi* and *P. njiomock*, with ‘*P. steindachneri*’ only partly overlapping the other two in the morphospace. PCA 2: PCA of the four mtDNA lineages of ‘*P. steindachneri*’ A-D (Fig. S5B) showed a partial overlap of all lineages with the genetically untested individuals from the Gotel Mts. mostly overlapping with ‘*P. steindachneri*’ A. PCA 3: When the four species as newly defined in this study were evaluated by PCA (Fig. S5C) and LDA 1 (Fig. 6A), a substantial overlap was found between the new species (“ami”) and *P. jimzimkusi* in both methods, and in PCA also between the new species and *P. njiomock*. PCA 4: The same PCA as PCA 2 but graphically displaying *P. steindachneri* s.s., the new species and their hybrids (Fig. 6B) uncovered a partial overlap between *P. steindachneri* s.s. and the new species, with their hybrids located in between the two. The genetically untested individuals from the Gotel Mts. were mostly classified using LDA 2 as *P. steindachneri* s.s. (20/24 cases), less often as the new species (Table S5). In most of the cases the discrimination was inferred with a high probability (≥ 95%), but in some cases the discrimination was less successful (~ 50%), and these individuals could represent hybrids. PCA 5: The last PCA was performed on individuals from the Oku Massif (Fig. 6C), where the new species displayed a substantial overlap with *P. njiomock*, and the hybrids of *P. njiomock* and *P. jimzimkusi* showed intermediate position in between the two parental species. Basic descriptive statistics of the measured variables is given in Table S6.

**Fig. 6.**
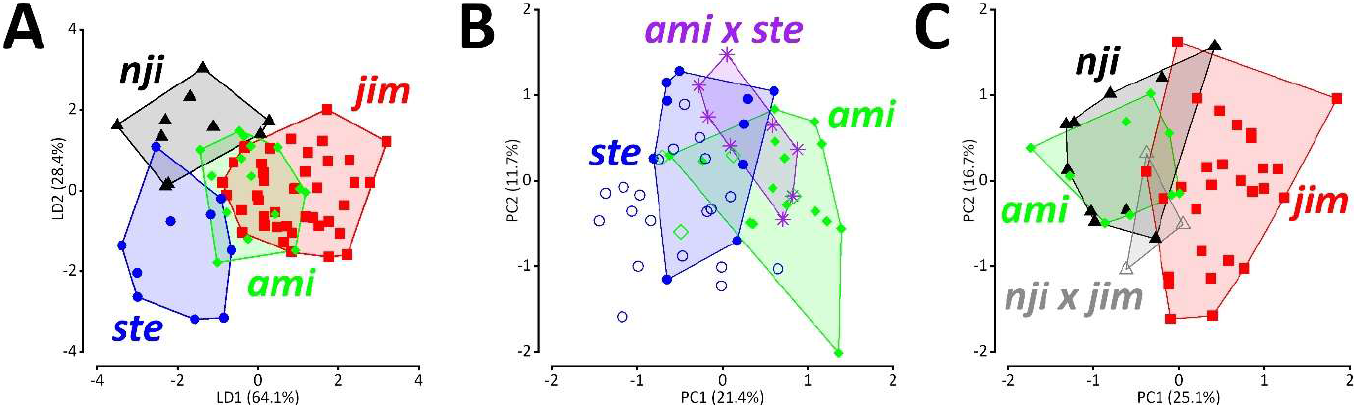
Multivariate morphometrics showing the variation in body shape of the species as newly defined in this study and hybrid individuals: *P*. sp. nov. (*ami*), *P. jimzimkusi* (*jim*), *P. njiomock* (*nji*), *P. steindachneri* (*ste*). (A) LDA 1, genotyped samples of the four species. (B) PCA 4, genotyped samples (full symbols and stars) of the new species, *P. steindachneri* and their hybrids *(ami* x ste), and non-genotyped specimens from the Gotel Mts. (empty symbols) as discriminated by LDA 2. (C) PCA 5, individuals from the Oku Massif representing three sympatric species and Mt. Oku’s hybrid individuals (*nji* x *jim*).

## 4. Discussion

### 4.1. Phylogeny and gene flow

Multilocus phylogenies in some cases bifurcate into a ladder-like pattern, hereafter referred to as LLP (ladder-like phylogeny/pattern), especially in fast evolving organisms like viruses (e.g., Luksza and Lässig, 2014; Colijn and Plazzotta, 2018). The viral imbalanced phylogenies (LLP) are typically result of selection given by the effect of immune escape (Volz et al., 2013). Here, we demonstrate an empirical case of this kind of phylogenetic pattern using genome-wide sequence capture data of a vertebrate (frog) species complex on a phylogeographic scale, i.e. multiple individuals per species (Fig. 2A). Conspecific specimens (of valid or candidate species) were expected to form monophyla, which did happen but in one case. In this case, the individuals branched into the LLP in between two clades. This unexpected pattern opened the question of whether the LLP specimens represented hybrids of the “outer” clades (species), i.e. if they established gene flow between them. Indeed, this hypothesis received a high support based on investigation of genome-wide single nucleotide polymorphisms (Table 1).

Here, we illustrated practical impacts of gene flow on phylogenomic reconstructions, using the maximum likelihood on a concatenated dataset, and a coalescent-based species tree (ASTRAL-III) when including and excluding hybrids (Figs. 2–3). The effect of gene flow on the phylogenetic reconstruction has already received a considerable attention (e.g., Leaché et al., 2014; Solís-Lemus et al., 2016; Long and Kubatko, 2018; Kubatko and Chifman, 2019; Jiao et al., 2020) but is not yet widely accented by empiricists in the systematic research (for exceptions see, e.g., Gottscho et al., 2017; Roos et al., 2019). In our case, hybrid individuals formed the LLP in both ML and the species-tree approach (when each individual was treated as a “species”). More interestingly, in both cases the topologies were inferred with high supports. The only topological difference between ML and the ASTRAL-III species tree was the position of one sample (D_Mambilla_), the sample originally presumed to be conspecific with the LLP individuals based on mtDNA. This sample was either forming the first “step” of the ladder leading to one clade (ML), or it was in the sister relationship to the second clade (species tree). However, when the specimens identified as hybrids were excluded, the discussed sample was consistently placed as the sister lineage (with a shallow divergence) of the second clade in both ML and ASTRAL (Figs. 2B, S3). An inclusion/exclusion of another individual identified as a hybrid of other two clades (*nji*_Forest_) did not change the topology but changed the branch length of the coalescent-based ASTRAL species tree (Fig. 3).

Our study demonstrates empirically how the presence of hybrid individuals may affect estimations of concatenated ML trees and species trees. Identification of hybrid samples in phylogenetic analyses is crucial, especially in species-tree analyses. The species-tree approach has the assumption of no gene flow in data, thus, an inclusion of hybrid samples results in a misleading tree (Rannala et al., 2020). As we demonstrated, the inclusion of hybrids may affect both species trees’ topologies and branch lengths while high supports may be received (Fig. 3). In practice, species assignments are often based on maternally-inherited mtDNA, which is certainly not the correct approach in the case of hybrids. Solís-Lemus et al. (2016) and Long and Kubatko (2018) tested consistency of ASTRAL and other species-tree methods under gene flow and found out that anomalous species trees can be estimated if the assumptions for the species-tree analysis are violated. Recently, it was shown that in species trees with short internal branch lengths even a small amount of gene flow causes species-tree estimations to become inconsistent (Jiao et al., 2020). Here, the LLP is shown as a sign of possible hybridizations in individual-based phylogenetic analyses. Similar ladder-like phylogenies were inferred in previous studies. For example, in the case of *Uma* iguanian lizards, the LLP-forming *U. rufopunctata* was identified as a hybrid population (Gottscho et al., 2017), or in crocodiles, the LLP was formed in hybrids between *Crocodylus acutus* and *C. moreletii* (Pacheco-Sierra et al., 2018).

When the LLP is inferred in a phylogenetic reconstruction, such a result might imply hybridization in the study system. Therefore, we recommend applying the network model of species tree (Solís-Lemus et al., 2016; Degnan, 2018; Long and Kubatko, 2018), instead of bifurcation-based inferences. This approach was also found to be effective with our study system (Fig. 4), which allowed us to estimate phylogenetic, spatial (as the dataset was phylogeographic) and temporal pathways of interspecific gene flows in a simultaneous analysis.

### 4.2. Reticulate evolution of the P. steindachneri species complex

#### 4.2.1. Phylogeny, divergences and species limits

There is a never-ending debate about species concepts and criteria proposed for species delimitation in different groups of organisms (Agapow et al., 2004; de Queiroz, 2007; Fišer et al., 2018; Hillis, 2019). In our research, we applied an integrative approach (Vieites et al., 2009; Padial et al., 2010) by combining three evolutionary independent main lines of evidence (genome-wide nDNA, mtDNA, and phenotype) to study diversification processes of the puddle frogs of the *Phrynobatrachus steindachneri* species complex from Afromontane habitats of the Cameroon Highlands. Since recordings of advertisement calls are available only for a single taxon of the complex (see Taxonomic implications), the analysis of phenotype was limited to the morphological data. There is also only scarce information on the ecology (Zimkus and Gvoždík, 2013), especially when individual populations are treated as several candidate species. Therefore, we focused on the genetic and morphological data.

The null hypothesis was six candidate species correspond to the six mtDNA lineages, *P. jimzimkusi, P. njiomock*, and ‘*P. steindachneri*’ A–D. Mitochondrial DNA phylogeny indicated a presence of two main clades (*P. jimzimkusi* + *P. njiomock*) vs. ‘*P. steindachneri*’ A–D, with the latter divided into A+B and C+D, though the topology received only intermediate to low support (Fig. S2; Zimkus and Gvoždík, 2013; Gvoždík et al., 2020). None of the lineages are known syntopically from a single site, but a sympatry is known from the Gotel Mts. (‘*P. steindachneri*’ A and D) and Oku Massif (*P. jimzimkusi, P. njiomock*, ‘*P. steindachneri*’ C).

The mtDNA marker *16S* has been widely used in anuran systematics as a theoretical indicator of candidate species when uncorrected pairwise divergence >3% (although, sometimes only 1-2% in confirmed species; Vieites et al., 2009). When we compared the average uncorrected *p*-distances among the six lineages of the *P. steindachneri* species complex, most of the comparisons (12/15) were above or near the 3% divergence (Table 2), indicating that the mtDNA lineages could represent species. The deepest divergence was found between the lineages A and B (3.7%, or 3.5% when the complete deletion approach in treating gaps and missing data was applied), which was about 2.8x more than smallest divergence (found between C and D; 1.3%). Based solely on the mtDNA divergence, this information could lead to the conclusion that A and B should be considered two separate species, and many taxonomists would probably proceed as such. However, the 3% divergence threshold in *16S* should not be universally applied, and an integrative taxonomic approach is needed (Vieites et al., 2009; Padial et al., 2010). Therefore, the genome-wide nDNA data were crucial to contrast the mtDNA divergence pattern. The phylogenomic analyses of nDNA (Figs. 2B, 3B, 5E) inferred—with high nodal supports for main clades—the same topology like mtDNA (Fig. 1). However, the divergence depths were substantially different.

It is important to note that there may be methodological issues when dealing with genetic distances based on genomic data. Phylogenomic datasets commonly have a considerable amount of missing data (Roure et al., 2013; in our case, 25.8% of the aligned positions contained missing data). When we applied two different approaches in how to treat gaps and missing data (pairwise and complete deletions), we received substantially different values (Table 2). The distances calculated with the pairwise deletion approach were estimated several times higher (average 3.1x within the ingroup; 2.3-5.5x) than the ones calculated with the complete deletion. This methodological detail on the approach used is often not given in publications. We therefore appeal to authors for providing the necessary technical details when genetic distances are calculated. As the complete deletion approach works only with existing data, in phylogenomic datasets we recommend to apply the complete deletion.

When we compared the complete-deletion uncorrected p-distances based on the anchored hybrid enrichment genomic data within the *P. steindachneri* species complex, the average divergence between the two main clades (*P. jimzimkusi* + *P. njiomock*) vs. ‘*P. steindachneri*’ was 0.25%. When individual lineages were compared, the interspecific divergence between *P. jimzimkusi* and *P. njiomock* was 0.12%, while among the four lineages of ‘*P. steindachneri*’ the divergences ranged from 0.03 to 0.12%. The divergences A-B (0.03%) and C-D (0.06%) were approximately four- and two-times smaller than between A (or B) and C (or D), with the mean 0.11%. Intraspecific and intra-lineage (candidate species) distances ranged from 0.01% (*P. njiomock* and A) to 0.06% (*P. jimzimkusi)*. Following these comparisons, the A-B and C-D genomic divergences fall within the intraspecific span, while the mean 0.11% divergence between (A + B) vs. (C + D) corresponds to the interspecific divergence (as between *P. jimzimkusi* and *P. njiomock*). When contrasting the genetic distances based on mtDNA and nDNA, the most striking difference could be found between ‘*P. steindachneri*’ A and B. As the found shallow divergence in the genome-wide nDNA is in accordance with the intraspecific variation, the deeply divergent mtDNA lineages must be evaluated as deep conspecific lineages (Vieites et al., 2009). In conclusion, the phylogenomic architecture supports four distinct species in accordance with the genomic cluster species concept (Mallet, 1995, 2001): *P. jimzimkusi, P. njiomock, P. steindachneri* sensu stricto (A+B), and a new species (*Phrynobatrachus* sp. nov. as described below, C+D).

#### 4.2.2. Hybridization

Species-network approaches are recommended when gene flow is detected or expected (Degnan, 2018), and are becoming commonly used (e.g., Wen et al., 2016; Mason et al., 2019; Masonick and Weirauch, 2020). Our species-network analysis (Fig. 4) estimated phylogenetic relationships congruent with the classical phylogenetic trees (Figs. 2B, 3B, 5E), but this analysis also estimated probable pathways of gene flows. The most probable gene flows were inferred as occurring between *P. jimzimkusi* and *P. njiomock* (geographically in contact on Mt. Oku), and between the new species and *P. steindachneri* s.s. dated to the Middle and Late Pleistocene. The latter also involved ghost lineages (probably extinct or not yet sampled) phylogenetically positioned in between the two species, which suggest a more complex scenario in the region between the Mambilla Plateau and Gotel Mts. in Nigeria.

Identifications of introgressions by the species network was congruent with results obtained from two additional analytical approaches: estimations of hybrid indices and phasing to gametic haplotypes. Both indicated that the four specimens uncovered already by their phylogenetic positions represented hybrids: three individuals from the Gotel Mts. *(Phrynobatrachus* sp. nov. and *P. steindachneri* s.s.), and one individual from Mt. Oku. The Mt. Oku individual carries a mtDNA haplotype of *P. njiomock*, but *P. jimzimkusi* dominates (59%) in the nuclear genome as based on the hybrid-index estimation (Table 1). The presence of three species on Mt. Oku in close proximity opens a possibility of gene flows occurring in between all of them. Such an indication of a possible hybrid origin of an individual of the lineage C from Mt. Oku (*Phrynobatrachus* sp. nov.) was demonstrated by Gvoždík et al. (2020). When the hybrid individual was tested against the new species, a higher proportion was found in support of *P. njiomock* (64%). Another remarkable finding with the Mt. Oku hybrid was that when the specimen was removed from the ASTRAL-III analysis (Fig. 3), *P. njiomock* showed a relatively long branch (given in coalescent units). We expect that this indicates a rapid evolution in a relatively short period of time in this geographically-restricted (Lake Oku) species. The phylogenetically “unstable” (when concatenated ML included/excluded hybrids) individual from the Mambilla Plateau (lineage D) was found substantially closer to the lineage C (77%) than to A+B (*P. steindachneri* s.s.) based on hybrid indices (Table 1). It was also recovered as the sister lineage of C in both ASTRAL-III analyses (with and without hybrids when all individuals were kept as terminals). Therefore, we assume that the Mambilla individual is closely related to the new species (conspecific), and does not represent a substantially admixed individual.

For the taxonomic implications, it is important to note that according to the classic biological species concept (Mayr, 1970), species are not admixing and producing viable offspring, and hybridization is seen as a breakdown in isolating mechanisms. However, it is long known and widely accepted now, that (typically) closely related and parapatric species often hybridize to some level (Mallet, 1995, 2007; Mallet et al., 2016; Shapiro et al., 2016). For examples in anurans, see Nürnberger et al. (2016), Chan et al. (2017), Peek et al. (2019). This is also the case with the *P. steindachneri* species complex.

#### 4.2.3. Morphology

Morphological analyses showed that *P. jimzimkusi* reaches the largest body size (and longest hind limbs), *P. njiomock* the smallest, and ‘*P. steindachneri*’ is placed in between the two (Fig. S4), without clear differentiations among the four lineages A-D. Color patterns of all nominal taxa and lineages are highly variable, with intraspecific (or intralineage) variation blurring interspecific distinctions (Figs. S8-S9). *Phrynobatrachus njiomock* is the only exception (details below). In terms of body shape, all nominal taxa are also largely cryptic, partly overlapping in their morphospace, with ‘*P. steindachneri*’ being the most differentiated (Fig. S5A). The four lineages A-D also share a common part of the morphospace (Fig. S5B). If we apply the species limits as based on the phylogenomic architecture (see above), the four species are partly differentiated (Figs. 6A, S5C). The new species has a similar body shape like *P. jimzimkusi* and *P. njiomock* (Figs. 6A, C, S5C) but is substantially differentiated from *P. steindachneri* s.s. (Fig. 6B). A relatively high level of the morphological crypsis of this species complex could mirror a conservatism of ecological niches of these mostly parapatrically distributed species (for a similar case in *Hyla* tree frogs, see Gvoždík et al., 2008). However, such a hypothesis must yet be tested.

Interestingly, despite the mostly cryptic morphology of the *P. steindachneri* species complex, the body shapes of the hybrid individuals from both the Gotel Mts. and Mt. Oku are intermediate between the parental species (Figs. 6B-C). For a general appearance of the hybrids, see Fig. S10.

### 4.3. Taxonomic implications

The present publication is registered on ZooBank under the following accession: urn:lsid:zoobank.org:pub: *will be added upon acceptance*

#### 4.3.1. Described taxa

##### 4.3.1.1. Phrynobatrachusjimzimkusi

Three specimens from three mountain massifs included in the genomic sequence-capture analyses supported the mtDNA and morphological species delimitation of *P. jimzimkusi* (Zimkus and Gvoždík, 2013). This species is distributed in the southern mountains of the Cameroon Highlands, specifically from Mt. Manengouba to the Oku Massif, including Mt. Bamboutos (type locality), Mt. Lefo, Obudu Plateau in Nigeria, and surrounding areas above 1200 m a.s.l. (Figs. 1, 5). Hybridization/gene flow from *P. njiomock* is documented from the north-eastern slope of Mt. Oku, as far as 8 km from Lake Oku – the core distribution region of the latter (Fig. 2). A possible hybridization (mitochondrial introgression) with *P. steindachneri* C (= *Phrynobatrachus* sp. nov.) was indicated also in a recent study (Gvoždík et al., 2020) but not yet further confirmed.

##### 4.3.1.2. Phrynobatrachus njiomock

This species was described primarily from around Lake Oku on Mt. Oku (type locality), but individuals were reported also from near streams several km far from the lake based on mtDNA (Zimkus and Gvoždík, 2013). The genomic analyses supported the sister relationship with *P. jimzimkusi* with the depth of divergence dated to approx. 1 Mya, in the Early Pleistocene (Figs. 4–5). Two of the three specimens identified by mtDNA and tested by genomic approach were *P. njiomock*, including an individual from 2 km NW from Lake Oku, while one individual from 8 km NE from the lake was a hybrid of *P. njiomock* and *P. jimzimkusi* (Fig. 2). This is a genomic evidence that *P. njiomock* is almost endemic to Lake Oku and nearest areas. Further from the lake it underlies to gene flow with *P. jimzimkusi*, and possibly also with the species (Gvoždík et al., 2020), which is described below.

##### 4.3.1.3. Phrynobatrachus steindachneri s.s

The species was described based on 28 syntypes collected by F.W. Riggenbach in “Banjo” [= Banyo; Amiet, 1971], Cameroon, at the beginning of the 20^th^ century (Nieden, 1910). Banyo is a town surrounded by mostly savanna habitat at elevation around 1100 m a.s.l. and occasional hills up to 1540 m a.s.l. Surveys of this region did not locate *P. steindachneri* in 2009 and 2016, which was not surprising given that habitats did not seem suitable for this species. However, the syntypes, including the lectotype ZMB 20429 (probably male; Natural History Museum in Berlin; formally designated by Zimkus and Gvoždík, 2013), are labeled as originated from “Banjo-Bezirk” [= Banyo District]. In this broader sense, the nearest suitable habitats are in the Gotel Mts. to the north-west or Tchabal Mbabo to the north-east. If we consider that Riggenbach collected between Banyo and Garoua (330 km to the north) with some other material collected in the region of Tchabal Mbabo (Nieden, 1910; Barej et al., 2010), we conclude that the type material was most probably collected in the Tchabal Mbabo area. Therefore, we propose to restrict the name *P. steindachneri* to the northernmost populations of the species complex (i.e., the clade comprising ‘*P. steindachneri*’ A and B). This species is sister to *Phrynobatrachus* sp. nov. with an inferred divergence of approx. 1.5 Mya, in the Early Pleistocene (Figs. 4–5). The known distribution comprises only montane areas of Tchabal Mbabo and the northern part of the Gotel Mts., including the highest peak Mt. Gangirwal, at elevations 2000–2400 m a.s.l. A survey of Tchabal Gangdaba (90 km to the northeast) did not reveal any *Phrynobatrachus* in 2009 (V. Gvoždík, unpublished). A possible existence of populations in the mountains to the east (as shown by IUCN SSC Amphibian Specialist Group, 2019) has never been confirmed.

On the north-western slopes of the Gotel Mts., *P. steindachneri* hybridizes with *Phynobatrachus* sp. nov. The new species correspond to ‘*P. steindachneri*’ C and D, and is formally described below.

#### 4.3.2. Description of a new species

***Phrynobatrachus sp. nov*.** Nečas, Dolinay, Zimkus & Gvoždík

**ZooBank:** *will be added upon acceptance*

**Synonymy:** *P. steindachneri* C, *P. steindachneri* D (Gvoždík et al., 2020).

**Common Name:** Amiet’s Puddle Frog.

**Etymology:** This species is named in honor of Jean-Louis Amiet, who worked as a professor at the University of Yaoundé for many years, and has dedicated his life to study amphibians, and other fauna and flora of Cameroon.

**Holotype:** MCZ A-138060, adult female (Figs. 7, S6E-F), Cameroon, West Region, Mt. Mbam, stream NNE of a settlement, 5.9823°N 10.7204°E, 2000 m, 28.VII.2006, leg. D.C. Blackburn, K.S. Blackburn, N.L. Gonwouo, M. Talla.

**Paratypes:** MCZ A-138056, male (allotype), Cameroon, West Region, Mt. Mbam, stream SSE of a settlement, 5.9780°N 10.7197°E, 1970 m, 29.VII.2006, leg. D.C. Blackburn, K.S. Blackburn, N.L. Gonwouo, M. Talla; MCZ A-138057-138059, three females, same collection data as holotype.

**Additional material:** MCZ A-138106, 138110 and 138113, three males, Cameroon, North-West Region, Mt. Oku, Elak Oku village, stream through farms, 6.2459°N, 10.5064°E, 1960 m, 16.VIII.2006, leg. D.C. Blackburn, K.S. Blackburn, P. Huang, M. Talla; MCZ A-138104-138105, 138107-138109, 138111-138112, 138114, 138116-138117, ten females, same collection data as the previous males; MCZ A-139608-139609, female and juvenile, Nigeria, Taraba State, Mambilla Plateau, Ngel Nyaki Forest Reserve, next to stream at forest margin, 7.0981°N, 11.0554°E, 1530 m, 14.IV.2009, leg. D.C. Blackburn.

**Definition:** Medium to large-sized *Phrynobatrachus* (SVL = 24.8-32.1 mm) with elongate oval body, pointed snout, well-developed foot webbing; and conspicuous dorsal chevron-shaped glands; very variable color patterns, typically brownish dorsal base and yellowish or white ventral base; venter in males and females marbled with dark blotches, which are larger in females; pectoral region in males dark with dense white (or colorless) small dots extending to gular region, less conspicuous and more often with larger light blotches in females, such blotches occur on ventral sides of arms in some individuals; gular region in posteromedial portion commonly with a lightly-colored line extending from pectoral region, its length varies between individuals and can be also absent; lower jaw lined with white spots along edge; thighs lightly colored with small white/colorless dots on front sides extending from body sides; naris small with raised narial ring; tympanum distinct but not very conspicuous, covered by skin; supratympanic fold present; canthus rostralis moderately sharp; snout relatively long; loreal region slightly concave; maxillary and premaxillary teeth present; vomerine teeth absent; pupil rounded, elliptical; front limbs without webbing; pedal webbing well-developed (two of four phalanges on fourth toe free of webbing); one mid-tarsal, one outer and one inner metatarsal tubercles; adult males bear minute asperities on foot and ventral tarsal surface.

**Description of holotype:** Adult female, SVL = 30.1 mm (Figs. 7, S6E–F). Measurements (mm): SUL 29.7, HW 10.1, HDL 8.9, TD 1.5, ED 3.6, IOD 3.0, EAD 4.8, EPD 7.1, SL 3.5, SNL 1.5, ENL 2.0, IND 3.0, HL 5.7, RL 6.9, MD1 2.4, MD2 2.9, MD3 5.3, MD4 4.2, FL 15.2, TL 15.8, FTL 15.8, PD1 2.8, PD2 2.8, PD3 5.7, PD4 8.2, PD5 5.9, IMTL 1.1, OMTL 0.5. For detailed description see Appendix S2. A female was selected because the only male (young specimen) from the type series (allotype, Fig. S6A–D) is in a stretched position, not optimal for a holotype specimen.

**Fig. 7.**
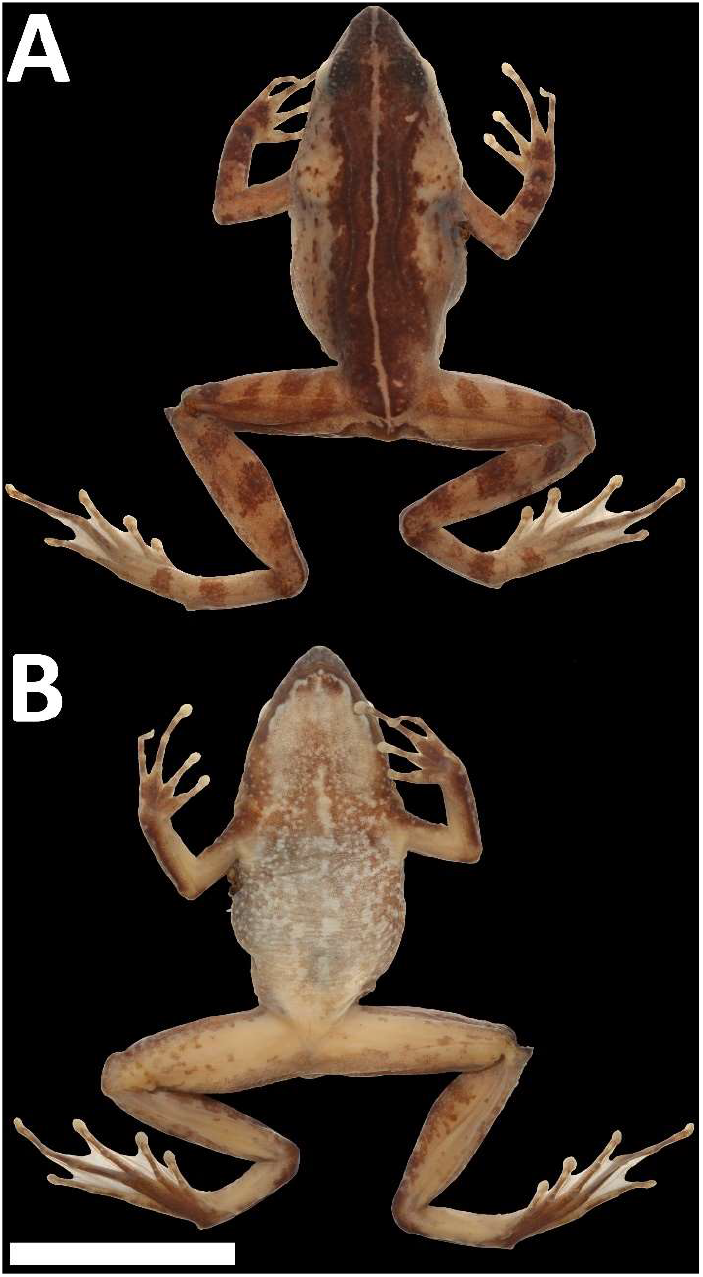
Holotype of *Phrynobatrachus* sp. nov. (MCZ A-138060, female) in (A) dorsal and (B) ventral views. Scale = 15 mm.

**Morphological variation:** SVL 28.8–32.1 mm (males, n = 4), 24.8–31.5 mm (females, n = 11). Relatively low variation in morphological traits except of variable coloration. For detailed description of morphological variation, including color variation, measurements of the type series, and descriptive morphometrics see Supplementary material (Appendix S3, Figs. S7–S8, Tables S4, S6).

**Differential diagnosis:** *Phrynobatrachus sp. nov*. (SVL = 24.8–32.1 mm) can be distinguished from all other (sub)montane species from different species groups of the Cameroon radiation (Gvoždík et al., 2020) by well-developed pedal webbing, larger adult size (SVL in parentheses), absence of intense black coloration of throats in males, and the following characters: from *P. arcanus* (13.1–17.0 mm) and *P. mbabo* (14.0–17.9 mm) by absence of medially clustered enlarged gular asperities in males (*P. arcanus* group, Gvoždík et al., 2020); from *P. chukuchuku* (14.8–19.2 mm) by absence of ventral spines (Zimkus, 2009); and from *P. danko* (19.6–21.8 mm), *P. manengoubensis* (14.1–19.6 mm), *P. schioetzi* (22.6–27.7 mm), and *P. werneri* (17.4–23.0 mm) by absence of pronounced skin folds in gular region along mandibles of males (*P. werneri* group, Blackburn, 2010; Blackburn and Rödel, 2011; Gvoždík et al., 2020). Within the *P. steindachneri* species group, *P. sp. nov*. can be distinguished from *P. cricogaster* (20.0-32.0 mm) by smoother dorsal skin, presence of only one tarsal tubercle, absence of heel spur, and absence of bullseye ventral pattern (Perret, 1966). Within the *P. steindachneri* species complex, *P. sp. nov*. has a diagnostic single nucleotide polymorphism in the studied 16S rRNA barcode [adenine (A) in position 112, instead of cytosine (C) or thymine (T); holotype GenBank: xxxxxxxx]. Morphologically, *P. sp. nov*. is largely cryptic with inter-population variation blurring the inter-species differentiations, yet with a tendency to differ (Figs. 6A, S4, S5C, S8, S9): from *P. jimzimkusi* (25.3-35.7 mm) by smaller average body size (29.5 vs. 30.6 mm in males; 28.5 vs. 31.4 mm in females), by more distinct tympanum, and by relatively shorter hind limbs; from *P. njiomock* (25.4-35.0 mm) by more prominent grey mottling in pectoral and abdominal regions, darker or more colorful ventral coloration, and more verrucose dorsal skin without reddish blotches (Zimkus and Gvoždík, 2013); and from *P. steindachneri* s.s. (26.8-31.6 mm) by relatively narrower head.

**Distribution and natural history:** *Phrynobatrachus sp. nov*. corresponds to the mtDNA lineages C and D of ‘*P. steindachneri*’ (Fig. 1). It is distributed in Cameroon and Nigeria from Mt. Mbam (Fig. S6G-H), across Mt. Oku (at least eastern portion of the Oku Massif) to the Mambilla Plateau and marginally Gotel Mts. (Fig. 5). Documented altitudinal range is 1530–2000 m a.s.l., but it probably lives also beyond these limits along streams surrounded by forests or at least shaded by bushes. It is documented in this study that hybridization occurs with *P. steindachneri* s.s. on the north-western slopes of the Gotel Mts. at elevations around 2000 m a.s.l. Hybridization presumably occurs also on Mt. Oku with either *P. jimzimkusi* or *P. njiomock* or both (see Gvoždík et al., 2020) but has not yet been well documented. *Phrynobatrachus sp. nov*. was observed active in leaf litter along stream edges as well as directly in streams during both daylight (morning) and night, but calls were not heard (D.C. Blackburn, pers. comm.).

Advertisement call remains unknown as it is the case in *P. njiomock* and *P. steindachneri* s.s. Calls of “*P. steindachneri*” analyzed and presented by Amiet and Goutte (2017) and briefly described by Channing & Rödel (2019) correspond to *P. jimzimkusi* based on the locality data. Similarly, larval stadia (tadpoles) have not been reliably recorded but supposedly inhabits calm passages of streams like in *P. jimzimkusi*. The tadpole of *P. jimzimkusi* was described by Pfalzgraff et al. (2015). The previously described tadpole of “*P. steindachneri*” (Channing et al., 2012) must be taken with caution as it is unclear which species it represents (Gvoždík et al., 2020).

Recorded sympatric anuran genera: *Astylosternus, Cardioglossa, Leptodactylodon*, and *Leptopelis* (D.C. Blackburn, pers. comm.).

**Threat status:** *Phrynobatrachus sp. nov*. is categorized as Critically Endangered following the IUCN criteria (IUCN Standards and Petitions Committee, 2019), in line with the other species of the *P. steindachneri* species complex. See *Conservation concerns* for more details.

### 4.4. Conservation concerns

Populations of the newly described species, *P. sp. nov*., have completely disappeared during the last decade (thus far discussed under the name “*P. steindachneri”)*. Not a single specimen has been found on Mt. Oku since 2009 (Doherty-Bone and Gvoždík, 2017), on Mt. Mbam since the type series was collected in 2006 (contrary to Tchassem et al., 2019, where *P. natalensis* was misidentified as “*P. steindachneri*”, according to the published photo). No representatives of the *P. steindachneri* complex were found in the Gotel Mts. during two surveys in 2016 (June, October), and the same situation has occurred with *P. steindachneri* s.s. on Tchabal Mbabo in June 2016 (Gvoždík et al., 2020). No data are available from the Mambilla Plateau since *P. sp. nov*. was collected there in 2009, but it is highly probable that this severe frog decline is widespread throughout the Cameroon–Nigeria mountains (Hirschfeld et al., 2016). Thus, based on all available evidence (references above, and unpublished data), *P. sp. nov*., similarly like *P. jimzimkusi, P. njiomock*, and *P. steindachneri*, may be on the brink of extinction. There are likely many causes for these dramatic montane frog declines (Tchassem et al., 2020), with the chytrid pathogenic fungus being probably a major catalyst (Hirschfeld et al., 2016; Gvoždík et al., 2020). Additional surveys and conservation actions are needed in the Cameroon Highlands to understand the causes of the amphibian populations declines and potentially reverse them.

### 4.5. Biogeography of the Cameroon Highlands forests

#### 4.5.1. Brief overview

The Cameroon Volcanic Line is known as an ancient refugial region serving as an endemism hotspot of animal and plant species with ecological affinities to both montane rainforest (Missoup et al., 2012, 2016; Zimkus and Gvoždík, 2013) and the lowland rainforest surrounding these mountains (Bohoussou et al., 2015; Charles et al., 2018; Leaché et al., 2019; Migliore et al., 2019). In some cases, the CVL was identified as a refugial source for longdistance dispersals (Demenou et al., 2020) or an important biogeographic barrier (Nicolas et al., 2012; Portik et al., 2017) of lowland rainforest biota. Interestingly, the CVL was identified as a biogeographic barrier also in savanna trees (Allal et al., 2011; Lompo et al., 2018), and in general as a phytogeographic transition zone between rainforest and savanna (Droissart et al., 2018).

Despite the obvious biogeographic importance of the CVL, little attention has been devoted to the study of the phylogeography within the mountains. With limited samplings, some studies included the CVL within broader researches of the Afromontane ecoregion. Phylogeographic studies of the montane forest endemic tree *Prunus africana* uncovered a close relationship between the CVL and Albertine Rift (in western Uganda; Dawson and Powell, 1999; Kadu et al., 2011), suggesting a relatively recent historical connection between these distant mountain systems dated to the Late Pleistocene, probably to the Last Glacial Maximum (Kadu et al., 2011). Two CVL’s species of the murine rodent *Otomys* (inhabitants of grasslands and swamps) were also found to have a relationship to an East African clade, but older than the *Prunus* one, with the divergence dated to the Early Pleistocene some 2.3 to 2.0 Mya (Taylor et al., 2014). Similarly, but even an older split dated to the Pliocene (approx. 3.5 Mya) was inferred between two endemic montane forests murid genera, *Lamottemys* from Mt. Oku and *Desmomys* from the Ethiopian Highlands (Missoup et al., 2016). Within the Cameroon Volcanic line, a phylogeographic study (Smith et al., 2000) on two bird species found three main clades in both species, geographically corresponding to Bioko Island, Mt. Cameroon, and the Cameroon Highlands forests (CHf). For the latter—area of our interest— Smith et al. (2000) found a divergence in one species between the southern (Mt. Kupe) and northern (Mt. Oku, Tchabal Mbabo) populations. However, the sample size was small and thus inconclusive. In the second species, they did not find a clear phylogeographic pattern indicating that at least some flying avian species are capable to disperse across different mountain blocks of the CHf. Beside the bird study, limited in both sampling and molecular markers (mtDNA), we are not aware of any other detailed phylogeographic study of the CVL or CHf, except for Zimkus and Gvoždík (2013).

#### 4.5.2. Phylogeography of the P. steindachneri species complex

Here, we build on our previous study (Zimkus and Gvoždík, 2013), and provide the first detailed phylogeographic study of a montane endemic restricted to the CHf, supported by genome-wide data. Our model system—the *P. steindachneri* species complex—originated within the CVL (Zimkus et al., 2010; Zimkus and Gvoždík, 2013) probably during the Pliocene (Gvoždík et al., 2020). Therefore, it did not allow us to test possible biogeographic connections of CVL to East Africa as previously shown (Kadu et al., 2011; Taylor et al., 2014). Within the species complex distributed throughout the CHf (Fig. 5), the basal split corresponds to a division of populations from southern and northern mountains at the onset of the Pleistocene (approx. 2.7-2.3 Mya). The “southern mountains” span the region from Mt. Manengouba to Mt. Oku (*P. jimzimkusi, P. njiomock*). The phylogeographic break was found within the Oku Massif, with the “northern mountains” spanning from Mt. Oku to Tchabal Mbabo (*P. sp. nov., P. steindachneri*). The next split dated to 1.7-1.5 Mya separated populations from the northernmost mountains (*P. steindachneri*: Tchabal Mbabo, Gotel Mts.) and the middle mountains (*P. sp. nov.:* Mt. Mbam, Mt. Oku, Mambilla Plateau, western slopes of Gotel Mts.), with the phylogeographic break found within the Gotel Mts. Subsequently, a split dated to 1.2-0.9 Mya separated the microendemic lineage inhabiting surroundings of Lake Oku on Mt. Oku (*P. njiomock*) from the remaining populations within the southern mountains (*P. jimzimkusi)*, again with the phylogeographic break within the Oku Massif. The main splits seem to correspond to the periods of increased aridity in Africa at around 2.8, 1.7 and 1.0 Mya (DeMenocal, 1995, 2004), when montane forests probably disintegrated in the pattern we found mirrored in the genetic architecture of our model system. Intraspecific diversifications as well as interspecific introgressions (hybridizations) within the *P. steindachneri* species complex seem to be related to the Middle and Late Pleistocene climatic oscillations, which were mediated by repeated montane forests disconnections and connections (see below). The wider mtDNA sampling of *P. jimzimkusi* provided additional information regarding the phylogeography of the southern mountains (Figs. 5F, S2). Although some haplotypes or haplogroups were found on more mountains, the main splitting pattern dated to 0.5–0.1 Mya separated populations from Mt. Manengouba to Obudu Plateau against Mt. Bamboutos, Mt. Lefo and Mt. Oku.

When we compare our results to previously published studies in animals and plants, we find some parallels. For example, the evolutionary importance of Mt. Oku, where we found several phylogeographic breaks (~2.5, 1.0, and 0.4 Mya), was underlined by the intraspecific divergence of Mt. Oku *Otomys occidentalis* dated approx. 0.6 Mya (Taylor et al., 2014) or unique haplotypes of *Prunus africana* (Kadu et al., 2011). The found repeated divergences between northern Mt. Oku and southern mountains are also roughly approximated in the divergence found between two *Otomys* species from Mt. Cameroon and Mt. Oku diverged around 1.7 Mya (Taylor et al., 2014). The sister relationship between *P. sp. nov*. and *P. steindachneri* (diverged 1.7–1.5 Myr) is paralleled by murine rodents *Praomys hartwigi* from Mt. Oku and *P. obscurus* from the Gotel Mts., with the divergence dated approx. 1 Mya (Missoup et al., 2012). In the Afromontane tree *Prunus africana*, Kadu et al. (2011) uncovered haplotype sharing between the Mambilla Plateau and Mt. Danoua (located between the Gotel Mts. and Tchabal Mbabo), which are paralleled in the presence of the same *P. sp. nov*. lineage (D) in the Mambilla Plateau and Gotel Mts. As an example from the south, we can link the (sub)montane population of the otherwise lowland forest frog *Afrixalus paradorsalis* to *P. jimzimkusi*. The (sub)montane *A. p. manengubensis* from Mt. Manengouba was estimated to originate some 2–1 Mya (Charles et al., 2018), coinciding with the mountain uplift (Marzoli et al., 2000), and paralleled to the age of origin of *P. jimzimkusi*.

#### 4.5.3. Pleistocene climatic oscillations

The main intraspecific diversifications across the *P. steindachneri* species complex seem roughly to coincide with the period 0.4 Mya when the longest (50,000 years) and possibly the warmest interglacial period took place (Lisiecki and Raymo, 2005). This opens a question about timing of montane forest connections and disconnections during the Middle and Late Pleistocene climatic oscillations. Naturally, the repeated montane forest expansions and shrinkages must have happened in a complex way. For example, Dupont (2011) proposed the spread of Afromontane forests during glacial-interglacial intermediate periods, which itself implies complicated processes. Lower elevational limits of montane organisms (i.e. interpopulation connectivity) surely fluctuated in the past, mirroring climatic changes, and they must have fluctuated differently in various parts of mountains, mirroring the topography of each particular area (for an example, see Fig. 5D). However, based on our results, we hypothesize that connections (expansions, allowing gene flow) of montane forests predominately occurred during glacial periods (typically cold and dry), when lowland rainforests were retracted (Fig. 5C). During cold periods, montane forests could have descended to lower elevations, where high humidity necessary for the montane forest ecosystem could have persisted in deep valleys. Conversely, during interglacials (including the Holocene), the warmer climate could have caused the spread of lowland forests to higher elevations, and pushed montane forests even higher, separating them on different mountains.

This proposed scenario may be seen as contradicting general views on Pleistocene rainforest fragmentations, typically linked to the glacial periods (Anhuf et al., 2006; Miller and Gosling, 2014; Duminil et al., 2015; Piñeiro et al., 2017). However, these glacial-related forest fragmentations have been referred to lowland rainforests. We hypothesize that montane forests have reacted differently, as suggested also by Dupont (2011). In reality, our scenario is in concordance with earlier views more specifically focused on montane forests (Flenley, 1979; Maley, 1987; Flenley, 1998).

## 5. Conclusions

This study provides empirical data on phylogenomic reconstructions to illustrate what occurs when genetically admixed individuals (hybrids) are included in analyses. We have demonstrated that such individuals form the ladder-like pattern (LLP) in phylogenetic trees based on individual samples, and may affect both topology and branch lengths of species trees. We therefore caution against uncritical utilization of all available specimens in phylogenomic studies of recently diverging populations (young species) where gene flow may be persistent. We have also demonstrated that for such cases species networks provide a useful analytical alternative.

We further caution against usage of mtDNA as a sole or main evidence for taxonomic implications. Here, we have shown the case of puddle frogs (*Phrynobatrachus*), where a deep mitochondrial divergence is not mirrored in the nuclear genome. Combining all available evidence (genome-wide nDNA, mtDNA, morphology), we have delimited species within the Afromontane *P. steindachneri* species complex, and described a new species, *P. sp. nov*..

Phylogeography of this forest-dwelling model system endemic to the Cameroon Highlands forests (part of Cameroon Volcanic Line) provided valuable insights into the Pleistocene history of this unique and highly threatened montane ecosystem on the intersection of West and Central Africa.

## Acknowledgements

We are grateful to David C. Blackburn for providing us with the necessary details on the collected material and photographs. We than J. Rosado (Museum of Comparative Zoology, Harvard University, Cambridge), J. Moravec (National Museum, Prague), and W. Böhme and U. Bott (Zoological Research Museum Alexander Koenig, Bonn) for loaning us voucher material, and M.-O. Rödel (Natural History Museum, Berlin) for access to the *P. steindachneri* type material. We also acknowledge technicians from the Center for Anchored Phylogenomics at Florida State University, namely M. Kortyna, S. Holland, A. Bigelow and K. Birch. We further extend our thanks to A. Kodádková and R. Tropek for additional samples, T.M. Doherty-Bone for valuable discussions, and G.C. Tasse Taboue and A. Hánová for help in the field and molecular laboratory, respectively. The field research was made possible with the permissions issued by the Cameroon Ministry of Scientific Research and Innovation (MINRESI: No. 00039/MINRESI/B00/C00/C10/C12) and Ministry of Forestry and Wildlife (MINFOF: Nos. 1010/PRBS/MINFOF/SG/DFAP/SDVEF/SC and 1099/PRS/MINFOF/SG/DFAP/SDVEF/SC). This study was funded by the Czech Science Foundation (grant number 15-13415Y), Ministry of Culture of the Czech Republic (DKRVO 2019–2023/6.VII.c, National Museum, 00023272), and the Masaryk University (MUNI/A/1098/2019).

## Supplementary material

Supplementary data to this article can be found online.

