## Supplementary material for "Gene flow in phylogenomics: Sequence capture resolves species limits and biogeography of Afromontane forest endemic frogs from the Cameroon Highlands"


**CONTENT** **Page**


**Table S1.** Material examined 1

**Fig. S1.** Measurements 6

**Appendix S1.** List and definitions of morphometric variables 7

**Fig. S2.** Maximum likelihood tree based on mtDNA (*16S*) 8

**Fig. S3.** Species tree (ASTRAL-III) with individuals as terminals 9

**Fig. S4.** Morphological variation of body size and selected variables 10

**Fig. S5.** Multivariate morphometrics of body shape (PCAs) 11

**Table S2.** Principal component analyses (PCAs) and MANOVAs 12

**Table S3.** Linear discriminant analyses (LDAs) 13

**Appendix S2.** Description of holotype of *Phrynobatrachus sp. nov.* Nečas, Dolinay, Zimkus & Gvoždík 14

**Fig. S6.** Allotype, hand/foot of holotype, and type locality of *P. sp. nov.* 15

**Table S4.** Measurements of the type series of *P. sp. nov.* 16

**Appendix S3.** Morphological variation of *P. sp. nov.* 17

**Fig. S7.** Morphological variation of *P. sp. nov.* 18

**Fig. S8.** Comparison of species and hybrids of the *P. steindachneri* species complex in life 19

**Fig. S9.** Comparison of species of the *P. steindachneri* species complex in preservative 20

**Fig. S10.** Hybrids *P. njiomock* x *P. jimzimkusi*, and *P. sp. nov.* x *P. steindachneri* 21

**Table S5.** LDA classification of genetically untested individuals from the Gotel Mts. 22

**Table S6.** Descriptive morphometrics of the *P. steindachneri* species complex 23

**Table S1.** Material examined. Holotypes in bold. Abbreviations: mtDNA = mitochondrial lineage of '*P. steindachneri*'; Morph = morphometrics (only adults were statistically analyzed, juveniles and subadults were investigated only for color pattern); AHE = anchored hybrid enrichment (sequence capture); *16S* = mitochondrial 16S rRNA gene fragment newly sequenced (+) or taken from GenBank (GB). Catalogue codes (museum collections): MCZ = Museum of Comparative Zoology, Harvard University, Cambridge; MVZ = Museum of Vertebrate Zoology, University of California, Berkeley; NHM = Natural History Museum, London; NMP = National Museum in Prague; vg = Václav Gvoždík's collection; ZFMK = "Zoologisches Forschungsmuseum Alexander Koenig", Bonn; ZMB = "Museum für Naturkunde Berlin". Country codes: CM = Cameroon, NG = Nigeria.

| *Phrynobatrachus* sp. (mtDNA) | Catalogue No. | Massif (Country) | Locality | GPS (°N) | GPS (°E) | Altitude | Sex/Stage | Morph | AHE | *16S* | Haplogroup | GenBank *16S* |
|---|---|---|---|---|---|---|---|---|---|---|---|---|
| *steindachneri* (B) | ZFMK 75705 | Tchabal Mbabo (CM) | Foungoy, 5 km NEE | 7.252 | 12.060 | 2060 m | male | + | + | GB | B_TMbabo | KF020556 |
| *steindachneri* (B) | ZFMK 75706 | Tchabal Mbabo (CM) | Foungoy, 5 km NEE | 7.252 | 12.060 | 2060 m | female | + | + | GB | B_TMbabo | KF020557 |
| *steindachneri* (B) | ZFMK 75707 | Tchabal Mbabo (CM) | Foungoy, 5 km NEE | 7.252 | 12.060 | 2060 m | male | + | + | + | B_TMbabo | xxxxxxxx |
| *steindachneri* (B) | ZFMK 75708 | Tchabal Mbabo (CM) | Foungoy, 5 km NEE | 7.252 | 12.060 | 2060 m | female | + | - | - | - | - |
| *steindachneri* (B) | ZFMK 75709 | Tchabal Mbabo (CM) | Foungoy, 5 km NEE | 7.252 | 12.060 | 2060 m | female | + | - | - | - | - |
| *steindachneri* (B) | ZFMK 75710 | Tchabal Mbabo (CM) | Foungoy, 5 km NEE | 7.252 | 12.060 | 2060 m | female | + | - | - | - | - |
| *steindachneri* (B) | ZFMK 75711 | Tchabal Mbabo (CM) | Foungoy, 5 km NEE | 7.252 | 12.060 | 2060 m | female | + | - | - | - | - |
| *steindachneri* (A) | NMP-P6V 74520/1 | Gotel Mts. (CM/NG) | Mt. Gangirwal, site 2 | 7.040 | 11.707 | 2250 m | female | + | + | GB | A_Gotel | KF020551 |
| *steindachneri* (A) | NMP-P6V 74520/2 | Gotel Mts. (CM/NG) | Mt. Gangirwal, site 2 | 7.040 | 11.707 | 2250 m | juvenile | - | - | - | - | - |
| *steindachneri* (A) | NMP-P6V 74520/3 | Gotel Mts. (CM/NG) | Mt. Gangirwal, site 2 | 7.040 | 11.707 | 2250 m | subadult | - | - | - | - | - |
| *steindachneri* (A) | NMP-P6V 74520/4 | Gotel Mts. (CM/NG) | Mt. Gangirwal, site 2 | 7.040 | 11.707 | 2250 m | subadult | - | - | - | - | - |
| *steindachneri* (A) | NMP-P6V 74520/5 | Gotel Mts. (CM/NG) | Mt. Gangirwal, site 2 | 7.040 | 11.707 | 2250 m | juvenile | - | + | GB | A_Gotel | KF020552 |
| *steindachneri* (A) | NMP-P6V 74520/6 | Gotel Mts. (CM/NG) | Mt. Gangirwal, site 2 | 7.040 | 11.707 | 2250 m | juvenile | - | - | GB | A_Gotel | KF020553 |
| *steindachneri* (A) | vg09-174 | Gotel Mts. (CM/NG) | Mt. Gangirwal, site 2 | 7.040 | 11.707 | 2250 m | juvenile | - | - | + | A_Gotel | xxxxxxxx |
| *steindachneri* (A) | NMP-P6V 74522/1 | Gotel Mts. (CM/NG) | Mt. Gangirwal, site 4 | 7.042 | 11.711 | 2330 m | female | + | + | GB | A_Gotel | KF020555 |
| *steindachneri* (A) | NMP-P6V 74522/2 | Gotel Mts. (CM/NG) | Mt. Gangirwal, site 4 | 7.042 | 11.711 | 2330 m | female | + | - | - | - | - |
| *sp. nov.* (D) x *steindachneri* | NMP-P6V 74519/1 | Gotel Mts. (CM/NG) | Mt. Gangirwal, site 1 | 7.031 | 11.702 | 1940 m | male | + | - | GB | D2_Gotel | KF020547 |
| *sp. nov.* (D) x *steindachneri* | NMP-P6V 74519/2 | Gotel Mts. (CM/NG) | Mt. Gangirwal, site 1 | 7.031 | 11.702 | 1940 m | male | + | - | GB | D2_Gotel | KF020548 |
| *sp. nov.* (D) x *steindachneri* | NMP-P6V 74519/3 | Gotel Mts. (CM/NG) | Mt. Gangirwal, site 1 | 7.031 | 11.702 | 1940 m | female | + | - | GB | D3_Gotel | KF020549 |
| *sp. nov.* (D) x *steindachneri* | NMP-P6V 74519/4 | Gotel Mts. (CM/NG) | Mt. Gangirwal, site 1 | 7.031 | 11.702 | 1940 m | male | + | - | - | - | - |
| *sp. nov.* (D) x *steindachneri* | NMP-P6V 74519/5 | Gotel Mts. (CM/NG) | Mt. Gangirwal, site 1 | 7.031 | 11.702 | 1940 m | male | + | + | GB | D3_Gotel | KF020550 |
| *sp. nov.* (D) x *steindachneri* | NMP-P6V 74519/6 | Gotel Mts. (CM/NG) | Mt. Gangirwal, site 1 | 7.031 | 11.702 | 1940 m | male | + | - | - | - | - |
| *sp. nov.* (D) x *steindachneri* | NMP-P6V 74519/7 | Gotel Mts. (CM/NG) | Mt. Gangirwal, site 1 | 7.031 | 11.702 | 1940 m | male | + | - | - | - | - |
| *sp. nov.* (D) x *steindachneri* | NMP-P6V 74519/9 | Gotel Mts. (CM/NG) | Mt. Gangirwal, site 1 | 7.031 | 11.702 | 1940 m | male | + | - | - | - | - |
| *sp. nov.* (D) x *steindachneri* | NMP-P6V 74521/1 | Gotel Mts. (CM/NG) | Mt. Gangirwal, site 3 | 7.029 | 11.706 | 2010 m | female | + | + | GB | D3_Gotel | KF020554 |
| *sp. nov.* (D) x *steindachneri* | NMP-P6V 74521/2 | Gotel Mts. (CM/NG) | Mt. Gangirwal, site 3 | 7.029 | 11.706 | 2010 m | subadult | - | + | + | D2_Gotel | xxxxxxxx |
| *sp. nov.* (D) x *steindachneri* | NMP-P6V 74521/3 | Gotel Mts. (CM/NG) | Mt. Gangirwal, site 3 | 7.029 | 11.706 | 2010 m | subadult | - | - | - | - | - |
| *sp. nov.* ? *steindachneri* | ZFMK 47959 | Gotel Mts. (CM/NG) | Mt. Gangirwal | 7.03 | 11.70 | ~2300 m | male | + | - | - | - | - |
| *sp. nov.* ? *steindachneri* | ZFMK 47961 | Gotel Mts. (CM/NG) | Mt. Gangirwal | 7.03 | 11.70 | ~2300 m | male | + | - | - | - | - |
| *sp. nov.* ? *steindachneri* | ZFMK 47962 | Gotel Mts. (CM/NG) | Mt. Gangirwal | 7.03 | 11.70 | ~2300 m | male | + | - | - | - | - |
| *sp. nov.* ? *steindachneri* | ZFMK 47963 | Gotel Mts. (CM/NG) | Mt. Gangirwal | 7.03 | 11.70 | ~2300 m | male | + | - | - | - | - |
| *sp. nov.* ? *steindachneri* | ZFMK 47964 | Gotel Mts. (CM/NG) | Mt. Gangirwal | 7.03 | 11.70 | ~2300 m | female | + | - | - | - | - |
| *sp. nov.* ? *steindachneri* | ZFMK 47965 | Gotel Mts. (CM/NG) | Mt. Gangirwal | 7.03 | 11.70 | ~2300 m | male | + | - | - | - | - |
| *sp. nov.* ? *steindachneri* | ZFMK 47967 | Gotel Mts. (CM/NG) | Mt. Gangirwal | 7.03 | 11.70 | ~2300 m | male | + | - | - | - | - |
| *sp. nov.* ? *steindachneri* | ZFMK 47968 | Gotel Mts. (CM/NG) | Mt. Gangirwal | 7.03 | 11.70 | ~2300 m | male | + | - | - | - | - |

| *Phrynobatrachus* sp. (mtDNA) | Catalogue No. | Massif (Country) | Locality | GPS (°N) | GPS (°E) | Altitude | Sex/Stage | Morph | AHE | *16S* | Haplogroup | GenBank *16S* |
|---|---|---|---|---|---|---|---|---|---|---|---|---|
| *sp. nov.* ? *steindachneri* | ZFMK 47969 | Gotel Mts. (CM/NG) | Mt. Gangirwal | 7.03 | 11.70 | ~2300 m | male | + | - | - | - | - |
| *sp. nov.* ? *steindachneri* | ZFMK 47970 | Gotel Mts. (CM/NG) | Mt. Gangirwal | 7.03 | 11.70 | ~2300 m | male | + | - | - | - | - |
| *sp. nov.* ? *steindachneri* | ZFMK 47971 | Gotel Mts. (CM/NG) | Mt. Gangirwal | 7.03 | 11.70 | ~2300 m | female | + | - | - | - | - |
| *sp. nov.* ? *steindachneri* | ZFMK 47972 | Gotel Mts. (CM/NG) | Mt. Gangirwal | 7.03 | 11.70 | ~2300 m | female | + | - | - | - | - |
| *sp. nov.* ? *steindachneri* | ZFMK 47973 | Gotel Mts. (CM/NG) | Mt. Gangirwal | 7.03 | 11.70 | ~2300 m | male | + | - | - | - | - |
| *sp. nov.* ? *steindachneri* | ZFMK 47974 | Gotel Mts. (CM/NG) | Mt. Gangirwal | 7.03 | 11.70 | ~2300 m | male | + | - | - | - | - |
| *sp. nov.* ? *steindachneri* | ZFMK 47975 | Gotel Mts. (CM/NG) | Mt. Gangirwal | 7.03 | 11.70 | ~2300 m | male | + | - | - | - | - |
| *sp. nov.* ? *steindachneri* | ZFMK 47976 | Gotel Mts. (CM/NG) | Mt. Gangirwal | 7.03 | 11.70 | ~2300 m | male | + | - | - | - | - |
| *sp. nov.* ? *steindachneri* | ZFMK 47977 | Gotel Mts. (CM/NG) | Mt. Gangirwal | 7.03 | 11.70 | ~2300 m | female | + | - | - | - | - |
| *sp. nov.* ? *steindachneri* | ZFMK 47978 | Gotel Mts. (CM/NG) | Mt. Gangirwal | 7.03 | 11.70 | ~2300 m | female | + | - | - | - | - |
| *sp. nov.* ? *steindachneri* | ZFMK 47979 | Gotel Mts. (CM/NG) | Mt. Gangirwal | 7.03 | 11.70 | ~2300 m | male | + | - | - | - | - |
| *sp. nov.* ? *steindachneri* | ZFMK 47980 | Gotel Mts. (CM/NG) | Mt. Gangirwal | 7.03 | 11.70 | ~2300 m | male | + | - | - | - | - |
| *sp. nov.* ? *steindachneri* | ZFMK 47981 | Gotel Mts. (CM/NG) | Mt. Gangirwal | 7.03 | 11.70 | ~2300 m | male | + | - | - | - | - |
| *sp. nov.* ? *steindachneri* | ZFMK 47982 | Gotel Mts. (CM/NG) | Mt. Gangirwal | 7.03 | 11.70 | ~2300 m | male | + | - | - | - | - |
| *sp. nov.* ? *steindachneri* | ZFMK 47985 | Gotel Mts. (CM/NG) | Mt. Gangirwal | 7.03 | 11.70 | ~2300 m | male | + | - | - | - | - |
| *sp. nov.* ? *steindachneri* | ZFMK 47986 | Gotel Mts. (CM/NG) | Mt. Gangirwal | 7.03 | 11.70 | ~2300 m | male | + | - | - | - | - |
| *sp. nov.* (D) | MCZ A-139608 | Mambilla Plateau (NG) | Ngel Nyaki | 7.098 | 11.055 | 1530 m | female | + | + | GB | D1_Mambilla | KF020545 |
| *sp. nov.* (D) | MCZ A-139609 | Mambilla Plateau (NG) | Ngel Nyaki | 7.098 | 11.055 | 1530 m | subadult | - | - | GB | D1_Mambilla | KF020546 |
| *sp. nov.* (C) | MCZ A-138104 | Oku Massif (CM) | Mt. Oku, Elak-Oku village | 6.246 | 10.506 | 1960 m | female | + | - | GB | C2_Oku | FJ769087 |
| *sp. nov.* (C) | MCZ A-138105 | Oku Massif (CM) | Mt. Oku, Elak-Oku village | 6.246 | 10.506 | 1960 m | female | + | - | GB | C2_Oku | KF020539 |
| *sp. nov.* (C) | MCZ A-138106 | Oku Massif (CM) | Mt. Oku, Elak-Oku village | 6.246 | 10.506 | 1960 m | male | + | + | + | C2_Oku | xxxxxxxx |
| *sp. nov.* (C) | MCZ A-138107 | Oku Massif (CM) | Mt. Oku, Elak-Oku village | 6.246 | 10.506 | 1960 m | female | + | + | + | C2_Oku | xxxxxxxx |
| *sp. nov.* (C) | MCZ A-138108 | Oku Massif (CM) | Mt. Oku, Elak-Oku village | 6.246 | 10.506 | 1960 m | female | + | - | GB | C2_Oku | KF020540 |
| *sp. nov.* (C) | MCZ A-138109 | Oku Massif (CM) | Mt. Oku, Elak-Oku village | 6.246 | 10.506 | 1960 m | female | + | - | GB | C2_Oku | KF020541 |
| *sp. nov.* (C) | MCZ A-138110 | Oku Massif (CM) | Mt. Oku, Elak-Oku village | 6.246 | 10.506 | 1960 m | male | + | - | - | - | - |
| *sp. nov.* (C) | MCZ A-138111 | Oku Massif (CM) | Mt. Oku, Elak-Oku village | 6.246 | 10.506 | 1960 m | female | - | - | + | C2_Oku | xxxxxxxx |
| *sp. nov.* (C) | MCZ A-138112 | Oku Massif (CM) | Mt. Oku, Elak-Oku village | 6.246 | 10.506 | 1960 m | female | + | - | GB | C2_Oku | KF020542 |
| *sp. nov.* (C) | MCZ A-138113 | Oku Massif (CM) | Mt. Oku, Elak-Oku village | 6.246 | 10.506 | 1960 m | male | + | - | - | - | - |
| *sp. nov.* (C) | MCZ A-138116 | Oku Massif (CM) | Mt. Oku, Elak-Oku village | 6.246 | 10.506 | 1960 m | female | - | - | GB | C2_Oku | KF020543 |
| *sp. nov.* (C) | MCZ A-138117 | Oku Massif (CM) | Mt. Oku, Elak-Oku village | 6.246 | 10.506 | 1960 m | female | - | - | GB | C2_Oku | KF020544 |
| *sp. nov.* (C) | MCZ A-138056 | Mt. Mbam (CM) | Mt. Mbam, site 2 | 5.978 | 10.720 | 1970 m | male | + | - | - | - | - |
| *sp. nov.* (C) | MCZ A-138057 | Mt. Mbam (CM) | Mt. Mbam, site 1 | 5.982 | 10.720 | 2000 m | female | + | + | GB | C1_Mbam | HM441256 |
| *sp. nov.* (C) | MCZ A-138058 | Mt. Mbam (CM) | Mt. Mbam, site 1 | 5.982 | 10.720 | 2000 m | female | + | - | GB | C1_Mbam | HM441257 |
| *sp. nov.* (C) | MCZ A-138059 | Mt. Mbam (CM) | Mt. Mbam, site 1 | 5.982 | 10.720 | 2000 m | female | + | - | GB | C1_Mbam | HM441258 |
| ***sp. nov.* (C)** | **MCZ A-138060** | **Mt. Mbam (CM)** | **Mt. Mbam, site 1** | **5.982** | **10.720** | **2000 m** | **female** | + | + | + | **C1_Mbam** | **xxxxxxxx** |

| *Phrynobatrachus* sp. (mtDNA) | Catalogue No. | Massif (Country) | Locality | GPS (°N) | GPS (°E) | Altitude | Sex/Stage | Morph | AHE | *16S* | Haplogroup | GenBank *16S* |
|---|---|---|---|---|---|---|---|---|---|---|---|---|
| *njiomock* | MCZ A-136875 | Oku Massif (CM) | Mt. Oku, Lake Oku | 6.202 | 10.458 | 2230 m | female | - | - | GB | nji_Lake | FJ769095 |
| *njiomock* | MCZ A-136876 | Oku Massif (CM) | Mt. Oku, Lake Oku | 6.202 | 10.458 | 2230 m | female | - | - | GB | nji_Lake | FJ769096 |
| *njiomock* | MCZ A-136877 | Oku Massif (CM) | Mt. Oku, Lake Oku | 6.202 | 10.458 | 2230 m | female | - | - | GB | nji_Lake | FJ769097 |
| *njiomock* | MCZ A-136879 | Oku Massif (CM) | Mt. Oku, Lake Oku | 6.202 | 10.458 | 2230 m | female | - | - | GB | nji_Lake | FJ769098 |
| *njiomock* | MCZ A-138135 | Oku Massif (CM) | Mt. Oku, Lake Oku | 6.203 | 10.458 | 2230 m | female | + | - | GB | nji_Lake | FJ769099 |
| *njiomock* | MCZ A-138136 | Oku Massif (CM) | Mt. Oku, Lake Oku | 6.203 | 10.458 | 2230 m | female | + | - | GB | nji_Lake | KF020531 |
| *njiomock* | MCZ A-138137 | Oku Massif (CM) | Mt. Oku, Lake Oku | 6.203 | 10.458 | 2230 m | female | + | - | + | nji_Lake | xxxxxxxx |
| *njiomock* | MCZ A-138138 | Oku Massif (CM) | Mt. Oku, Lake Oku | 6.203 | 10.458 | 2230 m | juvenile | - | - | - | - | - |
| *njiomock* | MCZ A-138139 | Oku Massif (CM) | Mt. Oku, Lake Oku | 6.203 | 10.458 | 2230 m | female | + | - | - | - | - |
| *njiomock* | MCZ A-138140 | Oku Massif (CM) | Mt. Oku, Lake Oku | 6.203 | 10.458 | 2230 m | female | + | - | - | - | - |
| *njiomock* | NMP-P6V 73382/1 | Oku Massif (CM) | Mt. Oku, Lake Oku | 6.203 | 10.458 | 2230 m | subadult | - | - | GB | nji_Lake | KF020532 |
| *njiomock* | NMP-P6V 73382/2 | Oku Massif (CM) | Mt. Oku, Lake Oku | 6.202 | 10.458 | 2230 m | female | + | + | GB | nji_Lake | KF020533 |
| *njiomock* | NMP-P6V 73382/3 | Oku Massif (CM) | Mt. Oku, Lake Oku | 6.202 | 10.458 | 2230 m | female | + | - | + | nji_Lake | xxxxxxxx |
| *njiomock* | NMP-P6V 73382/4 | Oku Massif (CM) | Mt. Oku, Lake Oku | 6.202 | 10.458 | 2230 m | female | + | - | + | nji_Lake | xxxxxxxx |
| *njiomock* | NMP-P6V 73382/5 | Oku Massif (CM) | Mt. Oku, Lake Oku | 6.202 | 10.458 | 2230 m | male | + | - | + | nji_Lake | xxxxxxxx |
| *njiomock* | NMP-P6V 73382/6 | Oku Massif (CM) | Mt. Oku, Lake Oku | 6.202 | 10.458 | 2230 m | female | + | - | + | nji_Lake | xxxxxxxx |
| *njiomock* | NMP-P6V 74523/1 | Oku Massif (CM) | Mt. Oku, Lake Oku | 6.202 | 10.458 | 2230 m | female | - | - | GB | nji_Lake | KF020534 |
| ***njiomock*** | **NMP-P6V 74523/2** | **Oku Massif (CM)** | **Mt. Oku, Lake Oku** | **6.202** | **10.458** | **2230 m** | **male** | **-** | **-** | **GB** | **nji_Lake** | **KF020535** |
| *njiomock* | NMP-P6V 74523/3 | Oku Massif (CM) | Mt. Oku, Lake Oku | 6.202 | 10.458 | 2230 m | female | - | - | + | nji_Lake | xxxxxxxx |
| *njiomock* | NMP-P6V 74523/4 | Oku Massif (CM) | Mt. Oku, Lake Oku | 6.202 | 10.458 | 2230 m | female | - | - | + | nji_Lake | xxxxxxxx |
| *njiomock* | NMP-P6V 74523/5 | Oku Massif (CM) | Mt. Oku, Lake Oku | 6.202 | 10.458 | 2230 m | female | - | - | + | nji_Lake | xxxxxxxx |
| *njiomock* | NMP-P6V 74523/6 | Oku Massif (CM) | Mt. Oku, Lake Oku | 6.202 | 10.458 | 2230 m | female | - | - | + | nji_Lake | xxxxxxxx |
| *njiomock* | NMP-P6V 74523/x | Oku Massif (CM) | Mt. Oku, Lake Oku | 6.202 | 10.458 | 2230 m | tadpole | - | - | + | nji_Lake | xxxxxxxx |
| *njiomock* | NMP-P6V 74524 | Oku Massif (CM) | Mt. Oku, Lake Oku, 2km NW | 6.221 | 10.444 | 2400 m | subadult | - | + | GB | nji_Lake | KF020536 |
| *njiomock* x *jimzimkusi* | MCZ A-138118 | Oku Massif (CM) | Mt. Oku, forest NE of summit | 6.224 | 10.526 | 2370 m | male | + | - | GB | nji_Forest | FJ769094 |
| *njiomock* x *jimzimkusi* | MCZ A-138119 | Oku Massif (CM) | Mt. Oku, forest NE of summit | 6.224 | 10.526 | 2370 m | female | + | + | + | nji_Forest | xxxxxxxx |
| *njiomock* x *jimzimkusi* | MCZ A-138122 | Oku Massif (CM) | Mt. Oku, forest NE of summit | 6.224 | 10.526 | 2370 m | female | + | - | - | - | - |
| *jimzimkusi* | NMP-P6V 74525/1 | Oku Massif (CM) | Babungo | 6.049 | 10.426 | 1770 m | female | + | - | GB | jim_Oku | KF020525 |
| *jimzimkusi* | NMP-P6V 74525/2 | Oku Massif (CM) | Babungo | 6.049 | 10.426 | 1770 m | female | + | - | GB | jim_Oku | KF020526 |
| *jimzimkusi* | NMP-P6V 74525/3 | Oku Massif (CM) | Babungo | 6.049 | 10.426 | 1770 m | female | + | - | + | jim_Oku | xxxxxxxx |
| *jimzimkusi* | NMP-P6V 74525/4 | Oku Massif (CM) | Babungo | 6.049 | 10.426 | 1770 m | female | + | - | + | jim_Oku | xxxxxxxx |
| *jimzimkusi* | NMP-P6V 74525/5 | Oku Massif (CM) | Babungo | 6.049 | 10.426 | 1770 m | male | + | - | + | jim_Oku | xxxxxxxx |
| *jimzimkusi* | NMP-P6V 74525/6 | Oku Massif (CM) | Babungo | 6.049 | 10.426 | 1770 m | female | + | - | + | jim_Oku | xxxxxxxx |
| *jimzimkusi* | NMP-P6V 74525/7 | Oku Massif (CM) | Babungo | 6.049 | 10.426 | 1770 m | male | + | - | GB | jim_Oku | KF020527 |
| *jimzimkusi* | NMP-P6V 73390/1 | Oku Massif (CM) | Mt. Babanki, above village | 6.102 | 10.274 | 1310 m | male | + | - | GB | jim_Oku | KF020523 |
| *jimzimkusi* | NMP-P6V 73390/2 | Oku Massif (CM) | Mt. Babanki, above village | 6.102 | 10.274 | 1310 m | male | + | - | GB | jim_Oku | KF020524 |
| *jimzimkusi* | NMP-P6V 73390/3 | Oku Massif (CM) | Mt. Babanki, above village | 6.102 | 10.274 | 1310 m | female | + | - | + | jim_Oku | xxxxxxxx |
| *jimzimkusi* | NMP-P6V 73390/4 | Oku Massif (CM) | Mt. Babanki, above village | 6.102 | 10.274 | 1310 m | female | + | - | + | jim_Oku | xxxxxxxx |

| *Phrynobatrachus* sp. (mtDNA) | Catalogue No. | Massif (Country) | Locality | GPS (°N) | GPS (°E) | Altitude | Sex/Stage | Morph | AHE | *16S* | Haplogroup | GenBank *16S* |
|---|---|---|---|---|---|---|---|---|---|---|---|---|
| *jimzimkusi* | NMP-P6V 73390/5 | Oku Massif (CM) | Mt. Babanki, above village | 6.102 | 10.274 | 1310 m | female | + | - | + | jim_Oku | xxxxxxxx |
| *jimzimkusi* | NMP-P6V 73390/6 | Oku Massif (CM) | Mt. Babanki, above village | 6.102 | 10.274 | 1310 m | female | - | - | + | jim_Oku | xxxxxxxx |
| *jimzimkusi* | NMP-P6V 74528 | Oku Massif (CM) | Mt. Babanki, above village | 6.102 | 10.274 | 1310 m | male | - | - | + | jim_Oku | xxxxxxxx |
| *jimzimkusi* | NMP-P6V 73385/1 | Oku Massif (CM) | Mt. Babanki, Mendong Buo | 6.089 | 10.297 | 2100 m | male | + | - | GB | jim_Oku | KF020520 |
| *jimzimkusi* | NMP-P6V 73385/2 | Oku Massif (CM) | Mt. Babanki, Mendong Buo | 6.089 | 10.297 | 2100 m | male | + | - | GB | jim_Oku | KF020521 |
| *jimzimkusi* | NMP-P6V 73385/3 | Oku Massif (CM) | Mt. Babanki, Mendong Buo | 6.089 | 10.297 | 2100 m | female | + | + | + | jim_Oku | xxxxxxxx |
| *jimzimkusi* | NMP-P6V 73385/4 | Oku Massif (CM) | Mt. Babanki, Mendong Buo | 6.089 | 10.297 | 2100 m | female | + | - | + | jim_Oku | xxxxxxxx |
| *jimzimkusi* | NMP-P6V 73385/5 | Oku Massif (CM) | Mt. Babanki, Mendong Buo | 6.089 | 10.297 | 2100 m | male | + | - | + | jim_Oku | xxxxxxxx |
| *jimzimkusi* | NMP-P6V 73385/6 | Oku Massif (CM) | Mt. Babanki, Mendong Buo | 6.089 | 10.297 | 2100 m | female | + | - | + | jim_Oku | xxxxxxxx |
| *jimzimkusi* | NMP-P6V 73385/7 | Oku Massif (CM) | Mt. Babanki, Mendong Buo | 6.089 | 10.297 | 2100 m | male | + | - | + | jim_Oku | xxxxxxxx |
| *jimzimkusi* | NMP-P6V 73385/8 | Oku Massif (CM) | Mt. Babanki, Mendong Buo | 6.089 | 10.297 | 2100 m | male | + | - | GB | jim_Oku | KF020522 |
| *jimzimkusi* | NMP-P6V 73385/9 | Oku Massif (CM) | Mt. Babanki, Mendong Buo | 6.089 | 10.297 | 2100 m | juvenile | - | - | + | jim_Oku | xxxxxxxx |
| *jimzimkusi* | NMP-P6V 73385/10 | Oku Massif (CM) | Mt. Babanki, Mendong Buo | 6.089 | 10.297 | 2100 m | juvenile | - | - | + | jim_Oku | xxxxxxxx |
| *jimzimkusi* | NMP-P6V 73385/11 | Oku Massif (CM) | Mt. Babanki, Mendong Buo | 6.089 | 10.297 | 2100 m | juvenile | - | - | + | jim_Oku | xxxxxxxx |
| *jimzimkusi* | NMP-P6V 73385/12 | Oku Massif (CM) | Mt. Babanki, Mendong Buo | 6.089 | 10.297 | 2100 m | male | + | - | + | jim_Oku | xxxxxxxx |
| *jimzimkusi* | NMP-P6V 73385/13 | Oku Massif (CM) | Mt. Babanki, Mendong Buo | 6.089 | 10.297 | 2100 m | female | + | - | + | jim_Oku | xxxxxxxx |
| *jimzimkusi* | NMP-P6V 74527/1 | Oku Massif (CM) | Mt. Babanki, Mendong Buo | 6.088 | 10.295 | 2100 m | male | + | - | + | jim_Oku | xxxxxxxx |
| *jimzimkusi* | NMP-P6V 74527/2 | Oku Massif (CM) | Mt. Babanki, Mendong Buo | 6.088 | 10.295 | 2100 m | female | + | - | + | jim_Oku | xxxxxxxx |
| *jimzimkusi* | NMP-P6V 74527/3 | Oku Massif (CM) | Mt. Babanki, Mendong Buo | 6.088 | 10.295 | 2100 m | male | + | - | + | jim_Oku | xxxxxxxx |
| *jimzimkusi* | NMP-P6V 74527/4 | Oku Massif (CM) | Mt. Babanki, Mendong Buo | 6.088 | 10.295 | 2100 m | male | + | - | + | jim_Oku | xxxxxxxx |
| *jimzimkusi* | NMP-P6V 74527/5 | Oku Massif (CM) | Mt. Babanki, Mendong Buo | 6.088 | 10.295 | 2100 m | female | + | - | + | jim_Oku | xxxxxxxx |
| *jimzimkusi* | NMP-P6V 74527/6 | Oku Massif (CM) | Mt. Babanki, Mendong Buo | 6.088 | 10.295 | 2100 m | female | + | - | + | jim_Oku | xxxxxxxx |
| *jimzimkusi* | vg05-P-I-A | Oku Massif (CM) | Mt. Babanki, Mendong Buo | 6.089 | 10.297 | 2100 m | tadpole | - | - | + | jim_Oku | xxxxxxxx |
| *jimzimkusi* | vg05-P-I-B | Oku Massif (CM) | Mt. Babanki, Mendong Buo | 6.089 | 10.297 | 2100 m | tadpole | - | - | + | jim_Oku | xxxxxxxx |
| *jimzimkusi* | vg05-P-I-C | Oku Massif (CM) | Mt. Babanki, Mendong Buo | 6.089 | 10.297 | 2100 m | tadpole | - | - | + | jim_Oku | xxxxxxxx |

| *Phrynobatrachus* sp. (mtDNA) | Catalogue No. | Massif (Country) | Locality | GPS (°N) | GPS (°E) | Altitude | Sex/Stage | Morph | AHE | *16S* | Haplogroup | GenBank *16S* |
|---|---|---|---|---|---|---|---|---|---|---|---|---|
| *jimzimkusi* | vg05-V | Oku Massif (CM) | Mt. Babanki, Mendong Buo | 6.089 | 10.297 | 2100 m | egg | - | - | + | jim_Oku | xxxxxxxx |
| *jimzimkusi* | NMP-P6V 74526/1 | Oku Massif (CM) | Mt. Oku, Bat | 6.159 | 10.449 | 2070 m | male | + | - | GB | jim_Oku | KF020528 |
| *jimzimkusi* | NMP-P6V 74526/2 | Oku Massif (CM) | Mt. Oku, Bat | 6.159 | 10.449 | 2070 m | subadult | - | - | GB | jim_Oku | KF020529 |
| *jimzimkusi* | NMP-P6V 74526/3 | Oku Massif (CM) | Mt. Oku, Bat | 6.159 | 10.449 | 2070 m | subadult | - | - | GB | jim_Oku | KF020530 |
| *jimzimkusi* | MCZ A-139119 | Oku Massif (CM) | Mt. Oku, near summit, site 1 | 6.192 | 10.522 | 2800 m | tadpole | - | - | GB | jim_Oku | FJ769089 |
| *jimzimkusi* | NHM? (isolate A14) | Oku Massif (CM) | Mt. Oku, near summit, site 2 | 6.17 | 10.50 | 2600 m | | - | - | GB | jim_Oku | KP122353 |
| *jimzimkusi* | MCZ A-138079 | Mt. Lefo (CM) | Lake Awing | 5.864 | 10.199 | 2040 m | | - | - | - | jim_Oku* | HM441259* |
| *jimzimkusi* | MCZ A-138080 | Mt. Lefo (CM) | Lake Awing | 5.864 | 10.199 | 2040 m | | - | - | GB | jim_Bamboutos1 | HM441260 |
| *jimzimkusi* | MCZ A-138081 | Mt. Lefo (CM) | Lake Awing | 5.864 | 10.199 | 2040 m | | - | - | GB | jim_Oku | HM441261 |
| *jimzimkusi* | MCZ A-136903 | Mt. Bamboutos (CM) | below summit | 5.661 | 10.127 | 2050 m | female | + | - | GB | jim_Bamboutos1 | FJ769085 |
| ***jimzimkusi*** | **MCZ A-136904** | **Mt. Bamboutos (CM)** | **below summit** | **5.661** | **10.127** | **2050 m** | **male** | **-** | **-** | **GB** | **jim_Bamboutos1** | **FJ769086** |
| *jimzimkusi* | MCZ A-136906 | Mt. Bamboutos (CM) | below summit | 5.661 | 10.127 | 2050 m | adult | - | - | + | jim_Bamboutos1 | xxxxxxxx |
| *jimzimkusi* | MCZ A-136907 | Mt. Bamboutos (CM) | below summit | 5.661 | 10.127 | 2050 m | male | + | - | + | jim_Bamboutos1 | xxxxxxxx |
| *jimzimkusi* | MCZ A-136908 | Mt. Bamboutos (CM) | below summit | 5.661 | 10.127 | 2050 m | female | + | - | - | - | - |
| *jimzimkusi* | MCZ A-136909 | Mt. Bamboutos (CM) | below summit | 5.661 | 10.127 | 2050 m | juvenile | - | - | GB | jim_Bamboutos1 | FJ769084 |
| *jimzimkusi* | MCZ A-138064 | Mt. Bamboutos (CM) | Mt. Bamboutos | 5.623 | 10.111 | 2020 m | juvenile | - | - | GB | jim_Bamboutos1 | FJ769083 |
| *jimzimkusi* | MCZ A-138065 | Mt. Bamboutos (CM) | Mt. Bamboutos | 5.623 | 10.111 | 2020 m | female | + | - | + | jim_Oku | xxxxxxxx |
| *jimzimkusi* | MCZ A-138066 | Mt. Bamboutos (CM) | Mt. Bamboutos | 5.623 | 10.111 | 2020 m | female | + | + | + | jim_Bamboutos2 | xxxxxxxx |
| *jimzimkusi* | MCZ A-138071 | Mt. Bamboutos (CM) | Mt. Bamboutos | 5.623 | 10.111 | 2020 m | female | + | - | - | - | - |
| *jimzimkusi* | MCZ A-138072 | Mt. Bamboutos (CM) | Mt. Bamboutos | 5.661 | 10.127 | 2050 m | female | + | - | - | - | - |
| *jimzimkusi* | MCZ A-138075 | Mt. Bamboutos (CM) | summit | 5.669 | 10.103 | 2540 m | male | + | - | - | - | - |
| *jimzimkusi* | MCZ A-138076 | Mt. Bamboutos (CM) | summit | 5.669 | 10.103 | 2540 m | female | + | - | - | - | - |
| *jimzimkusi* | MCZ A-136891 | Mt. Bamboutos (CM) | summit | 5.67 | 10.10 | 2550 m | female | + | - | GB | jim_Bamboutos1 | FJ769090 |
| *jimzimkusi* | MCZ A-136892 | Mt. Bamboutos (CM) | summit | 5.67 | 10.10 | 2550 m | female | + | - | GB | jim_Manengouba | FJ769091 |
| *jimzimkusi* | MVZ 253341 | Obudu Plateau (NG) | ranger station | 6.421 | 9.360 | 1550 m | | - | - | GB | jim_Obudu | FJ769088 |
| *jimzimkusi* | MVZ 253342 | Obudu Plateau (NG) | ranger station | 6.421 | 9.360 | 1550 m | | - | - | GB | jim_Obudu | FJ769133 |
| *jimzimkusi* | MCZ A-136927 | Mt. Manengouba (CM) | Mt. Manengouba | 5.011 | 9.857 | 2050 m | female | + | - | GB | jim_Manengouba | FJ769092 |
| *jimzimkusi* | MCZ A-136928 | Mt. Manengouba (CM) | Mt. Manengouba | 5.011 | 9.857 | 2050 m | male | + | + | GB | jim_Manengouba | FJ769093 |
| *jimzimkusi* | MCZ A-136929 | Mt. Manengouba (CM) | Mt. Manengouba | 5.011 | 9.857 | 2050 m | female | + | - | - | - | - |
| *jimzimkusi* | MCZ A-136930 | Mt. Manengouba (CM) | Mt. Manengouba | 5.011 | 9.857 | 2050 m | male | + | - | - | - | - |
| *jimzimkusi* | ZMB 80238 | Mt. Manengouba (CM) | Mt. Manengouba | | | | adult | - | - | GB | jim_Manengouba | KJ626418 |
| *jimzimkusi* | ZMB 79659 | Mt. Manengouba (CM) | Abdou | 5.039 | 9.861 | 2000 m | tadpole | - | - | GB | jim_Manengouba | KJ626414 |
| *jimzimkusi* | ZMB 79662 | Mt. Manengouba (CM) | near Pola | 5.058 | 9.828 | 1720 m | tadpole | - | - | GB | jim_Manengouba | KJ626416 |
| *jimzimkusi* | ZMB 79661 | Mt. Manengouba (CM) | near summit | 5.010 | 9.857 | 2140 m | tadpole | - | - | GB | jim_Manengouba | KJ626415 |
| *cricogaster* | NMP-P6V 74642 | Bakossi Mts. (CM) | near Lake Edib, site 2 | 4.960 | 9.657 | 1280 m | male | - | + | GB | outgroup | MN158674 |
| *mbabo* | ZFMK 75726 | Tchabal Mbabo (CM) | Foungoy, 5 km NEE | 7.252 | 12.060 | 2060 m | female | - | - | GB | outgroup | MN158668 |
| *batesii* | NMP-P6V 74620 | Bakossi Mts. (CM) | Edib to Messaka | 4.964 | 9.647 | 1170 m | male | - | + | GB | outgroup | MN158671 |

*sequence not covering *16S*, haplogroup based on 12S rRNA (Gvoždík et al., 2020)

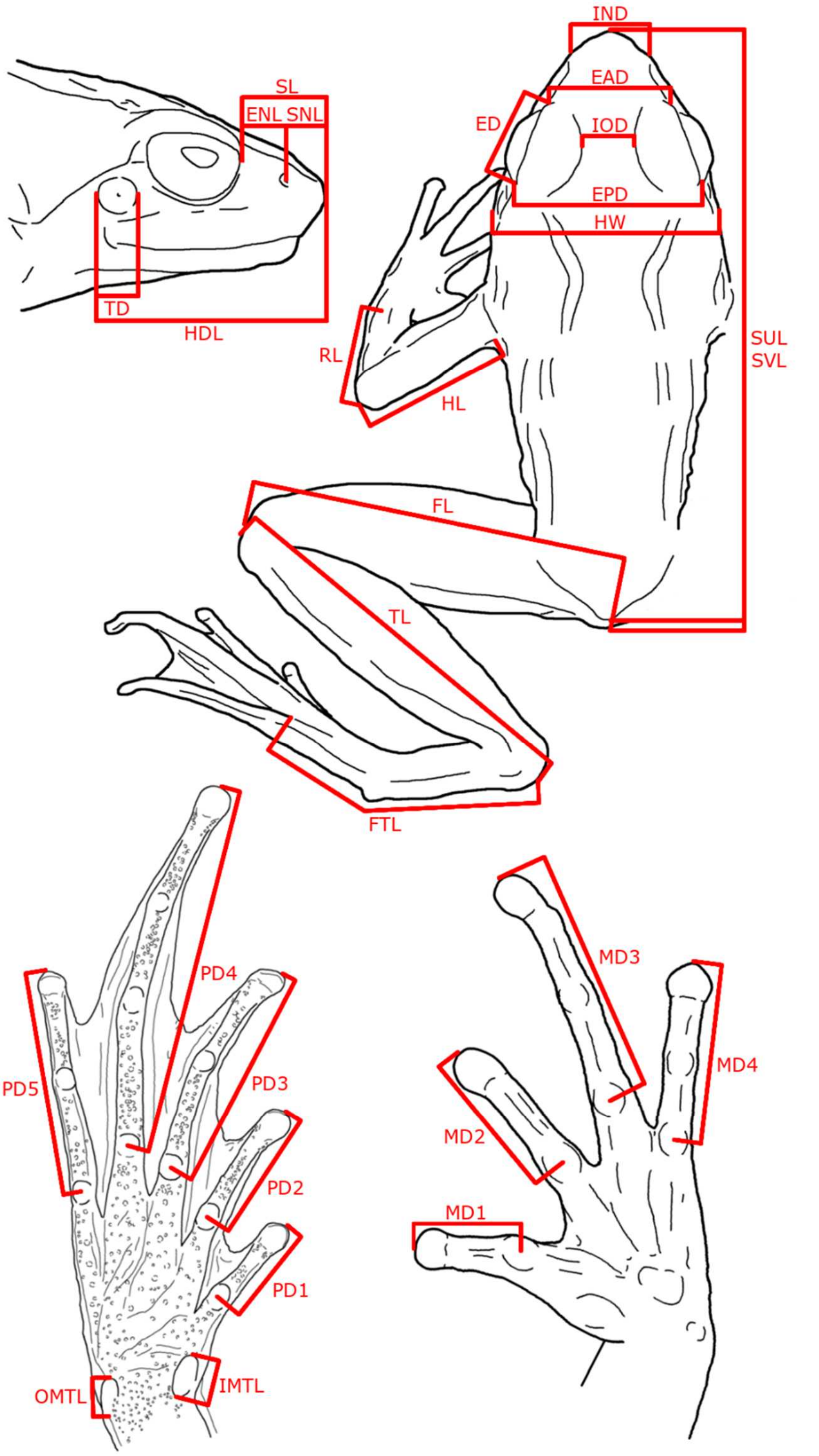


**Fig. S1.** Measurements.

**Appendix S1.** List and definitions of morphometric variables. Eighteen variables selected for multivariate morphometric analyses with explanations in bold.

**SVL,** snout-vent length, from snout tip to opening of cloaca;
**SUL, snout-urostyle length**, from snout tip to posterior edge of urostyle;
**HW, head width**, at greatest head width in close proximity to posterior edge of tympanum;
**HDL, head length**, from snout tip to posterior edge of tympanum;
**TD, tympanum diameter**, along anteroposterior body axis;
**ED, eye diameter**, between anterior and posterior corners of eye;
**IOD, interorbital distance**, shortest distance between upper eyelids at level of interorbital bar;
**EAD, eye anterior distance**, between anterior corners of eyes;
**EPD, eye posterior distance**, between posterior corners of eyes;
**SL, snout length**, from anterior eye corner to snout tip;
SNL, snout-nostril length, from snout tip to centre of nostril;
**ENL, eye-nostril length**, from anterior corner of eye to centre of nostril;
**IND, internarial distance**, between centres of nostrils;
**HL, humerus length**, from body wall to outer edge of elbow;
**RL, radioulna length**, from elbow to proximal edge of most proximal palmar tubercle;
**MD1–4, manual digit length (MD3)**, from middle of most proximal subarticular tubercle to tip of finger;
**FL, femur length**, from centre of vent to outer edge of knee;
**TL, tibiofibula length**, from outer edge of knee to outer edge of tibiotarsal articulation (heel);
**FTL, foot and tarsus length**, from outer edge of tibiotarsal articulation to most proximal subarticular tubercle of fourth pedal digit;
**PD1–5, pedal digit length (PD4)**, from middle of most proximal subarticular tubercle to tip of toe;
**IMTL**, inner metatarsal tubercle length, parallel to hind limb axis at greatest length;
**OMTL**, outer metatarsal tubercle length, parallel to hind limb axis at greatest length.

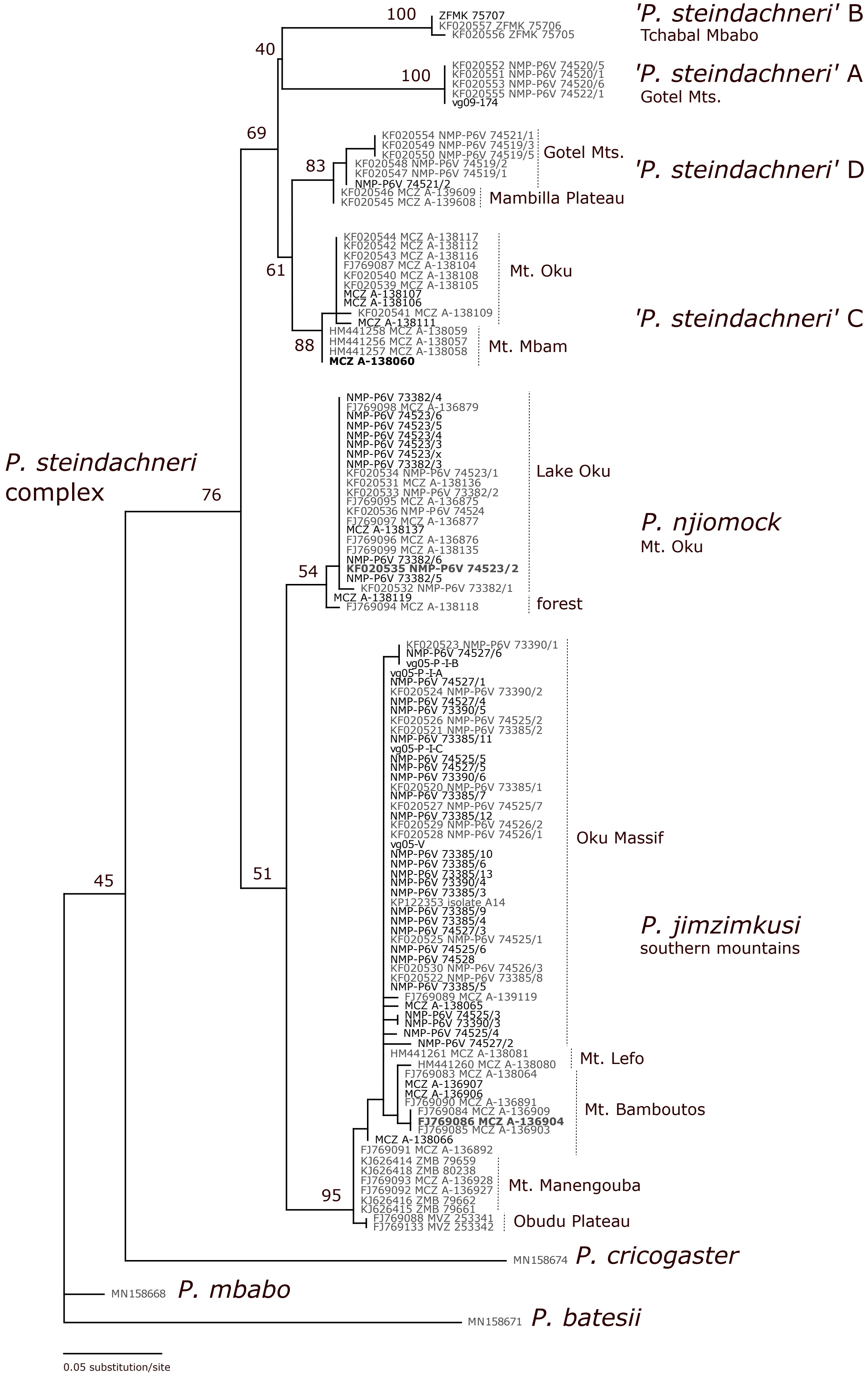


**Fig. S2.** Maximum likelihood tree based on mtDNA (*16S*). GenBank sequences in grey, holotypes in bold. Numbers at nodes are bootstrap support values.

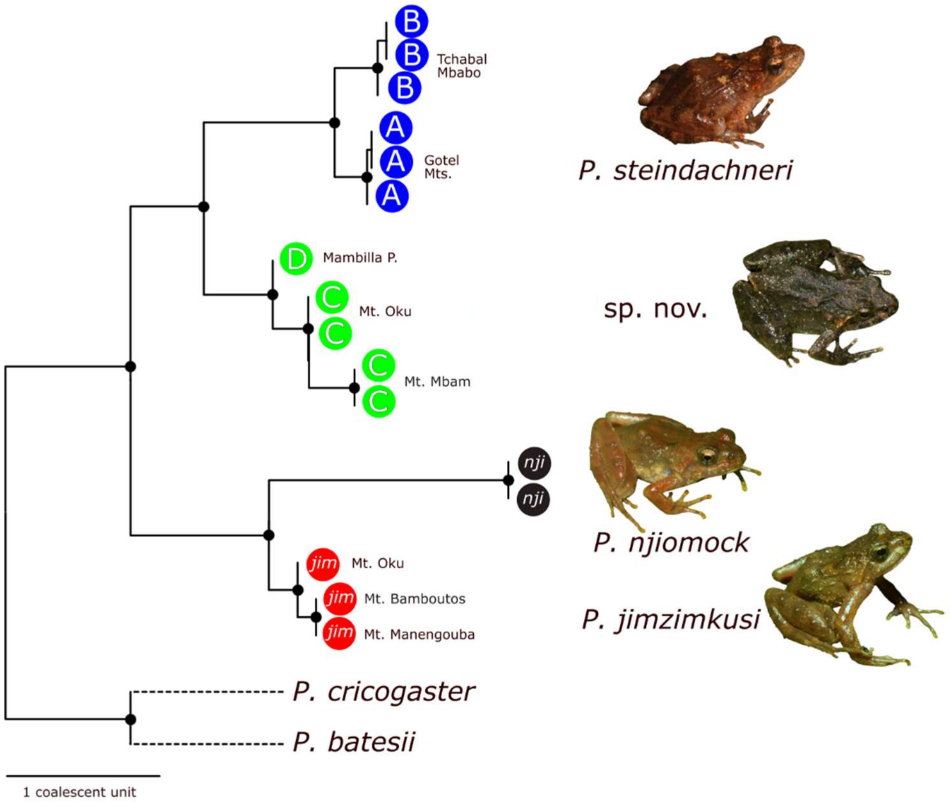


**Fig. S3.** Species tree (ASTRAL-III) with individuals as terminals (after exclusion of hybrids) based on sequence-capture genomic data.

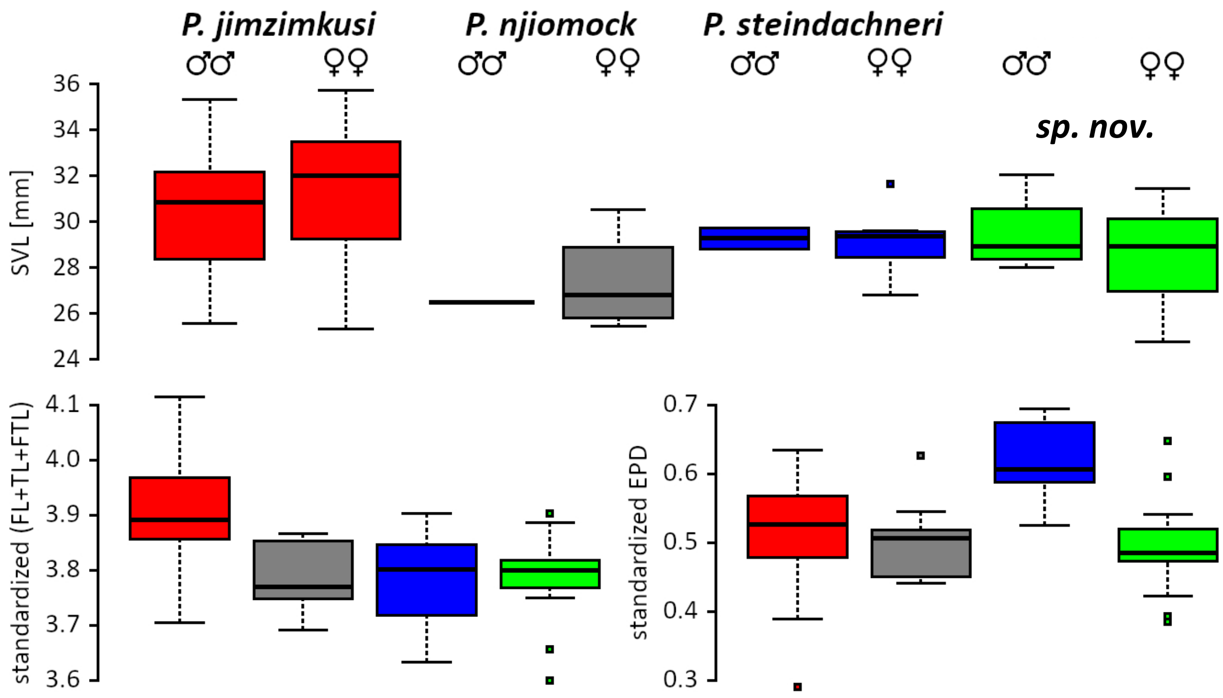


**Fig. S4.** Boxplots showing morphological variation in body size (snout-vent length; upper row) and selected variables of the four species as newly defined in this study, *P. jimzimkusi*, *P. njiomock*, *P. steindachneri* sensu stricto, and *P. sp. nov.* Hind limb length (lower row left) and head width as measured at posterior eye corners (lower row right), both after the size standardization. Horizontal lines showing average values, boxes upper and lower quartiles, whiskers minimums and maximums without outliers, and dots outliers. The four species are color-coded as throughout this study.

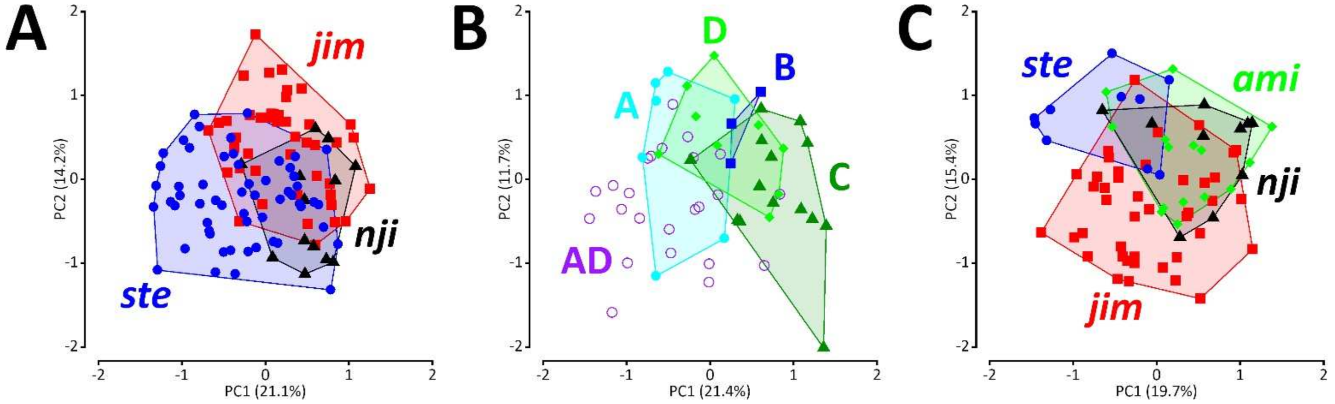


**Fig. S5.** Multivariate morphometrics showing the variation in body shape. (A) PCA 1, the starting situation corresponding to three nominal taxa, *P. jimzimkusi* (*jim*), *P. njiomock* (*nji*), and genetically diverse '*P. steindachneri*' (*ste*). (B) PCA 2, '*P. steindachneri*' showing the four lineages A–D and genetically untested individuals from the Gotel Mts. (AD). (C) PCA 3, four species as newly defined in this study: *P. sp. nov.* (*ami* = C+D), *P. jimzimkusi* (*jim*), *P. njiomock* (*nji*) and *P. steindachneri* sensu stricto (*ste* = A+B).

**Table S2.** Principal component analyses (PCAs) and Multivariate analyses of variance (MANOVAs). ** indicates $p < 0.01$, *** indicates $p \ll 0.001$.

| | Component | HW | HDL | TD | ED | IOD | EAD | EPD | SL | ENL | IND | HL | RL | MD3 | FL | TL | FTL | PD4 | MANOVA |
|---|---|---|---|---|---|---|---|---|---|---|---|---|---|---|---|---|---|---|---|
| PCA 1 | PC1 (21.1%) | -0.73 | -0.91 | 0.16 | -1.08 | -0.87 | -0.99 | -1.32 | 0.56 | 0.89 | -0.89 | -0.55 | -0.50 | 0.45 | -0.12 | 0.14 | 0.40 | 0.65 | $F_{2,6} = 14.5$ |
| | PC2 (14.2%) | 0.26 | 0.19 | -0.75 | 0.21 | -0.65 | 0.13 | 0.22 | 0.32 | 0.26 | -0.39 | 0.53 | 0.73 | -0.08 | 1.26 | 1.20 | 0.91 | 0.19 | $p = 6.1e{-}14$ |
| | PC3 (10.9%) | -0.14 | -0.65 | 0.30 | -0.18 | -0.10 | 0.17 | -0.09 | -1.33 | -1.04 | -0.03 | 0.12 | 0.20 | 0.76 | 0.30 | 0.29 | 0.04 | 0.70 | *** |
| PCA 2 | PC1 (21.4%) | -0.92 | -0.52 | -0.01 | -0.81 | -0.76 | -0.80 | -1.10 | 0.70 | 0.97 | -0.58 | -0.55 | -0.65 | 0.30 | -0.32 | -0.24 | 0.23 | 0.23 | $F_{3,29} = 4.4$ |
| | PC2 (11.7%) | 0.71 | 0.04 | -0.64 | 0.38 | -0.45 | 0.14 | 0.42 | 0.31 | 0.42 | -0.65 | -0.35 | -0.30 | 0.08 | 0.7 | 0.80 | -0.25 | 0.25 | $p = 0.0002$ |
| | PC3 (10.3%) | -0.44 | 0.61 | -0.59 | 0.27 | -0.09 | -0.32 | 0.12 | 0.81 | 0.50 | 0.38 | 0.72 | 0.25 | -0.59 | -0.10 | 0.09 | 0.10 | -0.22 | *** |
| PCA 3 | PC1 (19.7%) | -0.67 | -0.87 | 0.81 | -1.03 | -0.09 | -0.63 | -1.10 | -0.09 | 0.23 | -0.23 | -0.67 | -0.73 | 0.44 | -0.81 | -0.62 | -0.23 | 0.43 | $F_{3,9} = 7.8$ |
| | PC2 (15.4%) | 0.27 | 0.54 | 0.01 | 0.39 | 0.89 | 0.14 | 0.39 | 0.35 | 0.13 | 0.56 | -0.39 | -0.60 | -0.50 | -0.89 | -0.86 | -1.05 | -0.46 | $p = 1.3e{-}09$ |
| | PC3 (11.8%) | -0.26 | -0.25 | 0.08 | 0.39 | 0.08 | 0.16 | 0.21 | -1.24 | -1.21 | 0.51 | 0.32 | -0.14 | 0.58 | -0.18 | -0.22 | -0.23 | 0.19 | *** |
| PCA 4 | PC1 (21.4%) | -0.92 | -0.52 | -0.01 | -0.81 | -0.76 | -0.80 | -1.10 | 0.70 | 0.97 | -0.58 | -0.55 | -0.65 | 0.30 | -0.32 | -0.24 | 0.23 | 0.23 | $F_{2,31} = 3.2$ |
| | PC2 (11.7%) | 0.71 | 0.04 | -0.64 | 0.38 | -0.45 | 0.14 | 0.42 | 0.31 | 0.42 | -0.65 | -0.35 | -0.30 | 0.08 | 0.7 | 0.80 | -0.25 | 0.25 | $p = 0.008$ |
| | PC3 (10.3%) | -0.44 | 0.61 | -0.59 | 0.27 | -0.09 | -0.32 | 0.12 | 0.81 | 0.50 | 0.38 | 0.72 | 0.25 | -0.59 | -0.10 | 0.09 | 0.10 | -0.22 | ** |
| PCA 5 | PC1 (25.1%) | 0.62 | 0.66 | -0.64 | 0.64 | -0.50 | 0.71 | 0.91 | 0.27 | -0.02 | -0.22 | 0.46 | 0.77 | -0.30 | 1.04 | 0.97 | 0.84 | -0.54 | $F_{3,9} = 6.7$ |
| | PC2 (16.7%) | 0.41 | 0.70 | -0.13 | 0.21 | 0.68 | 0.05 | 0.37 | 0.97 | 0.74 | 0.37 | -0.55 | -0.33 | -0.97 | -0.38 | -0.41 | -0.35 | -0.34 | $p = 1.3e{-}07$ |
| | PC3 (10.6%) | -0.41 | -0.06 | -0.07 | -0.72 | -0.39 | -0.36 | -0.53 | 0.64 | 0.82 | -0.46 | 0.36 | 0.39 | -0.09 | 0.03 | -0.01 | 0.31 | -0.38 | *** |

**Table S3.** Linear discriminant analyses (LDAs).

| | Loadings | HW | HDL | ED | IOD | SL | IND | HL | RL | FL | TL | FTL | EAD | EPD |
|---|---|---|---|---|---|---|---|---|---|---|---|---|---|---|
| LDA 1 | LD1 (64.1%) | 6.49 | -5.77 | - | - | 0.71 | - | -5.32 | 14.91 | -5.85 | 16.37 | 10.84 | - | -6.43 |
| | LD2 (28.4%) | -5.72 | -6.61 | - | - | 1.06 | - | -1.61 | -8.77 | -6.78 | 18.75 | -0.16 | - | -7.73 |
| | LD3 (7.5%) | 8.1 | -0.53 | - | - | 1.60 | - | 5.60 | -10.10 | 0.14 | 17.10 | 8.70 | - | 3.53 |
| LDA 2 | LD1 (100%) | -13.19 | 18.72 | -0.81 | 4.09 | -5.38 | -4.63 | 3.96 | - | - | - | - | -4.95 | 18.68 |

**Appendix S2.** Description of holotype of ***Phrynobatrachus sp. nov.*** **Nečas, Dolinay, Zimkus & Gvoždík**.

MCZ A-138060, adult female, SVL = 30.1 mm (Fig. 7, Fig. S6E–F). Measurements (mm): SUL 29.7, HW 10.1, HDL 8.9, TD 1.5, ED 3.6, IOD 2.5, EAD 4.8, EPD 7.1, SL 3.5, SNL 1.5, ENL 2.0, IND 3.0, HL 5.7, RL 6.9, MD1 2.4, MD2 2.9, MD3 5.3, MD4 4.2, FL 15.2, TL 15.8, FTL 15.8, PD1 2.8, PD2 3.2, PD3 5.7, PD4 9.2, PD5 5.9, IMTL 1.1, OMTL 0.5. Elongate oval body with slender limbs; head relatively large, slightly wider (1.1x) than longer; canthus rostralis relatively long and moderately sharp; distance between nostrils slightly larger (1.2x) than between eyelids; tympanum distinguishable but rather indistinct, covered by skin, diameter less than half of eye diameter (0.4x); supratympanic fold present; dorsal chevron-shaped glands large, slightly protruding; relative length of fingers: III>IV>II>I; palmar and thenar tubercles present; fingers with round subarticular tubercles; no webbing present between manual digits; finger tips slightly bulbous; relative length of toes: IV>V>III>II>I; webbing between pedal digits well-developed (two phalanges free of webbing on fourth toe); one tarsal tubercle present; inner metatarsal tubercle length twice the outer metatarsal tubercle length (2.2x); dorsal and ventral skin relatively smooth; irregularly placed minute colorless asperities along upper jaw, shoulder, and body side, and a row along supratympanic fold extending onto eyelid.

*Coloration in preservative*: Wide brown band on dorsal side extends from snout tip to posterior edge of urostyle, margins of this stripe are ragged rather than straight and more or less follow lateral margins of dorsal glands; dorsal glands with thin white or colorless line along their medial lines, framed by darker brown; thin light dorsal stripe/line along medial body axis from snout tip to vent; tympanum covered by brown skin, part of darker but not very distinct lateral face mask, interrupted by three vertical lightly colored stripes below eye and on snout; region of jaw articulation dark brown with this color extending onto anterior side of humerus as a continuation of lateral face mask; dorsolateral portion of body light brown with sparse small brown dots; front limbs of same coloration as sides but with slight brown tint, brown stripes on radioulna, carpus, and manual digits (more pronounced on third and fourth digit); hind limbs dorsally striped with four brown stripes on femur, three on tibiofibula, two on tarsus, one on metatarsus and less pronounced on pedal digits/toes (mostly on fourth and fifth digit); background color on femur and tibiofibula is light brown; venter with brown and white marbling that extends to posterior gular region; posteromedial portion of gular region with light stripe extending to pectoral region and discontinuously in a form of white blotches across venter; white or colorless dots/asperities in pectoral region and along sides of venter, extend to body sides; background color of gular region of more yellowish tint than light venter; edge of lower jaw lined with dark brown color and adjacent series of lighter spots; posteroventral side of radioulna dark brown, extending to palm and ventral sides of phalanges; ventral side of femur lightly colored with sparse brown spots in its distal end; ventral sides of tibiofibula and tarsus also lightly colored; foot dark brown with grey tubercles; pedal webbing generally light with tiny dark pigmentation dots.

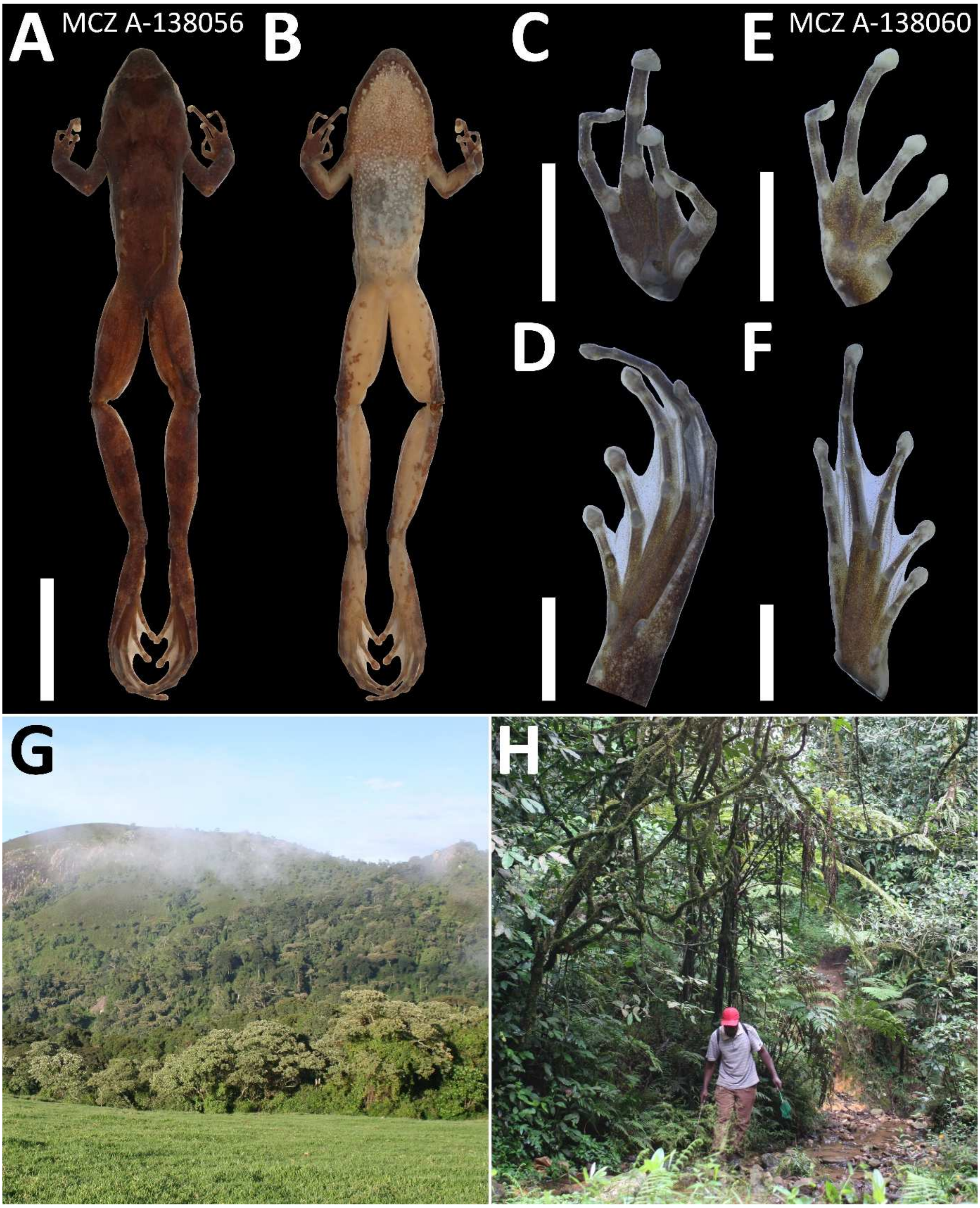


**Fig. S6.** Allotype (male paratype) of ***Phrynobatrachus sp. nov.*** (A–D); hand/foot of female holotype (E–F); and type locality (G–H) of the new species. (A) dorsal view, (B) ventral view, scale = 15 mm; (C, E) palmar and (D, F) plantar surfaces, scale = 5 mm; (G) site 1, and (H) site 2 on Mt. Mbam, Cameroon. Photos G–H by David C. Blackburn.

**Table S4.** Measurements of the type series of ***Phrynobatrachus sp. nov.*** (in mm), holotype in bold.

| Catalogue No. | MCZ A-138056 male | MCZ A-138057 female | MCZ A-138058 female | MCZ A-138059 female | **MCZ A-138060 female** |
|---|---|---|---|---|---|
| SVL | 32.1 | 28.9 | 27.8 | 28.5 | **30.1** |
| SUL | 30.7 | 28.8 | 27.7 | 28.4 | **29.7** |
| HW | 9.9 | 10.7 | 9.6 | 9.7 | **10.1** |
| HDL | 9.3 | 9.4 | 8.8 | 9.0 | **8.9** |
| TD | 1.5 | 1.7 | 1.6 | 1.5 | **1.5** |
| ED | 4.0 | 4.1 | 3.4 | 3.5 | **3.6** |
| IOD | 2.6 | 2.8 | 2.3 | 2.2 | **2.5** |
| EAD | 5.6 | 5.3 | 5.0 | 4.7 | **4.8** |
| EPD | 8.6 | 7.5 | 7.4 | 7.4 | **7.1** |
| SL | 4.2 | 3.6 | 3.7 | 3.2 | **3.5** |
| SNL | 1.4 | 1.4 | 1.2 | 1.3 | **1.5** |
| ENL | 2.8 | 2.1 | 2.5 | 2.0 | **2.0** |
| IND | 3.0 | 2.9 | 2.9 | 3.0 | **3.0** |
| HL | 6.0 | 5.8 | 5.7 | 5.8 | **5.7** |
| RL | 6.8 | 6.5 | 6.3 | 6.7 | **6.9** |
| MD1 | 2.2 | 2.0 | 2.5 | 2.4 | **2.4** |
| MD2 | 3.2 | 2.9 | 2.7 | 2.9 | **2.9** |
| MD3 | 5.5 | 4.3 | 5.1 | 5.6 | **5.3** |
| MD4 | 4.6 | 3.6 | 3.0 | 4.4 | **4.2** |
| FL | 14.4 | 16.2 | 15.5 | 16.4 | **15.2** |
| TL | 18.5 | 16.7 | 16.1 | 16.8 | **15.8** |
| FTL | 14.5 | 15.7 | 15.2 | 16.5 | **15.8** |
| PD1 | 2.9 | 2.8 | 2.6 | 2.6 | **2.8** |
| PD2 | 3.5 | 3.4 | 3.1 | 3.6 | **3.2** |
| PD3 | 6.0 | 6.0 | 5.9 | 6.1 | **5.7** |
| PD4 | 9.6 | 8.7 | 9.4 | 9.9 | **9.2** |
| PD5 | 6.2 | 6.2 | 6.0 | 6.2 | **5.9** |
| IMTL | 1.0 | 1.3 | 1.1 | 1.5 | **1.1** |
| OMTL | 0.7 | 0.6 | 0.7 | 0.7 | **0.5** |

**Appendix S3.** Morphological variation of ***Phrynobatrachus sp. nov.*** (see Figs. S7, S8A–B, S9A–B).

Snout-vent length 28.8–32.1 mm in males (n = 4), and 24.8–31.5 mm (n = 11) in females. Males have small asperities on foot and ventral tarsal surface. Descriptive morphometrics is provided in Table S6. Relatively low variation in morphological traits except of variable coloration. Elongate oval body with pointed snout, foot webbing well developed. Dorsal glands are distinct with different coloration than the rest of dorsum. The loreal region colored lighter than the rest of snout in most of specimens, making the dark lateral face mask less visible, pronounced mainly in the posterior part of head and anterior part of arm. Tympanum is distinguishable but not much distinct, having brown coloration. Interorbital band is usually well visible with lighter color positioned anteriorly. A light vertebral stripe/line of variable width might be present as well as a pair of lighter spots positioned laterally from the dorsal glands approximately in the halfway of the body length. Abdominal and pectoral coloration varies from almost white, especially in juveniles, or mostly white/light with only a few larger brown blotches to denser brown with light marbling. Coloration of gular region is generally darker than of abdominal and pectoral regions in both sexes. It varies from brown-grey background with small white dots or indistinct marbling in only posterior portion of the gular region and in angles of jaws to completely marbled throat. In majority of adults, a lighter medial stripe is indicated or well visible in the gular and pectoral regions, and disappearing in the abdominal region. Light spots that might fuse together are present along the edge of lower jaws. Slender hind limbs have dark brown bands dorsally and brown spots on light background ventrally. There is no clear differentiation in color patterns between sexes.

*Coloration in life*: Dorsal coloration varies from brown, to light brown or tan, to reddish, sometime with a yellowish vertebral stripe and/or a pair of lighter spots. Ventral surface is covered with different amounts of gray markings (both light and dark) on a creamy yellow base of variable intensity (to orangish or mostly white in some specimens), ventral surface of hindlimbs is salmony orange.

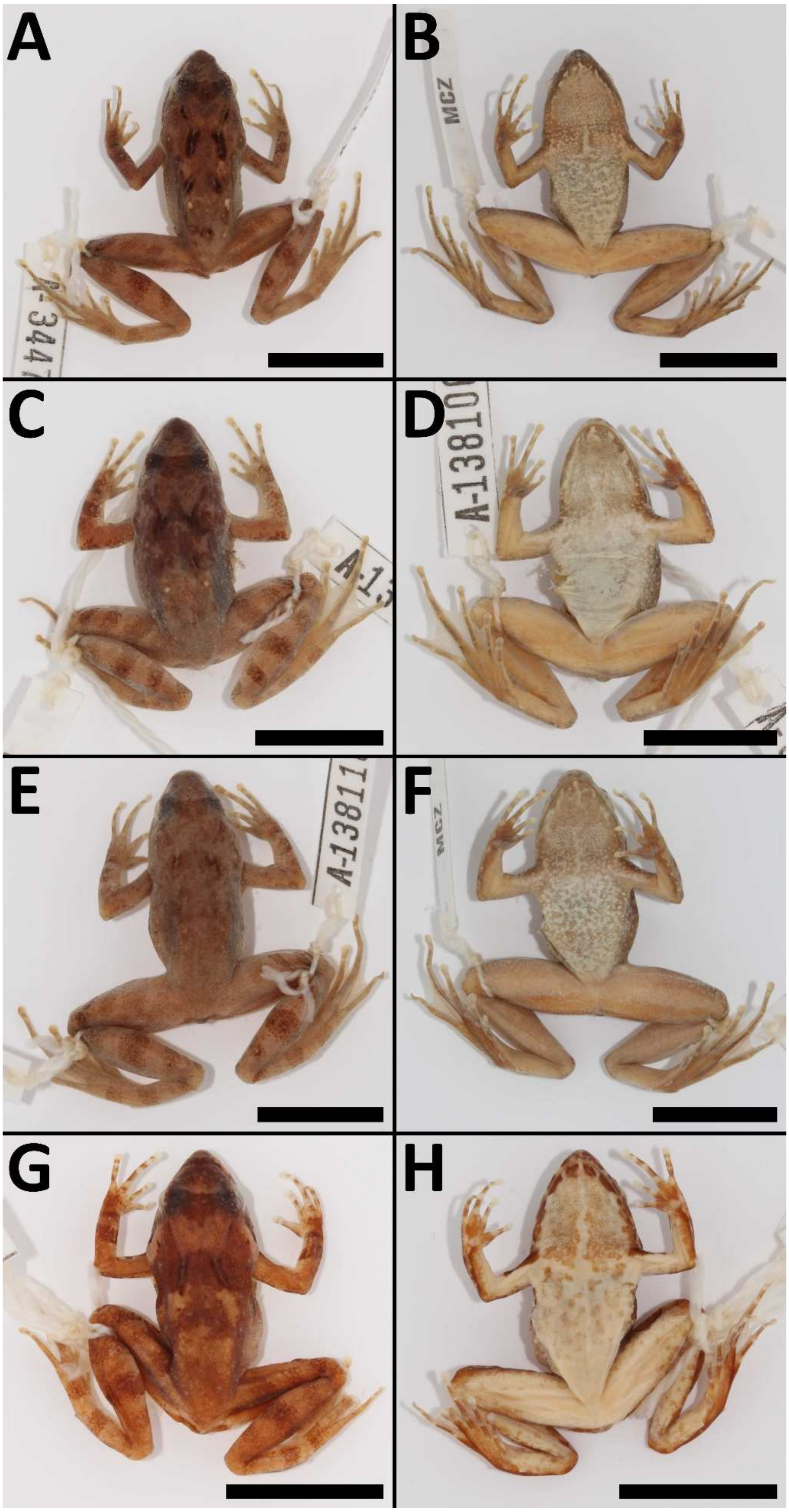


**Fig. S7.** Morphological variation of ***Phrynobatrachus sp. nov.*** (A–B) Mt. Mbam, Cameroon (MCZ A-138059, female); (C–D) Mt. Oku, Cameroon (MCZ A-138106, female); (E–F) Mt. Oku, Cameroon (MCZ A-138110, male); and (G–H) Mambilla Plateau, Nigeria (MCZ A-1389608, female); scale = 15 mm.

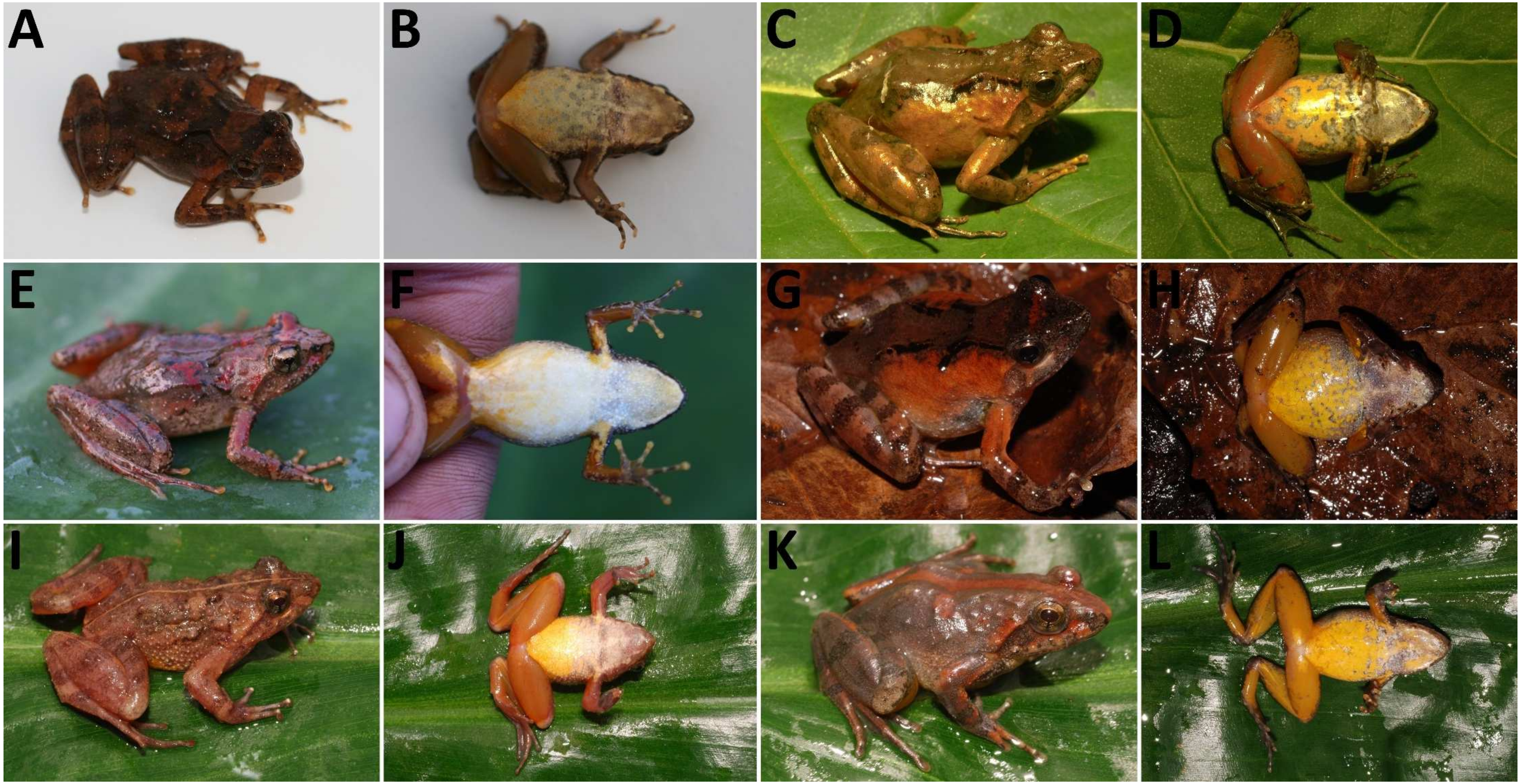


**Fig. S8.** Comparison of species and presumed hybrids of the *P. steindachneri* species complex in life. (A–B) *P. sp. nov.*, Mambilla Plateau, Nigeria (MCZ A-139608, female); (C–D) *P. jimzimkusi*, Oku Massif, Mt. Babanki, Cameroon (NMP-P6V 73385/3, female); (E–F) *P. njiomock*, Lake Oku on Mt. Oku, Cameroon (MCZ A-138139, female); (G–H) *P. steindachneri* sensu stricto, Gotel Mts., Cameroon–Nigeria (NMP-P6V 74522/1, female); and (I–L) specimens from a site, where *P. sp. nov.* x *P. steindachneri* hybrids were identified, Gotel Mts., Cameroon–Nigeria (I–J: NMP-P6V 74519/2, male, photo I horizontally flipped; K–L: NMP-P6V 74519/3, female). Note the overall morphological variation within the species complex. All color patterns can be found in all species, except for reddish blotches characteristic for *P. njiomock*. Photos by Václav Gvoždík, and David C. Blackburn (*P. sp. nov.*, *P. njiomock*).

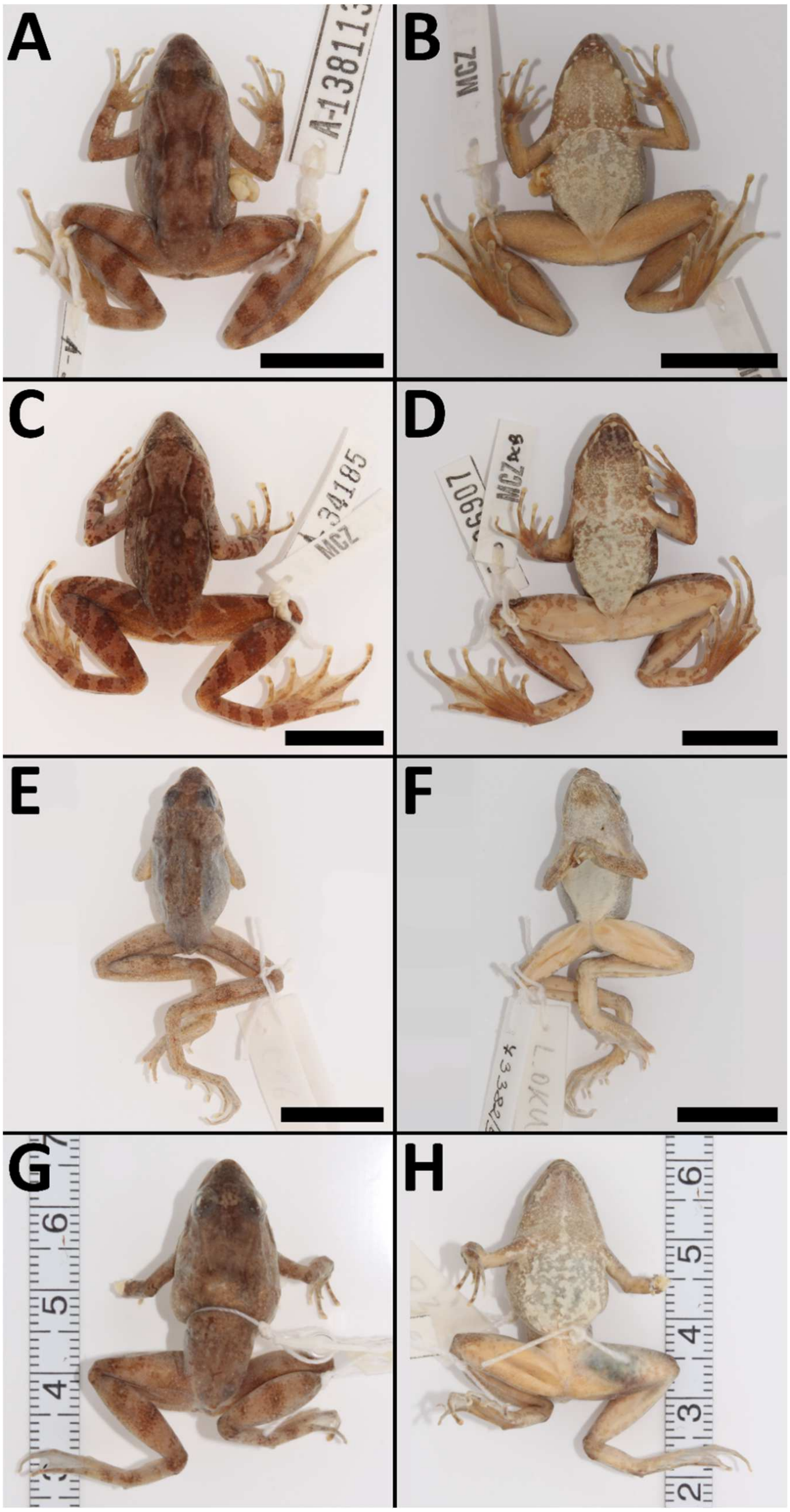


**Fig. S9.** Comparison of species of the *P. steindachneri* species complex (males) in preservative, all individuals from Cameroon. (A–B) *P. sp. nov.*, Mt. Oku (MCZ A-138113); (C–D) *P. jimzimkusi*, Mt. Bamboutos (MCZ A-136907); (E–F) *P. njiomock*, Lake Oku on Mt. Oku (NMP-P6V 73382/5); and (G–H) *P. steindachneri* sensu stricto, Tchabal Mbabo (ZFMK 75705); scale = 15 mm.

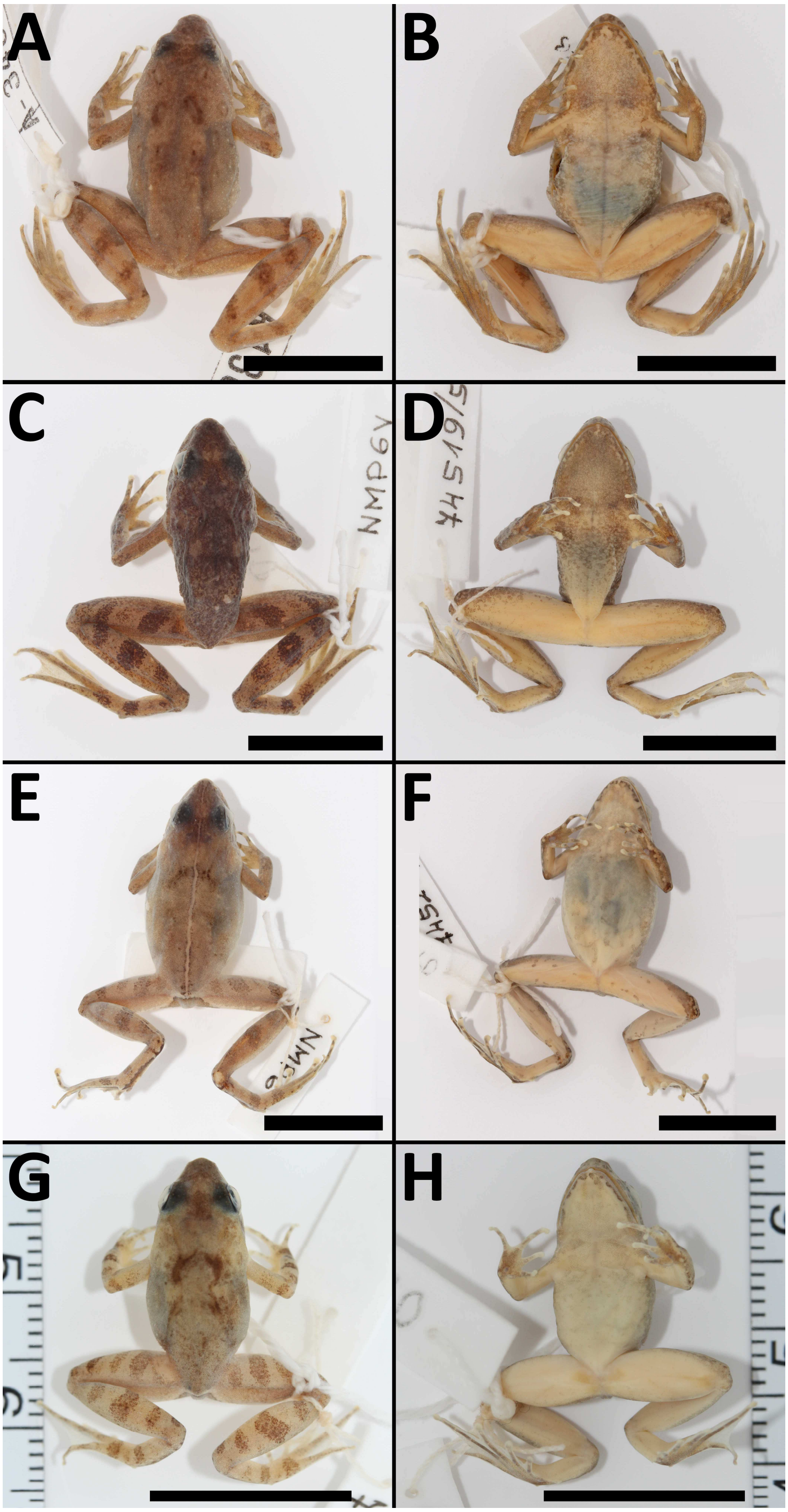

**Fig. S10.** Hybrids as evidenced by the genomic analyses. (A–B) *P. njiomock* x *P. jimzimkusi*, Mt. Oku, Cameroon (MCZ A-138119, female); and (C–H) *P. sp. nov.* x *P. steindachneri*, Gotel Mts., Cameroon–Nigeria (C–D: NMP-P6V 74519/5, male; E–F: NMP-P6V 74521/1, female; G–H: NMP6V 74521/2, subadult); scale = 15 mm.

**Table S5.** LDA classification of genetically untested individuals from the Gotel Mountains (LDA 2). Classifications to a species with probability above 95% in bold, otherwise both columns/species in bold – which may indicate hybrid individuals.

| Catalogue No. | *P. sp. nov.* [%] | *P. steindachneri* s.s. [%] |
|---|---|---|
| ZFMK 47959 | **82.3** | **17.7** |
| ZFMK 47961 | 0.1 | **99.9** |
| ZFMK 47962 | **49.2** | **50.8** |
| ZFMK 47963 | **89.0** | **11.0** |
| ZFMK 47964 | **8.6** | **91.4** |
| ZFMK 47965 | **99.7** | 0.3 |
| ZFMK 47967 | 2.2 | **97.8** |
| ZFMK 47968 | **98.9** | 1.1 |
| ZFMK 47969 | 2.2 | **97.8** |
| ZFMK 47970 | **45.5** | **54.5** |
| ZFMK 47971 | 0.0 | **100.0** |
| ZFMK 47972 | **49.5** | **50.5** |
| ZFMK 47973 | 0.0 | **100.0** |
| ZFMK 47974 | 0.0 | **100.0** |
| ZFMK 47975 | 1.3 | **98.7** |
| ZFMK 47976 | 0.0 | **100.0** |
| ZFMK 47977 | 0.8 | **99.2** |
| ZFMK 47978 | **7.8** | **92.2** |
| ZFMK 47979 | 0.0 | **100.0** |
| ZFMK 47980 | 0.0 | **100.0** |
| ZFMK 47981 | **24.9** | **75.1** |
| ZFMK 47982 | 0.2 | **99.8** |
| ZFMK 47985 | 0.5 | **99.5** |
| ZFMK 47986 | 0.0 | **100.0** |

**Table S6.** Descriptive morphometrics of the *P. steindachneri* species complex, SD = standard deviation.

| | *P. sp. nov.* Males (n = 4) | | | | *P. sp. nov.* Females (n = 11) | | | | *P. jimzimkusi* Males (n = 18) | | | | *P. jimzimkusi* Females (n = 26) | | | |
|---|---|---|---|---|---|---|---|---|---|---|---|---|---|---|---|---|
| [mm] | Mean | SD | Min | Max | Mean | SD | Min | Max | Mean | SD | Min | Max | Mean | SD | Min | Max |
| SVL | 29.5 | 1.5 | 28.0 | 32.1 | 28.5 | 2.1 | 24.8 | 31.5 | 30.6 | 2.7 | 25.5 | 35.3 | 31.4 | 2.9 | 25.3 | 35.7 |
| SUL | 28.9 | 1.0 | 28.1 | 30.7 | 28.5 | 1.9 | 24.9 | 31.5 | 30.6 | 2.7 | 25.3 | 35.3 | 31.2 | 3.0 | 24.6 | 36.0 |
| HW | 9.7 | 0.3 | 9.2 | 9.9 | 9.9 | 0.5 | 8.8 | 10.8 | 10.7 | 0.9 | 9.0 | 12.2 | 10.5 | 1.1 | 8.1 | 13.0 |
| HDL | 8.9 | 0.4 | 8.4 | 9.4 | 8.8 | 0.7 | 7.2 | 9.8 | 9.4 | 0.9 | 7.5 | 10.7 | 9.3 | 0.9 | 7.6 | 11.0 |
| TD | 1.6 | 0.2 | 1.5 | 1.9 | 1.5 | 0.2 | 1.2 | 1.7 | 1.5 | 0.3 | 1.1 | 2.0 | 1.5 | 0.3 | 1.1 | 2.1 |
| ED | 3.4 | 0.3 | 3.2 | 4.0 | 3.4 | 0.3 | 2.7 | 4.1 | 3.8 | 0.4 | 3.1 | 4.7 | 3.8 | 0.4 | 3.2 | 4.6 |
| IOD | 2.5 | 0.1 | 2.3 | 2.7 | 2.3 | 0.3 | 1.7 | 2.8 | 2.4 | 0.4 | 1.8 | 3.4 | 2.4 | 0.4 | 1.7 | 3.3 |
| EAD | 5.3 | 0.3 | 4.9 | 5.6 | 5.1 | 0.4 | 4.2 | 5.8 | 5.4 | 0.5 | 4.6 | 6.4 | 5.4 | 0.5 | 4.3 | 6.5 |
| EPD | 7.6 | 0.6 | 7.2 | 8.6 | 7.4 | 0.5 | 6.6 | 8.1 | 8.1 | 0.7 | 6.9 | 9.5 | 8.0 | 0.8 | 6.7 | 9.4 |
| SL | 3.9 | 0.3 | 3.5 | 4.2 | 3.6 | 0.4 | 3.0 | 4.3 | 3.9 | 0.4 | 3.1 | 4.5 | 3.9 | 0.4 | 3.2 | 4.9 |
| ENL | 2.6 | 0.2 | 2.3 | 2.8 | 2.2 | 0.3 | 1.9 | 2.7 | 2.3 | 0.3 | 1.7 | 2.8 | 2.3 | 0.4 | 1.7 | 3.2 |
| IND | 3.2 | 0.3 | 2.8 | 3.6 | 3.0 | 0.3 | 2.5 | 3.5 | 3.1 | 0.3 | 2.7 | 3.5 | 3.1 | 0.2 | 2.4 | 3.4 |
| HL | 6.4 | 0.4 | 6.0 | 7.0 | 5.9 | 0.6 | 5.0 | 7.0 | 7.0 | 0.9 | 5.5 | 8.6 | 6.3 | 0.7 | 4.9 | 7.6 |
| RL | 6.8 | 0.2 | 6.5 | 7.1 | 6.4 | 0.5 | 5.1 | 7.1 | 7.4 | 0.6 | 6.3 | 8.6 | 6.9 | 0.7 | 5.4 | 8.3 |
| MD3 | 5.2 | 0.4 | 4.7 | 5.5 | 5.1 | 0.6 | 4.2 | 5.9 | 5.6 | 0.9 | 3.8 | 7.2 | 5.5 | 0.6 | 4.0 | 6.3 |
| FL | 15.3 | 0.6 | 14.4 | 16.0 | 15.5 | 1.1 | 13.2 | 17.2 | 17.2 | 1.5 | 14.5 | 19.6 | 16.8 | 1.7 | 13.0 | 18.9 |
| TL | 17.5 | 0.6 | 17.0 | 18.5 | 16.5 | 1.3 | 14.4 | 18.3 | 18.6 | 1.7 | 15.4 | 21.5 | 18.2 | 1.7 | 14.9 | 20.9 |
| FTL | 15.9 | 0.9 | 14.5 | 16.9 | 15.8 | 1.4 | 13.6 | 17.6 | 17.9 | 1.7 | 14.5 | 21.0 | 17.3 | 1.7 | 14.4 | 19.9 |
| PD4 | 10.3 | 0.5 | 9.6 | 11.0 | 9.8 | 1.1 | 7.9 | 11.8 | 11.0 | 1.3 | 9.0 | 13.6 | 10.5 | 1.4 | 7.1 | 13.0 |

| | *P. njiomock* Males (n = 1) | | | | *P. njiomock* Females (n = 9) | | | | *P. steindachneri* s.s. Males (n = 2) | | | | *P. steindachneri* s.s. Females (n = 8) | | | |
|---|---|---|---|---|---|---|---|---|---|---|---|---|---|---|---|---|
| [mm] | Mean | SD | Min | Max | Mean | SD | Min | Max | Mean | SD | Min | Max | Mean | SD | Min | Max |
| SVL | 26.5 | - | - | - | 27.5 | 1.9 | 25.4 | 30.5 | 29.3 | 0.5 | 28.8 | 29.7 | 29.1 | 1.3 | 26.8 | 31.6 |
| SUL | 26.3 | - | - | - | 27.2 | 1.7 | 25.4 | 30.0 | 28.5 | 0.5 | 28.0 | 29.0 | 29.1 | 1.3 | 26.9 | 31.8 |
| HW | 9.4 | - | - | - | 8.9 | 0.6 | 8.1 | 9.9 | 9.3 | 0.3 | 9.0 | 9.6 | 9.7 | 0.5 | 8.9 | 10.5 |
| HDL | 8.7 | - | - | - | 8.0 | 0.4 | 7.4 | 8.7 | 8.7 | 0.2 | 8.5 | 8.9 | 8.8 | 0.4 | 8.1 | 9.3 |
| TD | 1.8 | - | - | - | 1.4 | 0.3 | 0.9 | 1.9 | 1.1 | 0.0 | 1.1 | 1.1 | 1.3 | 0.3 | 0.9 | 1.6 |
| ED | 3.6 | - | - | - | 3.4 | 0.2 | 3.1 | 3.7 | 3.9 | 0.1 | 3.8 | 4.0 | 3.8 | 0.4 | 3.2 | 4.3 |
| IOD | 2.5 | - | - | - | 2.3 | 0.2 | 1.9 | 2.7 | 2.4 | 0.1 | 2.2 | 2.5 | 2.4 | 0.2 | 2.0 | 2.7 |
| EAD | 5.1 | - | - | - | 4.5 | 0.4 | 3.8 | 5.1 | 5.2 | 0.0 | 5.2 | 5.2 | 4.9 | 0.4 | 4.2 | 5.4 |
| EPD | 7.7 | - | - | - | 6.8 | 0.3 | 6.3 | 7.2 | 8.1 | 0.4 | 7.6 | 8.5 | 7.9 | 0.5 | 7.0 | 8.6 |
| SL | 3.7 | - | - | - | 3.4 | 0.2 | 3.0 | 3.7 | 3.6 | 0.1 | 3.5 | 3.7 | 3.4 | 0.2 | 3.2 | 3.7 |
| ENL | 2.3 | - | - | - | 2.1 | 0.2 | 1.7 | 2.4 | 2.2 | 0.1 | 2.0 | 2.3 | 2.0 | 0.2 | 1.6 | 2.3 |
| IND | 3.3 | - | - | - | 2.7 | 0.1 | 2.5 | 3.0 | 3.0 | 0.0 | 2.9 | 3.0 | 2.9 | 0.3 | 2.5 | 3.3 |
| HL | 6.4 | - | - | - | 5.5 | 0.4 | 4.8 | 6.1 | 5.8 | 0.4 | 5.4 | 6.1 | 6.2 | 0.4 | 5.5 | 6.8 |
| RL | 6.0 | - | - | - | 5.4 | 0.4 | 4.6 | 6.1 | 5.6 | 0.4 | 5.2 | 6.1 | 6.2 | 0.4 | 5.7 | 6.8 |
| MD3 | 5.4 | - | - | - | 4.8 | 0.5 | 3.8 | 5.5 | 4.7 | 0.0 | 4.7 | 4.8 | 4.9 | 0.7 | 3.7 | 5.9 |
| FL | 15.0 | - | - | - | 14.1 | 0.9 | 12.5 | 15.6 | 14.8 | 0.4 | 14.4 | 15.2 | 14.9 | 0.8 | 13.3 | 16.1 |
| TL | 17.1 | - | - | - | 15.6 | 1.0 | 13.8 | 17.2 | 15.8 | 0.2 | 15.6 | 16.0 | 15.8 | 0.6 | 14.8 | 16.9 |
| FTL | 15.8 | - | - | - | 14.4 | 0.9 | 13.0 | 16.1 | 14.4 | 0.3 | 14.1 | 14.6 | 14.8 | 0.8 | 13.3 | 16.1 |
| PD4 | 10.7 | - | - | - | 9.8 | 0.9 | 8.5 | 11.2 | 9.6 | 0.2 | 9.4 | 9.8 | 9.0 | 0.5 | 8.5 | 10.0 |
